# Intravesical BCG in patients with non-muscle invasive bladder cancer induces trained immunity and decreases respiratory infections

**DOI:** 10.1101/2022.02.21.480081

**Authors:** Jelmer H. van Puffelen, Boris Novakovic, Liesbeth van Emst, Denise Kooper, Tahlita C.M. Zuiverloon, Ursula T.H. Oldenhof, J. Alfred Witjes, Tessel E. Galesloot, Alina Vrieling, Katja K.H. Aben, Lambertus A.L.M. Kiemeney, Egbert Oosterwijk, Mihai G. Netea, Joost L. Boormans, Antoine G. van der Heijden, Leo A.B. Joosten, Sita H. Vermeulen

## Abstract

Bacillus Calmette-Guérin (BCG) is recommended as intravesical immunotherapy to reduce the risk of tumor recurrence in patients with non-muscle invasive bladder cancer (NMIBC). Currently, it is unknown whether intravesical BCG application induces trained immunity. Here, we found that intravesical BCG does induce trained immunity based on an increased production of TNF and IL-1β after heterologous *ex-vivo* stimulation of circulating monocytes 6- 12 weeks after intravesical BCG treatment; and a 37% decreased risk (OR 0.63 (95% CI 0.40- 1.01)) for respiratory infections in BCG-treated versus non-BCG-treated NMIBC patients. An epigenomics approach combining ChIP-sequencing and RNA-sequencing with *in-vitro* trained immunity experiments identified enhanced inflammasome activity in BCG-treated individuals. Finally, germline variation in genes that affect trained immunity was associated with recurrence and progression after BCG therapy in NMIBC, suggesting a link between trained immunity and oncological outcome.

## Introduction

Intravesical instillations with *Bacillus Calmette-Guérin* (BCG) is the recommended adjuvant therapy in patients with high-risk non-muscle invasive bladder cancer (HR-NMIBC). HR-NMIBC is associated with a high risk of tumor recurrence and progression to muscle-invasive bladder cancer. BCG immunotherapy generally consists of an induction course with 6 weekly instillations, followed by 3-weekly maintenance courses at months 3 and 6 and then every 6 months for up to 3 years^1^. Although BCG is more effective in preventing tumor recurrences than intravesical chemotherapy^2, 3^, non-responsiveness to BCG is observed in more than 25% of HR-NMIBC patients within 5 years^4, 5^. The exact immunological mechanisms through which BCG mediates anti-tumor immunity in bladder cancer are still unclear. Initiation of a T helper 1 (Th1) cell immune response is important to achieve a good clinical response^6, 7^, and cell-mediated anti-tumor activity is achieved via immune cells such as CD8+ T cells^8, 9^, NK cells^10, 11^ and macrophages^12–14^.

It is known that activated myeloid cells can further increase Th1 immune responses by producing TNF, IL-1β, IL-6, IL-12 and IL-18^15–19^. These cytokines are produced after stimulation of Toll-like receptors (TLR) and NOD-like receptors (NLR) by BCG^20–22^. TLR2 and TLR4 are especially important for the production of IL-1β by monocytes after BCG stimulation^23^. Additionally, BCG vaccination and mechanistic studies have demonstrated that following BCG stimulation, myeloid cells can develop trained immunity (TI)^24^, which is characterized by long-term functional and epigenetic reprogramming of myeloid cells, with enhancement of their function^25^.This TI phenotype allows myeloid cells to respond with an increased cytokine response upon encountering a secondary stimulus or challenge^24, 25^. It contributes to the increased protection against non-tuberculosis infections, including respiratory infections, after BCG vaccination^26–28^. Mechanistic studies found that autophagy^29^, IL-1β signaling^26, 30, 31^, IL-32 signaling^32, 33^, NOD2-RIPK2 signaling^34^, epigenetic enzymes^35, 36^, metabolic reprogramming^37^, and reprogramming of hematopoietic stem and progenitor cells (HSPCs) in the bone marrow^38, 39^, are important for the induction of TI by BCG.

Whether repeated BCG instillations in NMIBC also induce TI, and whether this is relevant for clinical responses, has not been systematically studied. There are no data on epigenetic modifications and metabolic changes in myeloid cells of NMIBC patients during BCG therapy. Reports of increased urinary TNF, IL-1β, IL-2, IL-6, IL-8, IFNγ, M-CSF and GM-CSF after BCG instillations only provide indirect evidence for (systemic) TI^6, 40–43^. *Ex-vivo* LPS stimulation experiments with blood monocytes found that TNF, IL-1β and IL-6 production was generally increased during a BCG induction cycle^29, 44–46^, indicating induction of TI, but these experiments lacked relevant long-term data. Here, we extensively investigate and describe induction of TI by BCG instillations in NMIBC by i) analyzing long-term induction of systemic TI in blood monocytes of BCG-treated NMIBC patients, ii) evaluating whether BCG instillations result in reduced incidence of respiratory infections, and iii) exploring the relation between BCG-induced TI and oncological NMIBC outcome via genetic studies.

## Results

### BCG instillations induce long-term systemic trained immunity responses as indicated by increased cytokine production before BCG maintenance cycles

To determine whether BCG instillations induced TI in peripheral blood mononuclear cells (PBMCs) we performed a prospective cohort study (‘Tribute’) and isolated PBMCs collected before BCG therapy and at 8 time points during BCG therapy (Fig. 1a). A total of 17 BCG-naïve HR-NMIBC patients were included (see Supplementary Table 1 for inclusion and exclusion criteria), with a median age at inclusion of 67 years and 16 male patients (94%). Twelve patients received a complete induction regimen of 6 cycles of full-dose BCG. Four patients discontinued BCG therapy due to side effects, two patients due to disease progression, one patient because of BCG shortage, and for two patients the last BCG maintenance regimen was not performed (see Supplementary Table 2). Six patients received 1/3 dose BCG or a reduced number of BCG instillations (or both) due to BCG shortage.

**Fig. 1.**
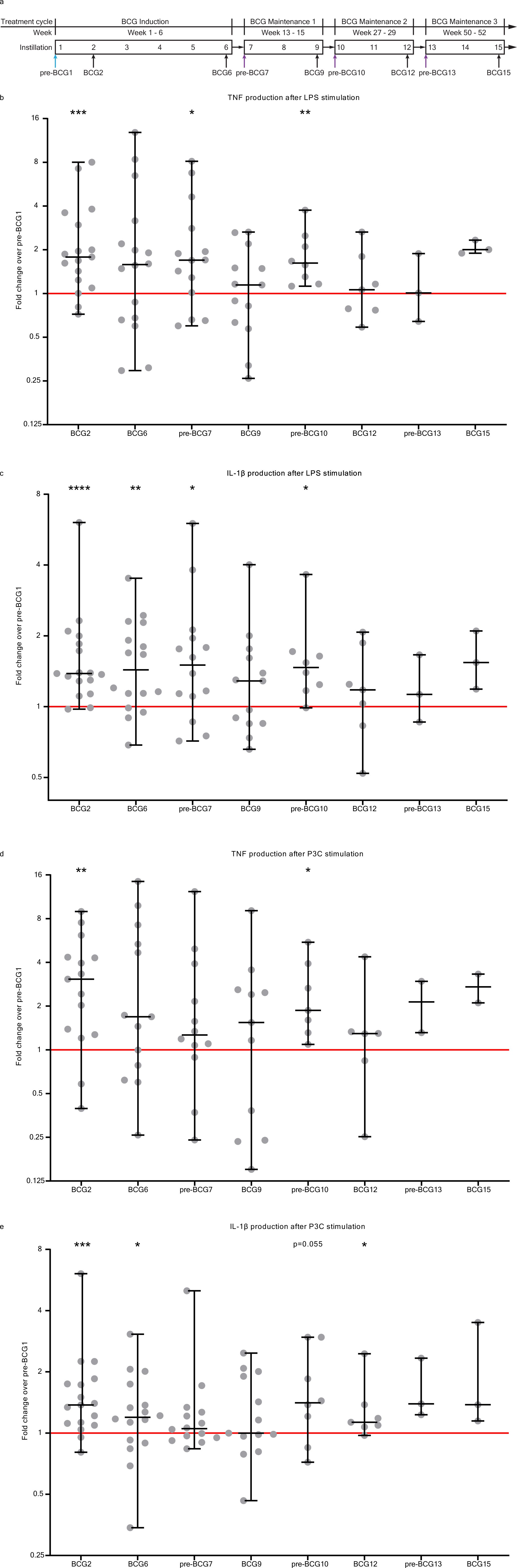
Cytokine production by PBMCs upon *ex-vivo* stimulation with heterologous stimuli is increased after BCG instillations. **a:** Study schedule of the Tribute study. Blood was collected and PBMCs were isolated at three time points during the BCG induction cycle: pre-BCG1, BCG2 and BCG6; and two time points during each subsequent BCG maintenance cycle: pre-BCG7, BCG9, pre-BCG10, BCG12, pre- BCG13 and BCG15. Some patients discontinued with BCG (see Supplementary Table 2). Light blue arrow indicates pre-BCG1 time point which is used to calculate fold change in cytokine production. Purple arrows indicate important time points for TI, as patients did not receive BCG for weeks to months and thus represent the best ‘innate immune memory’ time points. **b:** TNF production by PBMCs after 24 hour stimulation with LPS at 8 time points during BCG therapy compared to pre-BCG1. **c:** IL-1β production by PBMCs after 24 hour stimulation with LPS at 8 time points during BCG therapy compared to pre-BCG1. **d**: TNF production by PBMCs after 24 hour stimulation with Pam3Cys (P3C) at 8 time points during BCG therapy compared to pre-BCG1. e: IL-1β production by PBMCs after 24 hour stimulation with P3C at 8 time points during BCG therapy compared to pre-BCG1. Fold change values for each individual patient are displayed as grey dots. Group values for each time point are displayed as median ± range in fold change compared to pre-BCG1. Two- tailed matched-pairs Wilcoxon signed-rank test was used to determine statistical significance in cytokine production between time points. Statistical significance was accepted at p<0.05 and indicated as follows: * p<0.05, ** p<0.01, *** p<0.001, **** p<0.0001. Number of data points per time point for b, c, e: pre-BCG1: 17, BCG2: 17, BCG6: 16, pre-BCG7: 14, BCG9: 13, pre- BCG10: 8, BCG12: 7, pre-BCG13: 3, BCG15: 3. Number of data points per time point for d: pre-BCG1: 15, BCG2: 15, BCG6: 13, pre-BCG7: 12, BCG9: 11, pre-BCG10: 7, BCG12: 6, pre-BCG13: 2, BCG15: 2.

Immediately after PBMC isolation, we performed *ex-vivo* stimulation experiments for 24 hours and measured production of four innate cytokines, i.e., IL-1β, IL-6, TNF, and IL-1 receptor antagonist (IL-1Ra) (see Methods). The fold change in cytokine production between pre-BCG1 (baseline, BCG naïve) and time points during BCG therapy was used as indicator of the TI response, in line with methods used by BCG vaccination studies^47^. We focused on pre-BCG7, pre-BCG10 and pre-BCG13 (i.e., pre-BCG maintenance time points). At these time points, the patients had not received a BCG instillation for multiple weeks or months and hence are most informative for TI^48^.

Before the start of the first 3-weekly BCG maintenance cycle (pre-BCG7, ±6 weeks after BCG6, Fig. 1a), the LPS-induced production of TNF and IL-1β was elevated compared to pre-BCG1 (median fold change (MFC) TNF: 1.70 (p=0.035); IL-1β: 1.50 (p=0.020)) (Fig. 1b and 1c). The TNF and IL-1β production after P3C stimulation was also increased, but not at the level of nominal statistical significance (MFC TNF: 1.26 (p=0.110); IL-1β: 1.05 (p=0.194)) (Fig. 1d and 1e). Similar findings were observed for IL-6 (LPS: MFC 1.21 (p=0.080); P3C: MFC 1.55 (p=0.058)) and IL-1Ra (LPS MFC 1.12 (p=0.020); P3C MFC 1.16 (p=0.058)) (Supplementary Fig. 1).

At the start of the second BCG maintenance cycle (pre-BCG10), approximately 3 months after BCG9, TNF and IL-1β production after stimulation was still increased compared to pre-BCG1 (Fig. 1). TNF was increased with a MFC of 1.62 and 1.87 for LPS and P3C stimulation, respectively (p=0.008 and p=0.016), and the MFC for IL-1β was 1.48 and 1.41 (p=0.016 and p=0.055). Production of IL-6 and IL-1Ra was not increased compared to pre-BCG1 (Supplementary Fig. 1). Between pre-BCG7 and pre-BCG10 there was no further increase in TNF production after LPS or P3C stimulation. The three patients from whom PBMCs were isolated at the pre-BCG13 time point did not show an increased production of TNF, IL-1β, IL- 6 or IL-1Ra compared to pre-BCG1, but small numbers prevent meaningful conclusions.

Thus, for the first time, we showed that BCG instillations induce long-lasting increased innate cytokine production by circulating monocyte-enriched PBMCs and we confirmed the presence of a persistent TI phenotype before the first and second BCG maintenance cycle.

We also assessed the short-term effects of BCG on innate immune responses during the BCG induction regimen and found that one week after the first BCG instillation (at BCG2) the total number of white blood cells in the circulation was increased (p<0.01, Supplementary Table 3) compared to pre-BCG1. More specifically, the neutrophil (p<0.01) and monocyte (p<0.01) cell populations were increased. The increase in neutrophil count after a single BCG instillation is in line with a study that found increased peripheral blood neutrophil counts in human newborns 3 days after vaccination with BCG^49^. For other time points, changes in innate immune populations were either less strong or absent (Supplementary Table 3). At BCG2, we also observed increased TNF, IL-1β and IL-1Ra production by PBMCs after *ex-vivo* stimulation with LPS or P3C. IL-6 production was not increased (Fig. 1 and Supplementary Fig. 1). IL-1β production after LPS and P3C stimulation remained increased at BCG6 compared to pre-BCG 1, whereas on a group level, TNF, IL-6 and IL-1Ra production was not increased (Fig. 1 and Supplementary Fig. 1).

### Epigenetic changes at cytokine-gene promoters in monocytes isolated after BCG instillations

Epigenetic modifications at the gene promoters of *IL6*, *TNF* and *IL1B* are hallmarks of trained immunity^26, 50^. Using ChIP-qPCR we assessed histone 3 lysine 4 trimethylation (H3K4me3) modifications, an activating epigenetic modification linked to BCG-induced TI^50^, at these promoters in a subset of 7 patients (those 7 that were included first in Tribute). Again, we focused on the pre-BCG1, pre-BCG7 and pre-BCG10 time points to assess innate immune memory. Surprisingly, in contrast to observations for cytokine production, we did not find an increase on group level in H3K4me3 modifications at pre-BCG7 or pre-BCG10 compared to pre-BCG1 (Fig. 2). An increased H3K4me3 signal was only observed in 2 to 4 out of 7 patients for the various cytokine gene loci. Importantly though, these 2 to 4 patients all had an increased TNF and IL-1β production after LPS stimulation. Furthermore, 3 out of the 4 patients with an increase in H3K4me3 at region 1 of the *IL6* promoter showed a strong increase in IL-6 production after LPS stimulation (MFC of 2 or higher). However, there were also patients with increased IL-6, TNF or IL-1β production upon LPS or P3C stimulation that showed decreased H3K4me3 signal.

**Fig. 2.**
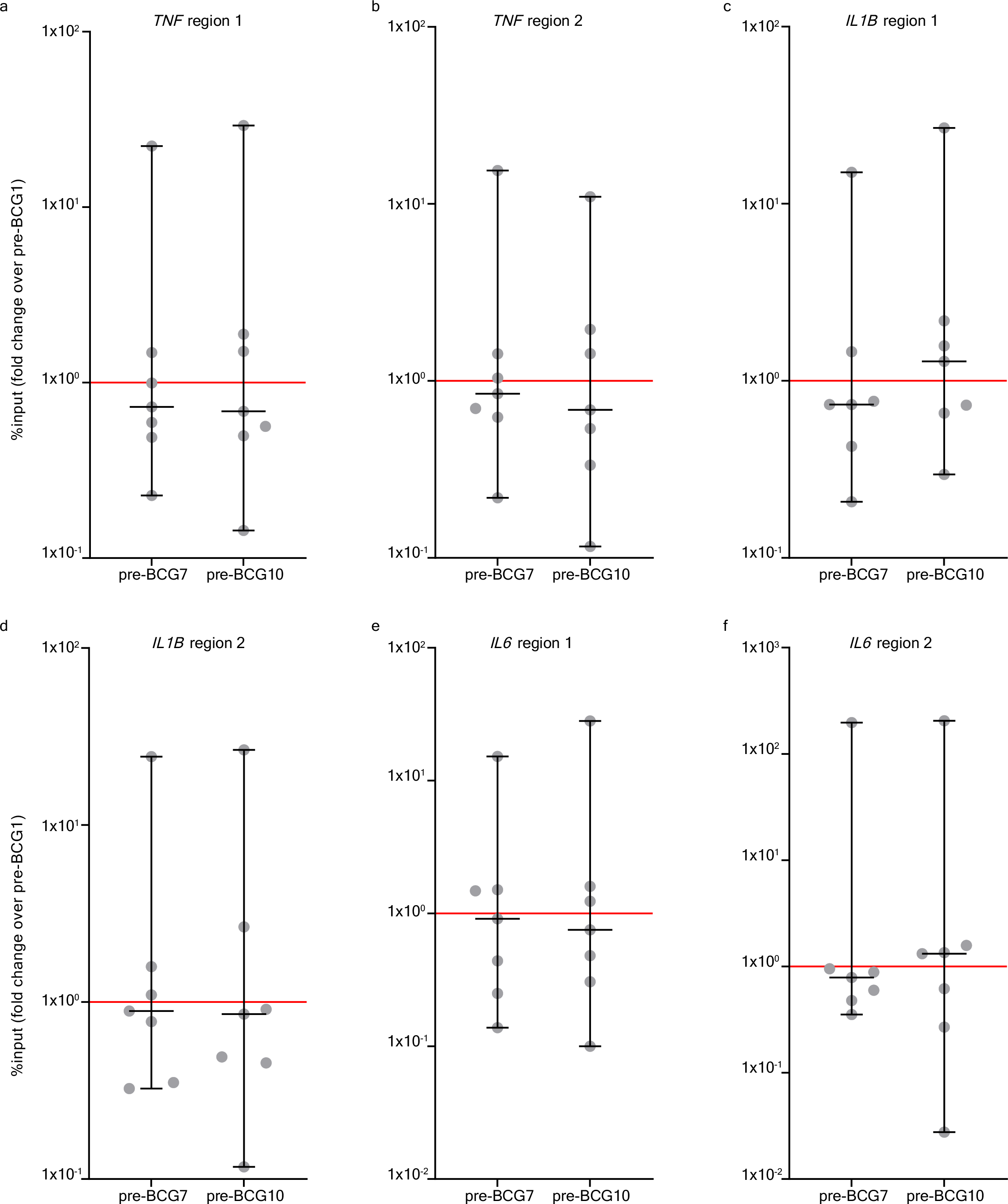
BCG instillations induce H3K4me3 modifications in monocytes. Results of the ChIP-PCR analysis for H3K4me3 modifications. H3K4me3 signal was calculated as %input and the fold change (of %input) is shown for pre-BCG7 and pre-BCG10 compared to pre-BCG1. The fold change is shown for 6 genomic regions in the promoter of *TNF*, *IL1B* and *IL6* (the primers are displayed in Supplementary Table 13). **a:** *TNF* region 1, **b:** *TNF* region 2, **c:** *IL1B* region 1, **d:** *IL1B* region 2, **e:** *IL6* region 1, **f:** *IL6* region 2. Two-tailed matched-pairs Wilcoxon signed-rank test was used to determine statistical significance in between time points. Statistical significance was accepted at p<0.05 and indicated as follows: * p<0.05, ** p<0.01, *** p<0.001 **** p<0.0001.

The analysis of H3K4me3 modifications via ChIP-PCR only covered specific parts of the promoter regions of *TNF*, *IL6* and *IL1B*. To exclude the possibility of missing out on modification signals in other parts of the promoter due to the selection of specific primers only, we investigated changes in H3K4me3 signal in other parts of these promoters as well via analysis of genome-wide ChIP-sequencing data of these 7 patients. This more in-depth analysis revealed that, on a group level, there were no changes in H3K4me3 signal at either the gene promoter or gene body of *IL6*, *TNF* or *IL1B*. This result was consistent for all comparisons between pre-BCG1 and post-BCG time points. Notably, both patients without (Fig. 3a) and with (Fig. 3b) recurrence of urothelial carcinoma after BCG therapy showed increases in H3K4me3 signal at the *TNF* promoter indicating no deterministic association with clinical outcome.

**Fig. 3.**
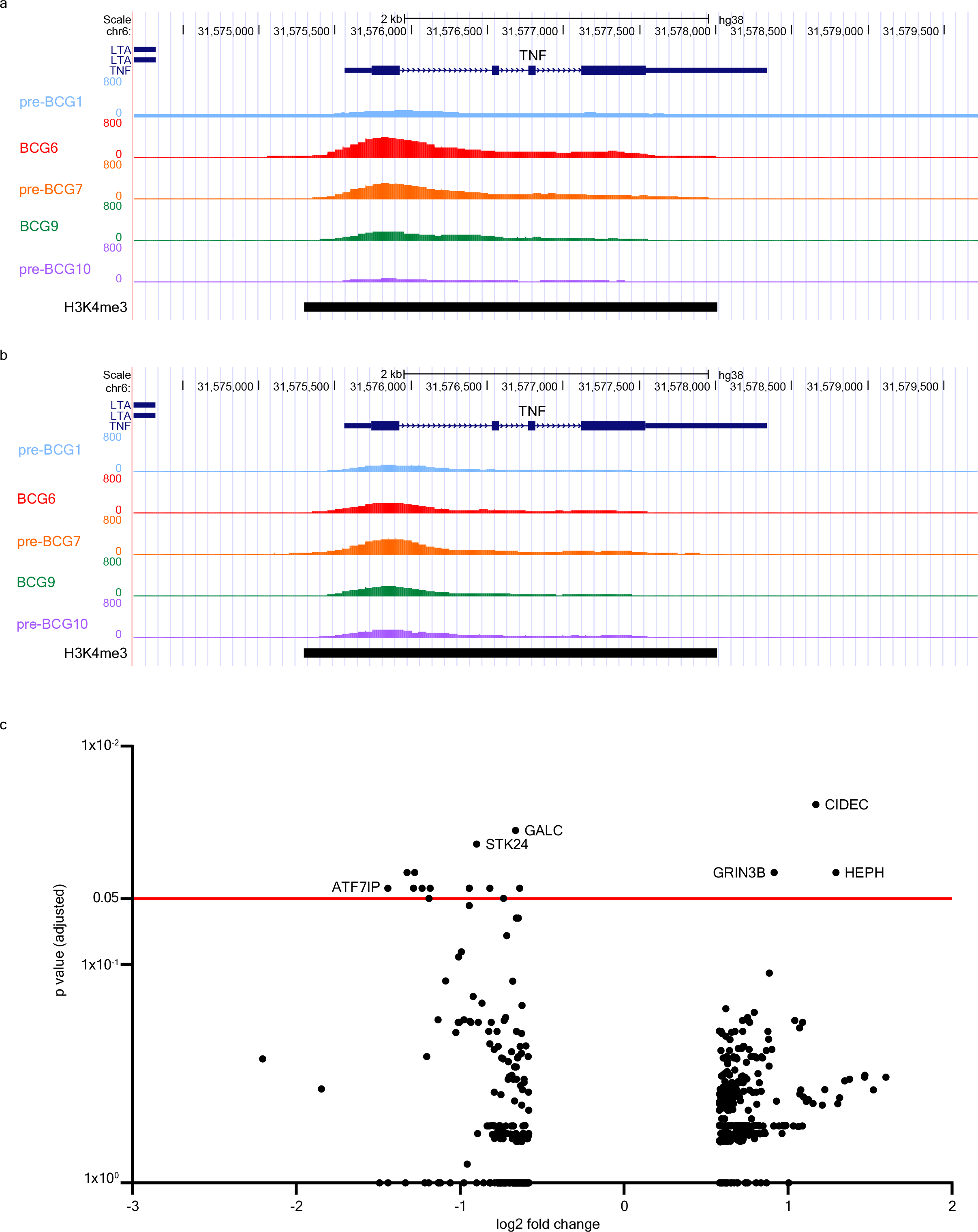
BCG instillations induce epigenetic modifications in genes involved in autophagy, IL-1β, and epigenetic enzymes. Results of the ChIP-sequencing analysis. a: UCSC genome tracks of H3K4me3 signal found around the *TNF* gene locus of a NMIBC patient with good clinical response to BCG. **b:** H3K4me3 signal near the TNF gene locus of a patient who had a recurrence after BCG therapy. H3K4me3 signal is displayed for pre-BCG1, BCG6, pre-BCG7, BCG9 and pre- BCG10. c: Volcano plot of ChIP-sequencing data. The x-axis displays the fold change (in log2) in H3K4me3 signal between pre-BCG1 and any of the post-BCG time points (BCG6, pre- BCG7, BCG9 and pre-BCG10; see Supplementary Table 4). The y-axis displays the p value adjusted for multiple testing using the Benjamini-Hochberg method. The red line indicates an adjusted p-value of 0.05.

### BCG instillations induce epigenetic modifications in genes involved with inflammasome signaling and chromatin remodeling

We further analyzed the ChIP-seq data to evaluate genomic regions that changed in H3K4me3 signal on a genome-wide scale. In total, there were 481 dynamic peaks (increasing or decreasing in H3K4me3 signal) that showed unadjusted p<0.05 between pre-BCG1 and any of the measured post-BCG time points (i.e., BCG6, pre-BCG7, BCG9, pre-BCG10) of which 375 were unique (Supplementary Table 4). Seventeen peaks showed an adjusted p<0.05. The strongest statistical evidence for an increase in H3K4me3 signal was found for pre-BCG10 and a region near *CIDEC*, coding for Cell-death-inducing-DFF45-like-effector-C and important for LPS-induced IL-1β production by epithelial cells^51^ (Fig. 3c; at pre-BCG10); the signal was increased at BCG6 and BCG9 as well (Supplementary Table 4). CIDEC is also involved in the formation of pro-inflammatory macrophage foam cells^52^ and controls inflammasome activation in mouse adipocytes^53^. In contrast, H3K4me3 near *ATF7IP*, a binding partner and regulator of *SETDB1*^54, 55^, which is a histone 3-lysine 9 (H3K9) methyltransferase, was decreased (Fig. 3c and Supplementary Table 4). We specifically assessed changes in H3K4me3 signal in 34 candidate TI genes (Supplementary Table 5), which are genes related to IL-1β/inflammasome signaling^26^, autophagy^29^, epigenetic enzymes^35, 36^, myelopoeisis^39^ and glycolysis^37^. Among the candidate genes, 3 regions showed a significant unadjusted p-value. *IL32* increased in H3K4me3 signal at BCG9 and pre-BCG10, *ATG16L2* and *KDM4E* were increased at pre- BCG10, and H3K4me3 near *TLR4*, the receptor for LPS, was decreased at BCG9. IL-32 plays a crucial role in the induction of TI by BCG^32, 33^. *ATG16L2* is involved in autophagy and KDM4E is an epigenetic enzyme which demethylates H3K9 and regulates trained immunity responses^36^. KDM4E and ATF7IP (indirectly) affect the methylation status of H3K9 which is interesting because the methylation status of H3K9 affects BCG-induced TI responses^35^.

Overall, the ChIP-seq results suggested that H3K4me3 modifications at the promoter regions of *TNF*, *IL6*, and *IL1B* can be induced by BCG instillations but not in all patients and without a clear link to clinical response in our small patient series. Analysis of the top hits and candidate genes suggested that genes involved with IL32, autophagy and H3K9-remodeling epigenetic enzymes, important for TI responses, may be epigenetically modified.

### BCG instillations upregulate genes involved in inflammasome activation

To determine how BCG instillations affect gene expression in monocytes, we performed RNA-sequencing on circulating Percoll-isolated monocytes isolated from six (out of 7 ChIP-seq) patients for timepoints pre-BCG1, BCG6 and pre-BCG7 (Fig. 4a). Note that the monocytes were not stimulated *ex-vivo* before RNA-seq analysis. We did not find statistically significant changes in gene expression of *TNF*, *IL6*, *IL1B* or *IL1RN* at BCG6 or pre-BCG7 compared to pre-BCG1 (Supplementary Table 6 and 7 and Supplementary Fig. 2). This is in line with BCG vaccination results described by Arts *et al*.^26^. Among the top hits for BCG6 versus pre-BCG1, *FANCA* (log2 FC: 0.41; padj=0.0199) and *APOL2* (log2 FC: 0.39; padj=0.0303) showed upregulation. The FANCA protein is involved in DNA damage repair during hematopoiesis^56^. Increased *APOL2* expression is associated with polarization of macrophages to the M1 (pro-inflammatory phenotype)^57^. Furthermore, two defined TI candidate genes, *GBP1* and *GBP2*, involved in inflammasome activation^58–61^, showed strong but borderline non-significant evidence for increased gene expression (log2 FC: 0.83; padj=0.052 and log2 FC: 0.54; padj=0.114). Among the candidate genes that did not reach statistical significance after multiple testing, we found that in addition to *GBP1* and *GBP2*, also *GBP4* and *GBP5* were upregulated (Fig. 4b) Interferon regulatory factor 1 (*IRF1*), an upstream transcription factor that regulates *GBP* gene expression, and the cytosolic DNA sensing receptor *AIM2*, which is activated by GBPs and IRF1^58, 62^, were upregulated as well (at punadj<0.05). Interestingly, *IL8,* a major chemokine for neutrophil recruitment, was downregulated (Supplementary Table 6). Motif enrichment analysis revealed that 17% of the scanned promoters from the differentially expressed genes at BCG6 are enriched for an interferon-sensitive response element (ISRE), an IRF responsive motif, compared to just 1.3% of all genes (Fig. 5a). These results suggest that IRFs, which are also transcription factors that control M1 macrophage differentiation^63, 64^, are important mediators between BCG exposure and gene induction^38^, also when BCG is instilled in the bladder. In line with this observation, we found upregulated expression of *IRF1* and *IRF9* at BCG6 (Fig. 5b) and increased IFNγ concentration in blood plasma (Fig. 5c).

**Fig. 4.**
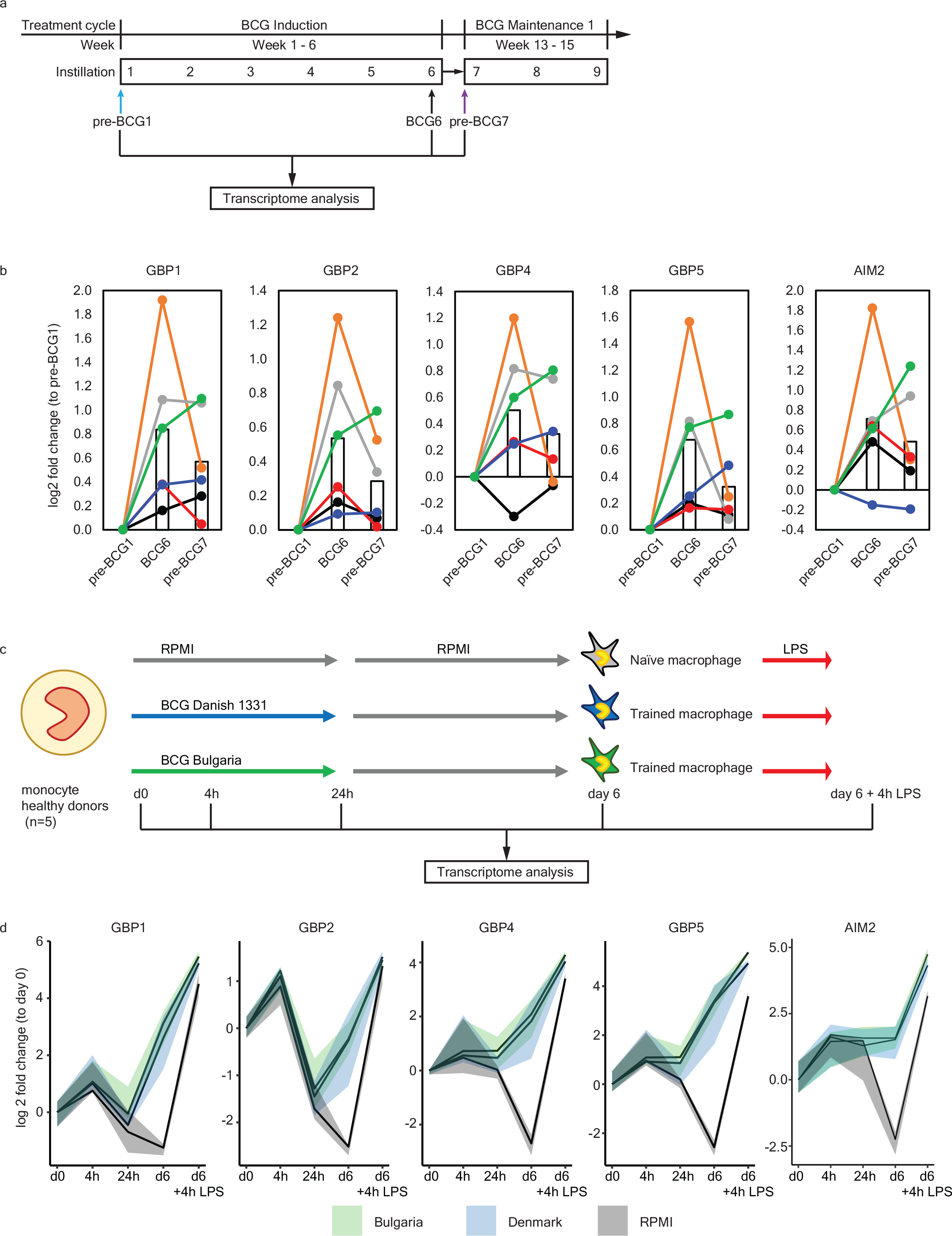
Upregulation of *GBPs* and *AIM2 in-vivo* and during induction of trained immunity in-vitro. **a:** Protocol for transcriptome analysis of monocyte-enriched cell suspensions before and during intravesical BCG therapy. Note that the cells were not stimulated *ex-vivo* before RNA- sequencing. **b:** Gene expression of *GBP1*, *GBP2*, *GBP4*, *GBP5* and *AIM2* at pre-BCG1, BCG6 and pre- BCG7. Each colour indicates an individual patient. The bar represents the mean gene expression of all 6 patients for each time-point. For the changes in mean gene expression between pre-BCG1 and BCG6: *GBP1* p=0.0001, padjust=0.053; *GBP2* p=0.0002, padjust=0.114; *GBP4* p=0.0070, padjust=1; *GBP5* p=0.0016, padjust=0.5051; *AIM2* p=0.0027, padjust=0.7203. And between pre-BCG1 and pre-BCG7 only *GBP1* and *AIM2* were significant: *GBP1* p=0.0084, padjust=1; *AIM2* p=0.0427, padjust=1. **c:** The *in-vitro* trained immunity protocol as described by Domínguez-Andrés *et al*^72^. Monocytes are isolated from the blood, seeded on a cell culture plate and subsequently incubated with a trained immunity-inducing stimulus (BCG), and a control condition (RPMI), for the first 24 hours. After 24 hours the training stimulus is washed away and the culture medium is refreshed. On day 6, the restimulation stimulus (LPS), or a negative control restimulation (RPMI) is added. After 24 hours of restimulation the supernatant is collected for cytokine measurements. For this particular experiment, cells were harvested at baseline (d0), after the first 4 hours (4h), 24 hours (24h), at day 6 before restimulation (d6), and at day 6 four hours after restimulation with LPS (d6 +4h LPS), and RNA was isolated and gene expression was measured. d: Changes in gene expression for *GBP1*, *GBP2*, *GBP4*, *GBP5* and *AIM2* are displayed as median (solid line), and 25^th^ and 75^th^ quartiles (shaded) for three conditions: BCG Bulgaria (green), BCG Denmark (blue), and RPMI (grey, negative control). The expression of all genes was significantly increased in D6-BCG-trained macrophages compared to RPMI macrophages (*GBP1* p =0.0001, padjust = 0.03; *GBP2* p = 7.78^-5^, padjust = 0.02; GBP4 p = 2.79^- 6^, padjust = 0.0013; GBP5 p = 7.40^-8^, padjust = 8.10^-5^, AIM2 p = 0.00015, padjust = 0.034). All adjusted p values in this figure were calculated using the Benjamini-Hochberg method.

**Fig. 5.**
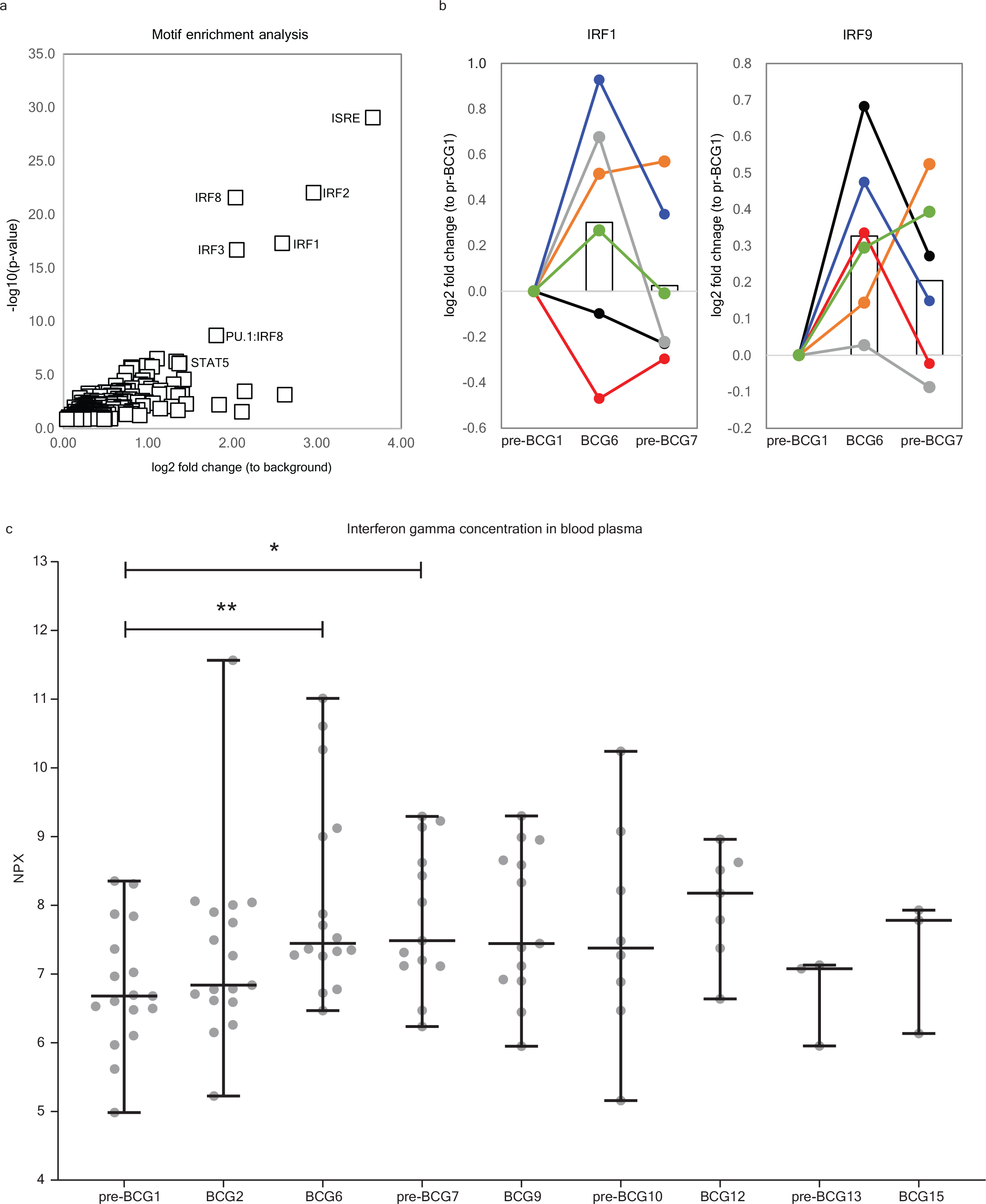
Increased activity of IRF transcription factors and higher IFNγ plasma concentration may drive upregulation of genes after BCG instillations. a: Motif enrichment analysis at the promoters of the upregulated genes between pre-BCG1 and BCG6 reveals that Interferon-Sensitive Response elements (ISRE) and Interferon Regulatory Factors (IRFs) are significantly more present at the promoter regions of upregulated genes. b: Gene expression of *IRF1* and *IRF9* is increased at BCG6 (*IRF1* p=0.0146, padjust=1; *IRF9* p=0.0004, padjust=0.1763) c: Concentration of IFNγ in blood plasma of Tribute patients at each time-point. Data are presented in NPX values, which is a log2 scale. Individual patient values are displayed as grey dots. Group values are displayed as median ± range. Two tailed matched-pair Wilcoxon signed-rank test was used to determine statistical significance between time points. Statistical significance was accepted at p<0.05 and indicated as follows: * p<0.05 and **p< 0.01. *** p≤ 0.001 **** p≤ 0.0001. Number of data points per time point: pre-BCG1: 17, BCG2: 17, BCG6: 16, pre-BCG7: 14, BCG9: 13, pre-BCG10: 8, BCG12: 7, pre-BCG13: 3, BCG15: 3.

At the start of BCG maintenance (pre-BCG7), 6 weeks after BCG induction there were no genes upregulated that reached the level of statistical significance after multiple testing (Supplementary Table 7).

A total of 26 genes remained upregulated at punadj<0.05 from BCG6 to pre-BCG7 and 27 genes remained downregulated (Supplementary Table 8). Among those that were upregulated, we identified four genes involved in inflammasome activation: *GBP1, AIM2, CASP5 and KCNMA1*^65, 66^, and one gene involved in positive regulation of autophagy: *RUFY4*^67–69^. Interestingly, three genes coding for hemoglobin subunits (*HBA1*, *HBA2*, *HBB*) remained downregulated.

GBPs regulate intracellular innate immune responses and activate the inflammasome for IL- 1β production^58–62^, also after LPS stimulation^58, 60, 70^. GBPs also mediate the release of DNA from vacuoles into the cytosol for activation of DNA-sensing receptors such as CGAS and AIM2^71^. Thus, our RNA-seq data suggested that GBP-inflammasome signaling and DNA-sensing by AIM2 may be increased.

### In-vitro trained immunity experiments with BCG are characterized by increased GBP- AIM2 inflammasome gene expression

The increased IL-1β production by PBMCs and the upregulation of AIM2 and GBPs suggests that inflammasome activity is enhanced and may be an important mechanism of TI during BCG therapy. To determine whether *GBP* and *AIM2* gene expression is also upregulated in monocytes trained by direct exposure to BCG vaccine, we performed *in-vitro* experiments with monocytes from 5 healthy donors using a standard TI protocol^37, 72^ (Fig. 4c), and measured gene expression at baseline, after 4 hours, 24 hours, 6 days, and 6 days plus 4 hours after LPS restimulation. In line with the RNA-seq data from the BCG-treated bladder cancer patients, we found that *in-vitro* induction of TI by two strains of BCG increased the expression of *GBP1*, *GBP2*, *GBP4*, *GBP5* and *AIM2* (Fig. 4d) compared to the non-BCG trained control condition, especially after LPS restimulation. These results suggested that BCG-trained monocytes, compared to the non-trained monocytes, had a primed AIM2-GBP inflammasome prior to restimulation which is then enhanced even more upon LPS restimulation. BCG may thus increase AIM2-mediated DNA-sensing and subsequently lead to increased inflammasome activation and IL-1β production.

### BCG bladder instillations decrease the risk of respiratory infections in NMIBC patients

BCG *vaccination* is associated with a reduced risk of pneumonia and influenza due to TI^27, 73–75^. If BCG *instillations* also induce a TI phenotype, we would expect to see a reduced frequency of respiratory infections. We studied this association by comparing self-reported data on respiratory infections experienced in the 18 months prior to reporting (i.e., between November 2018 and May 2020) from 407 BCG-treated NMIBC patients and 250 non-BCG treated NMIBC patients. Out of the 407 BCG-treated patients, 109 were considered BCG-exposed during the *whole* respiratory infection outcome assessment period; the remaining 298 BCG-treated patients were exposed during a part of this period or prior to this period (partially exposed) (see Methods and Supplementary Table 9 for patient characteristics). Results of the univariable regression analysis showed a 37% decreased risk of respiratory infections for the BCG-exposed vs BCG-unexposed group (OR 0.63 (95% CI 0.40-1.01)), and a 17% reduced risk for the partially BCG-exposed versus BCG-unexposed (OR 0.83 (95% CI 0.58-1.18)) (Table 1). The strongest evidence for a risk reduction in BCG-treated patients was seen for pneumonia and common cold. Multivariable analyses that included potential confounders showed very similar results (Supplementary Table 10). These findings suggest that BCG instillations induce heterologous protective effects against infectious diseases, in line with the induction of a systemic TI phenotype.

**Table 1.** Results of univariable association analysis for BCG exposure status and respiratory infections among NMIBC patients participating in the BlaZIB and UroLife studies. Comparisons were made between NMIBC patients that were considered BCG exposed, partially BCG exposed, and unexposed to BCG (see Methods).

|  | NMIBC BCG |  |  | NMIBC<br>no BCG | Unadjusted odds ratio (OR) (95% CI) |  |  |
| --- | --- | --- | --- | --- | --- | --- | --- |
|  | Total | Exposed | Partially<br>exposed |  | BCG exposed vs<br>no BCG | Partially BCG<br>exposed vs no<br>BCG | BCG vs no BCG |
| Total | 407 | 109 | 298 | 250 | - | - | - |
| Pneumonia<br>(%) | 23 (5.7) | 4 (3.7) | 19 (6.4) | 22 (8.8) | 0.40 (0.13-1.17) | 0.71 (0.37-1.34) | 0.62 (0.34- 1.14) |
| Bronchitis<br>(%) | 11 (2.7) | 5 (4.6) | 6 (2.0) | 10 (4.0) | 1.15 (0.39-3.46) | 0.49 (0.18-1.38) | 0.67 (0.28-1.59) |
| Laryngitis<br>(%) | 20 (4.9) | 6 (5.5) | 14 (4.7) | 16 (6.4) | 0.85( 0.32-2.24) | 0.72 (0.35-1.51) | 0.76 (0.38-1.49) |
| Flu (%) | 72 (17.7) | 19 (17.4) | 53 (17.8) | 50 (20.0) | 0.84 (0.47-1.51) | 0.87 (0.56-1.33) | 0.86 (0.58-1.28) |
| Cold (%) | 223 (54.8) | 53 (48.6) | 170 (57.0) | 155 (62.0) | 0.58 (0.37-0.91) | 0.81 (0.58-1.15) | 0.74 (0.54-1.02) |
| All above<br>RIs (%) | 254 (62.4) | 63 (57.8) | 191 (64.1) | 171 (68.4) | 0.63 (0.40-1.01) | 0.83 (0.58-1.18) | 0.77 (0.55-1.07) |

### Germline DNA variants in genes that affect TI induction are associated with recurrence and progression in BCG-treated NMIBC patients

We previously described an association between single-nucleotide polymorphism rs3759601 in *ATG2B*, that influenced both the *in-vitro* and *in-vivo* training effect of BCG, and response to BCG in NMIBC patients from the Nijmegen Bladder Cancer Study (NBCS)^29, 76^. Here, we extended our genetic research and assessed whether rare to common DNA variants in the 34 TI candidate genes (Supplementary Table 5) affect clinical outcome of BCG-treated patients. We analysed available exome chip data of 215 BCG-treated NMIBC patients from the NBCS (see Supplementary Table 11 for patient characteristics). Gene-based association analysis could be performed for 29 of the 34 genes and revealed no multiple testing-adjusted statistically significant finding for any of the candidate genes (Supplementary Table 12). However, an unadjusted p<0.06 was found for candidates *ATG2B*, *ATG7*, *ATG16L1*, *HNF1B*, and *EHMT2*, with either recurrence or progression after BCG induction or BCG induction and maintenance (Table 2). These results suggest that genetic variation in genes known to affect TI induction by BCG, i.e., autophagy genes^29^, a gene controlling myelopoiesis^39^, and a mediator of H3K9 methylation^35^, affect the clinical response after BCG therapy.

**Table 2.**
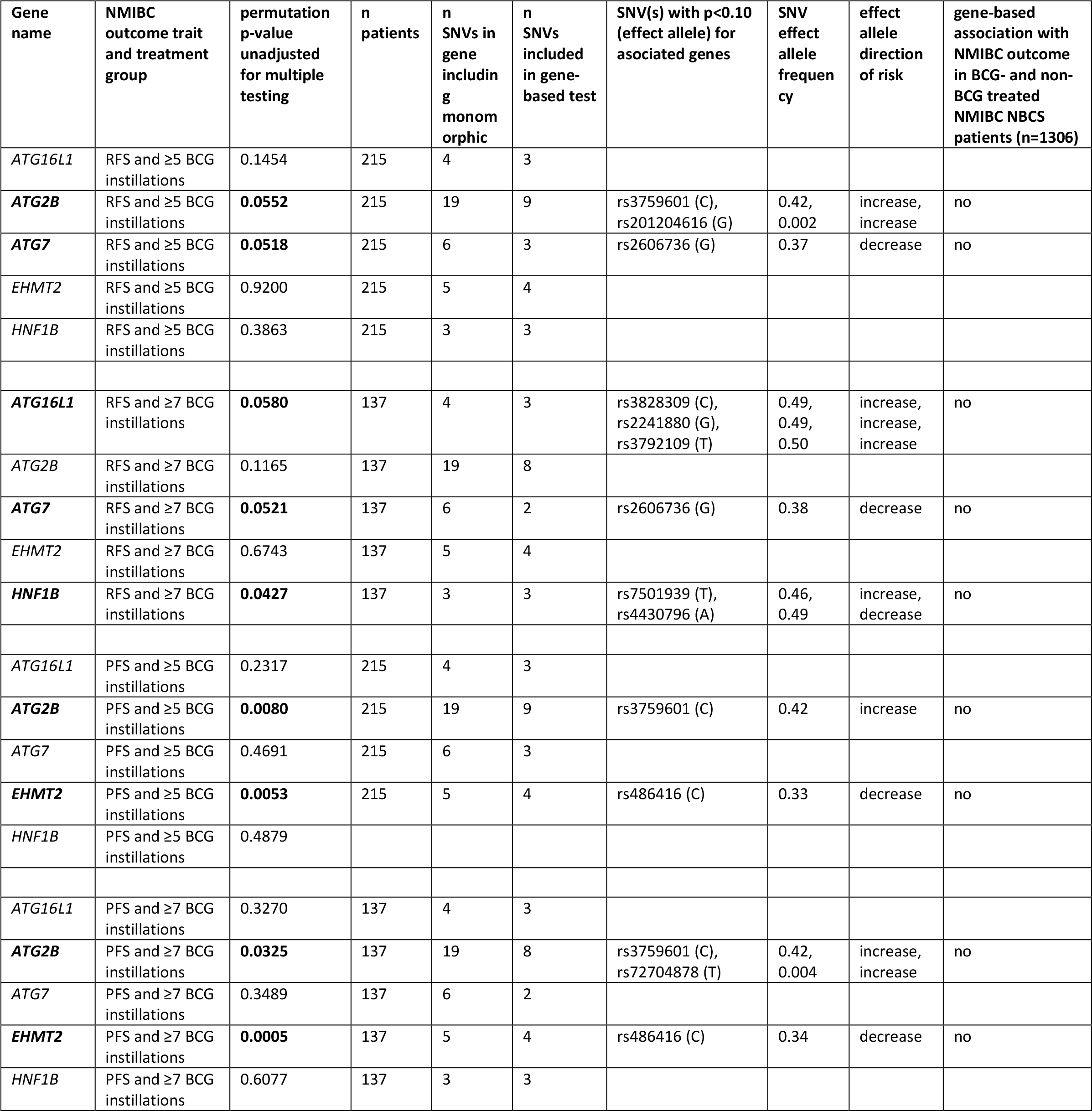
Results of the association analysis of exome chip variants in trained immunity candidate genes and clinical outcome in patients of the Nijmegen Bladder Cancer Study (NBCS). Depicted are the trained immunity candidate genes (Sup. Table 5) that showed an unadjusted p-value<0.06 for any of the four outcome/treatment combinations. The clinical outcomes which were assessed were tumor recurrence and disease progression in patients who received at least 5 induction BCG instillations or in patients who received at least BCG induction and one BCG maintenance cycle (defined as ≥7 instillations). RFS=recurrence free survival, PFS=progression free survival, SNV=single nucleotide variant.

## Discussion

In this study we showed for the first time that intravesical BCG treatment induces trained immunity at a systemic level, reflected by the increased cytokine production by PBMCs isolated from peripheral blood and increased protection against respiratory infections. Also, we found that germline DNA variants in genes important for the induction of TI influence the clinical oncological outcome after BCG instillations in NMIBC.

Previous research assessing the effects of intravesical BCG application on TI responses is scarce and provides incomplete evidence. Conti *et al.* showed that after 18 hours of *ex-vivo* LPS stimulation, monocytes from three BCG-treated NMIBC patients had a significantly higher production of TNF and IL-1α compared to monocytes from three non-BCG treated NMIBC patients or three healthy age-matched controls^45^. Furthermore, Buffen *et al.* found an increased production of TNF, IL-1β and IL-6 after the sixth BCG instillation compared to before the first BCG instillation^29^. Kim *et al*. observed that TNF production increased with consecutive BCG instillations of an induction cycle and peaked after the fourth BCG instillation, but for three out of seven patients the TNF production declined between the fourth and sixth instillation^46^. Graham *et al.* described variable post versus pre-BCG ratios of cytokines released by monocytes following LPS stimulation in 33 NMIBC patients during BCG induction, with approximately half of the patients showing an increase in ratios^77^. A limitation of all these studies is that they only assessed the cytokine production during a BCG induction cycle which may be influenced by the immunological processes of priming or tolerance^48^.

We now showed that repeated intravesical BCG instillations induce systemic TI and added new long-term data that showed that the TI phenotype is present at the start of BCG maintenance cycle 1 and 2, despite a 6 to 12-week time interval without BCG instillations. TI may already be induced after a single BCG instillation given the observed increased cytokine production one week after the first BCG instillation by us and others^46^. Importantly, some patients showed a decreased cytokine production capacity at the sixth BCG instillation, as also observed by Kim et al.^46^, which could indicate a phenotype of immune tolerance.

We also found large inter-patient variation in *in-vitro* cytokine production and in activating epigenetic modifications (H3K4me3) at promoters of hallmark trained immunity genes. Indeed, variability of the trained immunity responses in human volunteers has been well described, with only approximately half of the individuals being good trainers^78^. Our Tribute patient series was too small to test for associations between patient, tumor, and clinical characteristics and magnitude of TI responses but this is a relevant topic for further research. We could only study epigenetic modifications in a small number of patients, which may explain the lack of statistical significance of changes at the level of the entire group. Furthermore, we were able to assess only one histone mark (H3K4me3), and it may be hypothesized that other epigenetic modifications (such as H3K4me1, H3K27Ac, H3K9me3) could also contribute to the effects of BCG instillations at these time points.

Previous research showed that IL-1β is a key mediator controlling the induction of TI by BCG vaccination and that IL-1β was highly correlated with a reduction of viremia after BCG vaccination in an experimental model of viral infection in humans^26^. Furthermore, IL-1β signaling promotes glycolysis and proliferation of HSPCs in β-glucan trained mice^30, 31^. Functional reprogramming of HSPCs is also a hallmark of BCG-induced trained immunity in humans^39^ and mice^38^. Our data confirmed that IL-1β is also a key mediator of the TI phenotype induced by BCG instillations in humans. We showed that NMIBC patients treated with BCG have the capacity to produce increased amounts of IL-1β compared to pre-BCG1, and most importantly, this capacity is still present at the start of BCG maintenance 1 and 2. This indicates that circulating monocytes of BCG-treated NMIBC patients, which have proliferated in the bone marrow, have acquired a long-term functional program for increased IL-1β production upon immunological stimulation. Indeed, the increased expression of *GBPs* and *AIM2* in our RNA-seq data indicates that the BCG-treated patients had an increased capacity to produce IL-1β, which is controlled by the inflammasome. Our results are in line with previous studies which show that GBPs activate the inflammasome upon sensing cytosolic LPS, and thereby increase IL-1β production^59, 60, 79^. Furthermore, the increase in *GBP5* expression reflects data from Lim *et al.*, who showed increased *GBP5* expression in non-tumor urothelial tissue three months post-BCG compared to pre-BCG^8^. Finally, we showed that *GBP* and *AIM2* expression was increased in *in-vitro* TI experiments as well. Thus, BCG seems to increase *GBP* expression in circulating monocytes (*GBP5* also in the bladder^8^) and GBP-enhanced inflammasome activation may control TI responses both *in-vivo* and *in-vitro*.

The GBP-AIM2 inflammasome can also indirectly become activated via the signaling cascade of CGAS-STING-IRF1^62, 80^. CGAS (*MB21D1*) is a DNA-sensing receptor which activates the STING pathway which is important for the production of pro-inflammatory cytokines and type 1 interferon^81^. Recently, a recombinant BCG overexpressing c-di-AMP (a STING agonist) induced trained immunity and improved antitumor efficacy in animal models of NMIBC^82^. To what extent DNA-sensing by CGAS and AIM2 is involved in TI responses during BCG therapy in NMIBC patients remains to be elucidated. In our study we did not stimulate PBMCs with double-stranded DNA, so we cannot confirm whether TI responses to DNA stimulation are also increased. If this is indeed the case, one might hypothesize that (tumor- and BCG- derived) DNA-sensing in the bladder microenvironment may be increased and this may be one of the anti-tumor immune mechanisms of BCG instillations.

We already previously reported the association of genetic variation in *ATG2B*^29^ and *IL18*^83^ with clinical response after BCG therapy. Our exome chip analysis identified associations for additional autophagy genes and other genes that are known to affect TI induction directly at the epigenetic level (i.e., *EHMT2*^35^) and the level of hematopoiesis (i.e., *HNF1B*^39^). These genetic findings suggest a connection between TI and clinical efficacy of BCG in NMIBC. However, the genetic associations have not been replicated in independent NMIBC cohorts.

Also note that due to the small study sample and incomplete coverage of DNA variation via the exome chip, negative gene findings have low informative value.

In conclusion, we present comprehensive evidence from *in-vivo* and *ex-vivo* studies that BCG instillations in patients with NMIBC induce trained immunity. However, the TI responses induced varied widely between patients and this may also be the substrate of variable clinical responses, although this remains to be investigated in future studies. Most patients showed an augmented long-term systemic innate immune response, which may very well boost anti-tumor and anti-pathogen immune responses. Thus, exploiting TI responses may be of use to further increase efficacy of BCG therapy.

## Methods

### Study design and patient population of the Tribute study

We set up a prospective observational study in high-risk NMIBC patients (i.e. Ta high-grade or T1 with or without concomitant carcinoma in situ (CIS)) that were scheduled to start BCG therapy at the urology departments of two Dutch academic hospitals (Radboud university medical center Nijmegen and Erasmus MC Rotterdam). Inclusion for this study, named Tribute (acronym for ‘TRained Immunity induced by BCG in UroThElial carcinoma’), was initiated in June 2018 and ended in April 2021. Eligibility criteria are given in Supplementary Table 3. In short, all patients were BCG naive and had primary or recurrent HR-NMIBC, with or without CIS. All patients were free of visible papillary tumor at the start of BCG therapy as determined via re-TURT or negative cystoscopy and/or cytology at most 6 weeks before start of BCG therapy. Patients with tumor stage >cN0 or cM1 were excluded from participation, as well as those with upper urinary tract tumors or another malignancy other than basal cell carcinoma of the skin or prostate cancer under active surveillance. The BCG treatment schedule in this observational study was up to the treating urologist and was based on a standard regimen of a 6-week induction course followed by 3-weekly maintenance courses. Clinical data and follow-up were collected from the medical files of the participating patients. A baseline questionnaire was used to extract patient information on e.g. smoking and BCG vaccination status and a diary was used to collect data on smoking and co-medication during BCG therapy. The study received ethical approval from the medical research ethics committee Arnhem-Nijmegen (METC number: NL60341.091.17).

### Blood sample collection

Blood for the Tribute study was collected (40-50mL in EDTA tubes) at nine time points during the first year of BCG therapy and was drawn at the outpatient clinic, either directly before the BCG bladder instillation or 5-20 minutes after the instillation. Blood was drawn directly before the BCG bladder instillation at study visits: pre-BCG1, pre-BCG7, pre-BCG10 and pre-BCG13.

Blood was drawn 5 to 20 minutes after applying the BCG instillation at study visits: BCG2, BCG6, BCG9, BCG12 and BCG15 (Fig. 1a).

### Whole blood cell counts

Complete blood leukocyte counts were determined in 100uL EDTA whole blood using a Sysmex XN-450 analyzer (Sysmex) within 5 hours after blood draw.

### PBMC isolation

Isolation of PBMCs and purification of monocytes was performed as described in previously published articles by our group^26, 34, 84^. Briefly, PBMCs were isolated by density centrifugation of Ficoll-Paque (GE healthcare, UK) within 5 hours after blood draw. Cells were washed twice in PBS and resuspended in RPMI culture medium (Roswell Park Memorial Institute medium; Invitrogen, CA, USA) supplemented with 50 mg/mL gentamicin, 2 mM Glutamax (GIBCO), and 1 mM pyruvate (GIBCO) and diluted to a concentration of 5x10^6^/mL for *in-vitro* PBMC stimulation experiments.

After PBMC isolation, purification of monocytes was performed using a hyper-osmotic density gradient medium Percoll (Sigma-Aldrich). The PBMC suspension was adjusted to a concentration of 50-60x10^6^ cells/mL and subsequently applied on top of the hyper osmotic Percoll solution, after which the cells were centrifuged at 580g for 15 minutes at room temperature, with no brake and slow acceleration (Sorvall Legend RT). The interphase layer was collected with a Pasteur pipette and the cells were washed with cold PBS once. The cells were diluted in 2-3 mL RPMI and this monocyte-enriched cell suspension was further used for epigenetics analyses, and RNA isolation.

### PBMC stimulation experiments

PBMC stimulation experiments were performed within 6 hours after drawing of blood. PBMC stimulation experiments were performed on round bottom 96 wells plates (Greiner). In each well 100uL of PBMC cell suspension (5x10^6^/mL) was seeded. Subsequently, 100uL of stimulus was added. The final cell culture concentrations of the used stimuli were as follows: 10 ng/mL *Escherichia coli* LPS (serotype 055:B5, Sigma-Aldrich), 10 µg/mL Pam-3-Cys (EMC micro- collection). The experiments were performed in duplicate. PBMCs were stimulated for 24 hours in an incubator at 37°C and 5% CO2. After 24 hours the plates were spun for 8 minutes at 1400RPM (Rotina 380R Hettich) and supernatants were collected and stored at -80°C until analysis.

### Cytokine measurements in PBMC supernatants

Cytokine concentrations in PBMC supernatants were measured to determine whether cytokine production after non-specific stimulation for 24 hours was altered throughout BCG instillation therapy. These measurements were performed for both duplicates and the mean value of both duplicates was used for data analysis. IL-1β, IL-6, TNF and IL-1 receptor antagonist (IL-1Ra) were measured in thawed supernatants using commercial ELISA kits (R&D Systems). IL-1Ra was measured because it inhibits the binding and bioactivity of IL-1β^85^, thus we expect IL-1Ra to inhibit TI responses.

### Chromatin Immuno Precipitation for histone 3 lysine 4 trimethylation (H3K4me3 ChIP)

Isolated monocyte-enriched suspensions were fixed with 1% formaldehyde (Sigma). Fixed cell suspensions were sonicated using a Bioruptor Pico (Diagenode) for 7 cycles (30s on; 30s off). For each ChIP, chromatin of 0.5x10^6 cells was incubated with 254 uL dilution buffer, 12uL protease inhibitor cocktail (25x), and 1µg of H3K4me3 antibody (Cell Signaling Technology, Danvers USA) and incubated overnight at 4C with rotation. Protein A/G magnetic beads were washed in dilution buffer with 0.15% SDS and 0.1% BSA, added to the chromatin/antibody mix and rotated for 60 min at 4C. Beads were washed with 500ml buffer for 5 min at 4C with five rounds of washes. After washing, chromatin was eluted using elution buffer for 20 min. Supernatant was collected, 8 µL 5M NaCl, 2 µL proteinase K were added and samples were incubated on a shaking heat-block for 4 hours at 1000rpm, 65C. This elution procedure was also repeated for chromatin input samples, where 0.5x10^6 cells were used for input as well.

As final step, the DNA was isolated using QIAGEN MinElute PCR purification Kit and eluted in 20 µL elution buffer.

### ChIP-qPCR

We included the first 7 Tribute patients for ChIP-qPCR to have access to chromatin from long-term time points (i.e., pre-BCG10). Immunoprecipitated chromatin was used for RT-qPCR analysis. For qPCR analysis the input samples were diluted 25 times and the ChIP samples were diluted 3 times. Primers used in the reaction are listed in Supplementary Table 12. Samples were analysed with a comparative Ct method on the StepOne PLUS qPCR machine (Thermo Fisher Scientific) using SYBR green (Invitrogen) in accordance with the manufacturer’s instructions.

### Chromatin Immuno Precipitation sequencing (ChIP-seq)

ChIP-seq was performed for the first 7 patients that were included in Tribute. Illumina library preparation was performed using the KAPA HyperPrep kit (KAPA Biosystems). Library concentration was measured using the KAPA Library Quantification Kit (KAPA Biosystems); library size was determined using the BioAnalyzer High Sensitivity DNA Kit (Agilent). Sequencing was performed using an Illumina NextSeq500, and 42-bp paired-end reads were generated.

### Processing of genome-wide histone modification data

Chromatin Immunoprecipitation and sequencing (ChIP-seq) reads were aligned to human genome assembly hg38 (NCBI version 38) using bwa^86^. BAM files were first filtered to remove the reads with mapping quality less than 15, followed by fragment size modelling. MACS2 was used to call the peaks. Peaks from all samples were merged into a single ‘H3K4me3 peaks’ bed file and reads per peak were counted using bedtools coverage^87^. Data (reads/peak) were normalized using the R package DESeq2 and then pairwise comparisons were performed.

### RNA isolation and RNA-sequencing

After isolation, 1 – 5 x 10^6^ cells of the monocyte-enriched cell suspension were used for RNA extraction. The cells were spun down at 500g for 5 minutes. Supernatant was removed and the cell pellet was resuspended with 500µL RNAlater solution (Thermo Fisher Scientific) and kept at room temperature for 20 minutes. The resuspended solution was stored at -80C until RNA isolation. The cell pellets which were frozen in RNAlater solution were thawed to RT and centrifuged for 5 minutes at 300g. The cell pellets were then dissolved in RLT buffer (Qiagen) for 20 minutes. RNA was isolated using RNeasy Mini Kit (Qiagen) according to manufacturer’s protocol, and including DNAse treatment.

RNA sequencing (RNA-seq) was performed by BGI Genomics (Denmark) for 6 patients (all included in ChIP-seq). The DNBseq RNA transcriptome PE101 pipeline was used with 30 million reads per sample. Read length was 100bp. After sequencing, the raw reads were filtered. Data filtering included removing adaptor sequences, contamination and low-quality reads from raw reads. To infer gene expression levels, RNA-seq reads were aligned to hg19 human transcriptome using Bowtie^88^. Quantification of gene expression levels as RPKM was performed using MMSEQ^89^. Reads/transcripts were normalized using DEseq2 and pair-wise comparisons were performed.

### Statistical analysis for ChIP-seq and RNA-seq datasets

Differentially expressed genes were identified using DEseq2 with fold change > 1.33 and p-value <0.05, with a mean RPKM > 1. For longitudinal comparisons, the outputs of all pair-wise comparisons (e.g. pre-BCG1 vs BCG6, pre-BCG1 vs pre-BCG7) were combined and separated into groups using k-means clustering in MeV^90^. Dynamic H3K4me3 peaks were identified using DEseq2 with change in signal determined as fold change > 1.5 and unadjusted p-value <0.05, with a mean reads/peak > 25^91^. The Benjamini-Hochberg method was used to correct for multiple testing in DEseq2 and the adjusted p-values are shown in the supplementary tables. Genes of interest were downloaded from Kyoto Encyclopedia of Genes and Genomes (KEGG)^92^. For gene ontology, H3K4me3 peaks were assigned to the nearest transcription start site (TSS) within 1Mb using the GREAT tool^93^. Promoter H3K4me3 peaks were identified as those within 5kb from a transcription start site using bedtools. Motif analysis was performed on gene promoters using HOMER findMotifs^94^. Motif enrichment was calculated using the hypergeometric distributions, comparing the input promoter list to a background list of promoters. In addition to the p-value, we used a fold change cut-off of >2 to designate a motif as significantly enriched.

### Analysis of genes of interest in *ex-vivo* BCG RNA sequencing data

Genes of interest were validated for BCG responsiveness *ex-vivo* using the standard trained immunity model^72^ with a time-resolved transcriptome analysis approach^91^. Briefly, monocytes from 5 healthy volunteers were exposed to BCG *ex-vivo* for 24h and RNA was collected at baseline, 4h, 24h, day 6 (5 days after removal of stimulus), and day 6 + 4 hours of LPS stimulation. Time-course RNA expression data, as normalized counts, was extracted for *GBP1*, -*2*, -*3*, -*4*, *5* and *AIM2* and plotted as median log2 fold change relative to baseline. Raw data files for *ex-vivo* RNA sequencing are available at GSE168468.

### Measurement of IFNγ in blood plasma

Measurement of IFNγ in blood plasma was performed for the following number of patients and time points: pre-BCG1: 17, BCG2: 17, BCG6: 16, pre-BCG7: 13, BCG9: 13, pre-BCG10: 8, BCG12: 7, pre-BCG13: 3, BCG15: 3. Measurement of circulating IFNγ in blood plasma was performed using the commercially available Olink Proteomics AB (Uppsala Sweden) inflammation panel (92 inflammatory proteins). Data values are depicted as Normalized Protein eXpression (NPX), which is Olink’s arbitrary unit and is in Log2 scale.

### Statistical analysis of cytokine levels in supernatants and blood plasma

Descriptive statistics are presented as median ± range, as indicated in the legend of each figure, unless otherwise stated. The statistical significance of the differences between study visits was evaluated using matched pair two-tailed Wilcoxon signed-rank test. In figures, asterisks denote statistical significance (*, p<0.05; **, p<0.01; ***, p<0.001; ****, p<0.0001). Statistical analysis was performed in GraphPad PRISM 8.

### Study into respiratory infections in NMIBC patients from the BlaZIB and UroLife studies

NMIBC patients with date of diagnosis between May 2014 and April 2020 who are participating in BlaZIB^95^ or UroLife^96^, two ongoing observational studies (METC Region Arnhem-Nijmegen file numbers 2017-3240 and 2013-494, respectively), were invited in May 2020 to participate. Only those patients were invited that provided consent within the BlaZIB or UroLife study to be approached for additional research and for whom an email address was available. Patients were invited via email to fill out a digital questionnaire. A total of 1230 patients were invited of which 678 consented and filled out the questionnaire.

The questionnaire contained questions on bladder instillations, disease history, smoking, vaccinations for flu and tuberculosis (BCG), respiratory infections (i.e. pneumonia, bronchitis, laryngitis, flu, and common cold) between November 2018 and May 2020, and questions on COVID-19 symptoms. Clinical and treatment characteristics for the patients were extracted from the Netherlands Cancer Registry (NCR) in the second half of 2020, but follow-up on treatment was incomplete for some patients. Data from both the questionnaires and the NCR were used to define the (timing of) BCG exposure for the patients. Given the period of outcome assessment of November 2018 to May 2020, we defined a subgroup of BCG-treated NMIBC patients that we considered as exposed to BCG during the whole outcome assessment period, i.e. first BCG instillation prior to October 15 2018 and final BCG instillation after April 2019. Here, we assumed that a TI phenotype is fully induced two weeks after the first instillation and will be present up to one year after the last BCG instillation. The other BCG-treated patients were either partially exposed during the outcome assessment period or exposed prior to the outcome assessment period.

Statistical analyses entailed the description of characteristics in cross tables for comparison of groups that differed in their exposure to BCG. Univariable logistic regression analysis was performed to generate odds ratios (OR) and 95% confidence intervals (CIs). Multivariable logistic regression analysis was performed to allow for adjustment of factors that are known to affect the incidence of respiratory infections and may be associated with BCG exposure or NMIBC patient status. More specifically, we considered older age, male sex, lack of flu vaccination (in the year 2018 and/or 2019), smoking cigarettes, and history of chronic lung disease (i.e. asthma, COPD, chronic bronchitis, lung emphysema) as potentially relevant respiratory infection-risk increasing covariables.

### Genotyping and analysis of exome DNA variation in BCG-treated NMIBC patients

BCG-treated NMIBC patients were included from the previously described Nijmegen Bladder Cancer Study^76^. NBCS patients were genotyped using the Illumina HumanExome BeadChip (‘exome chip’) containing over 240,000 exonic markers of which ∼90% with minor allele frequency (MAF) <5%. Variants were called and cleaned according to^97^ and as an additional quality check, cluster plots of variants that showed a difference of >5% in MAF compared to European samples from the 1000 Genomes project were visually checked and adjusted, where appropriate. Additionally, standard quality control was performed using PLINK v1.9 which included removal of samples based on call rate, gender mismatch, relatedness, heterozygosity and population outliers, and removal of variants with call rate <98%, that were monomorphic, not in HWE, or should be treated with caution (https://genome.sph.umich.edu/wiki/Exome_Chip_Design). Also, duplicate and tri-allelic variants were removed. After QC, 92,364 exome chip variants were available for analysis in a total of 215 NMIBC patients that had received at least 5 BCG induction instillations (see Supplementary Table 11 for patient characteristics). Given the known clinical benefit of BCG maintenance cycles, we also performed analyses in the subset of patients that received at least one maintenance cycle (defined as at least 7 instillations). Gene-based analyses of candidate TI genes were performed using the coxKM R-package^98^ (linear kernel and default settings). Five of the 34 candidate genes could not be analyzed (*IL1B* and *KDM4E*: no exome chip variants; *ASC* and *ATG5*: only monomorphic DNA variants in study population; *LAMP5*: X chromosome, not analyzed). Multiple testing-adjusted p-value was set at 4.3*10^-4^ (29 genes and 4 outcome/treatment combinations). Single variant analyses were performed using univariable Cox proportional hazards regression analysis and an additive genotype model. Recurrence was defined as a new, histologically confirmed bladder tumor following at least one tumor-negative urethrocystoscopy or following two surgical resection attempts for the primary bladder tumor. Disease progression was defined as a shift to a higher tumor stage and/or grade. RFS and PFS were defined as the time period between the initial transurethral resection of the tumor and the first event (recurrence or progression, respectively), date of censoring or date of five-year follow-up, whichever came first, as described before^99^.

### Data availability

Raw data files for RNA-sequencing and ChIP-sequencing are deposited in the NCBI Gene Expression Omnibus under the reference series number GSE190530.

## Acknowledgements

We thank all participants of Tribute, UroLife, BlaZIB and NBCS for their participation. We thank H. Dijkstra and H. Lemmens for support with ELISA analysis. We thank M.A.B. Rosmalen, H. van de Pol, J.C.M. Smits-van de Camp, J.M. Jansen-Jansen and V.E. de Kleijnen for their help with blood and urine collection for Tribute and J.S.F. Maurits for help with data collection for the respiratory infection study. The authors thank the registration team of the Netherlands Comprehensive Cancer Organisation (IKNL) for the collection of data for the Netherlands Cancer Registry as well as IKNL staff for scientific advice.

M.G.N. was supported by an ERC Advanced Grant (#833247) and a Netherlands Organization for Scientific Research Spinoza Grant (NWO SPI 94-212). L.A.B.J. was supported by a Competitiveness Operational Programme grant of the Romanian Ministry of European Funds (P_37_762, MySMIS 103587). S.H.V. is supported by a grant from the Netherlands Organization for Scientific Research (NWO Vidi 91717334). B.N. is supported by an NHMRC (Australia) Investigator Grant (APP1173314) and Project Grant (APP1157556). The BlaZIB study was financially supported by the Dutch Cancer Society (IKNL 2015-7914). The UroLife study was financially supported by Alpe d’HuZes/Dutch Cancer Society (KUN 2013-5926) and Dutch Cancer Society (2017-2/11179). Funding for the exome chip study was provided by the Netherlands Organization for Scientific Research under award number 184021007 and made available as a Rainbow Project of the Biobanking and Biomolecular Research Infrastructure Netherlands (BBMRI-NL). Genetic analyses were carried out on the Dutch national e-infrastructure with the support of SURF Cooperative.

---

Supplementary Fig. 1 –Production of IL-6 and IL-1Ra by PBMCs upon *ex-vivo* **stimulation with heterologous stimuli after BCG instillations**

**a:** Study schedule of the Tribute study. Blood was collected and PBMCs were isolated at three time points during the BCG induction cycle: pre-BCG1, BCG2 and BCG6; and two time points during each subsequent BCG maintenance cycle: pre-BCG7, BCG9, pre-BCG10, BCG12, pre- BCG13 and BCG15. Some patients discontinued with BCG (see Supplementary Table 2). Light blue arrow indicates pre-BCG1 time point which is used to calculate fold change in cytokine production. Purple arrows indicate important time points for TI, as patients did not receive BCG for weeks to months and thus represent the best ‘innate immune memory’ time points.

b: IL-6 production by PBMCs after 24 hour stimulation with LPS at 8 time points during BCG therapy compared to pre-BCG1.

**c:** IL-6 production by PBMCs after 24 hour stimulation with P3C at 8 time points during BCG therapy compared to pre-BCG1.

**d:** IL-1Ra production by PBMCs after 24 hour stimulation with LPS at 8 time points during BCG therapy compared to pre-BCG1.

**e:** IL-1Ra production by PBMCs after 24 hour stimulation with P3C at 8 time points during BCG therapy compared to pre-BCG1.

Individual patient fold change values are displayed as grey dots. Group values for each time point are displayed as median ± range in fold change compared to pre-BCG1. Two tailed matched-pair Wilcoxon signed-rank test was used to determine statistical significance in cytokine production between time points. Statistical significance was accepted at p<0.05 and indicated as follows: * p<0.05 and **p< 0.01. *** p≤ 0.001 **** p≤ 0.0001. Number of data points per time point for b: pre-BCG1: 17, BCG2: 17, BCG6: 16, pre-BCG7: 13, BCG9: 13, pre- BCG10: 8, BCG12: 7, pre-BCG13: 3, BCG15: 3. Number of data points per time point for c, d, e: pre-BCG1: 17, BCG2: 17, BCG6: 16, pre-BCG7: 14, BCG9: 13, pre-BCG10: 8, BCG12: 7, pre-BCG13: 3, BCG15: 3.

Supplementary Fig. 2 – Volcano plots of the differentially expressed genes at BCG6 and pre-BCG7.

a: Volcano plot of RNAseq data comparing gene expression at pre-BCG1 with BCG6. Genes highlighted in red are downregulated and genes highlighted in blue are upregulated. Genes in grey either did not increase in expression (FC<1.3) or did not meet statistical significance (p>0.05).

b: Volcano plot of RNAseq data comparing gene expression at pre-BCG1 with pre-BCG7. Genes highlighted in red are downregulated and genes highlighted in blue are upregulated. Genes in grey either did not increase in expression (FC<1.3) or did not meet statistical significance (p>0.05).

Supplementary Table 1 - Inclusion and exclusion criteria for the Tribute study.

**Supplementary** Table 2 - Characteristics of patients included in the Tribute study.

^1^The highest tumor stage/grade is reported as the highest stage or grade which was found in the pathology report at the transurethral resection of the bladder tumor (TURT) or re-TURT of study inclusion.

Supplementary Table 3 - Overview of the concentration of circulating leukocytes in whole blood measured at different time points during BCG therapy in the Tribute study.

Values are displayed as median ± range. Two-tailed matched-pairs Wilcoxon signed-rank test was used to determine statistical significance in fold change between time points. Statistical significance was accepted at p<0.05.

Supplementary Table 4 - Results from the ChIP-seq analysis.

Comparisons in H3K4me3 signal were made between pre-BCG1 and any of the measured post-BCG time points (i.e., BCG6, pre-BCG7, BCG9 and pre-BCG10). Candidate trained immunity genes are indicated in the final column.

Supplementary Table 5 - List of selected candidate trained immunity genes

The mechanism/pathway based on which this gene is important for trained immunity is displayed in the second column and the reference, if available, is shown in the third column.

Supplementary Table 6 - Table with the differentially expressed genes between Pre- BCG1 and BCG6.

This table shows the differentially expressed genes at punadj<0.05 between pre-BCG1 and BCG6. Candidate trained immunity genes are indicated in the last column.

Supplementary Table 7 - Table with the differentially expressed genes between Pre- BCG1 and Pre-BCG7.

This table shows the differentially expressed genes at punadj<0.05 between pre-BCG1 and pre- BCG7. Candidate trained immunity genes are indicated in the last column.

Supplementary Table 8 – Maintained upregulated and downregulated genes.

This table shows the differentially expressed genes which were upregulated at BCG6 and remain upregulated at pre-BCG7, as well as the downregulated genes at BCG6 which remain downregulated at pre-BCG7 (all at punadj<0.05). Candidate trained immunity genes are indicated.

Supplementary Table 9 - Characteristics of NMIBC patients from the BlaZIB and UroLife studies that were part of the study into BCG exposure and respiratory infections.

**Supplementary** Table 10 - Results of the multivariable association analysis for BCG exposure and respiratory infections among NMIBC patients in the BlaZIB and UroLife studies.

* Adjusted for age (continuous variable), sex (female versus male), smoking (3 categories), chronic lung disease (yes vs no), flu vaccination in 2018/2019 (yes vs no). CI=confidence interval.

Supplementary Table 11 - Characteristics of the Nijmegen Bladder Cancer Study population that was included in the exome chip analyses for NMIBC recurrence and progression.

CIS = Carcinoma in situ; BCG = Bacillus Calmette-Guérin; iBCG = induction therapy of BCG; mBCG = maintenance therapy of BCG; IQR = interquartile range; ^1^highest tumor stage/grade based in transurethral resection of the (bladder) tumor (TURT) and reTURT combined. ^2^ restricted to the first five years after diagnosis.

Supplementary Table 12 – Results of the gene-based analysis of the 34 TI candidate genes using exome chip data from 215 BCG-treated NMIBC patients.

The clinical outcomes which were assessed were recurrence free survival and progression free survival. Each sheet of the excel file shows the candidate trained immunity genes and their association with RFS or PFS after 5 or more and 7 or more BCG instillations. RFS=recurrence free survival. PFS=progression free survival.

Supplementary Table 13 - Primers that were used for ChIP-PCR analysis.

