## Supplemental Figures and Tables for "Intravesical BCG in patients with non-muscle invasive bladder cancer induces trained immunity and decreases respiratory infections"

Supplemental Data

Supplementary Figure 1

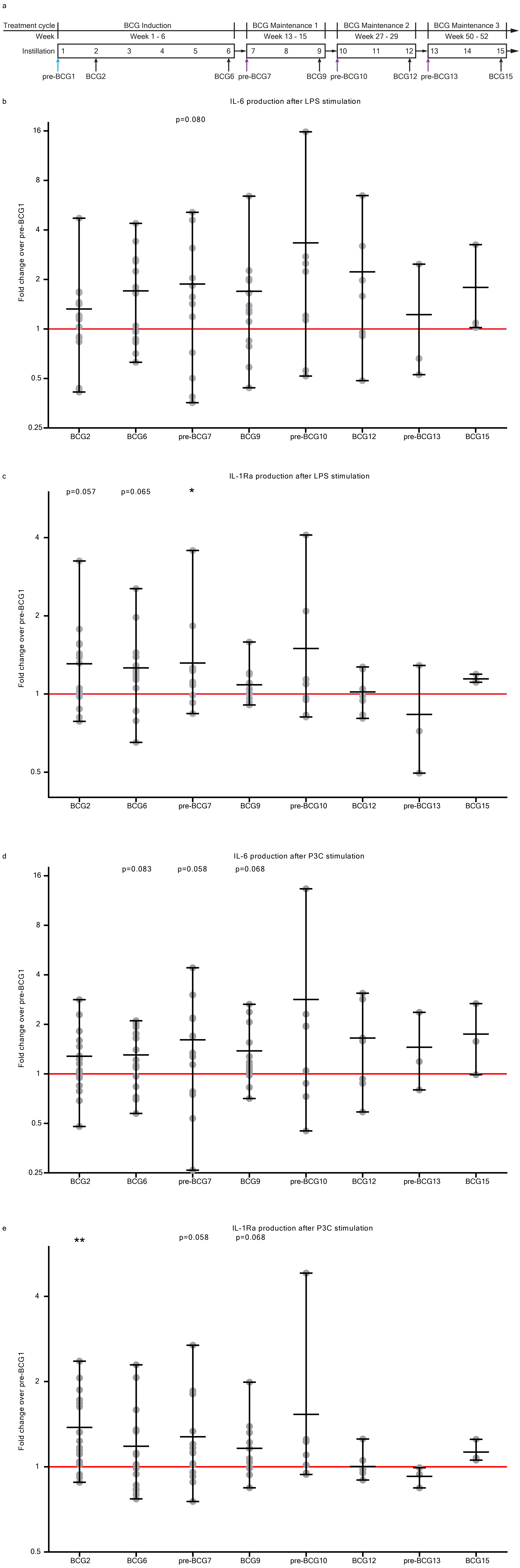

Supplementary Figure 2

a

differentially expressed genes between pre-BCG1 and BCG6

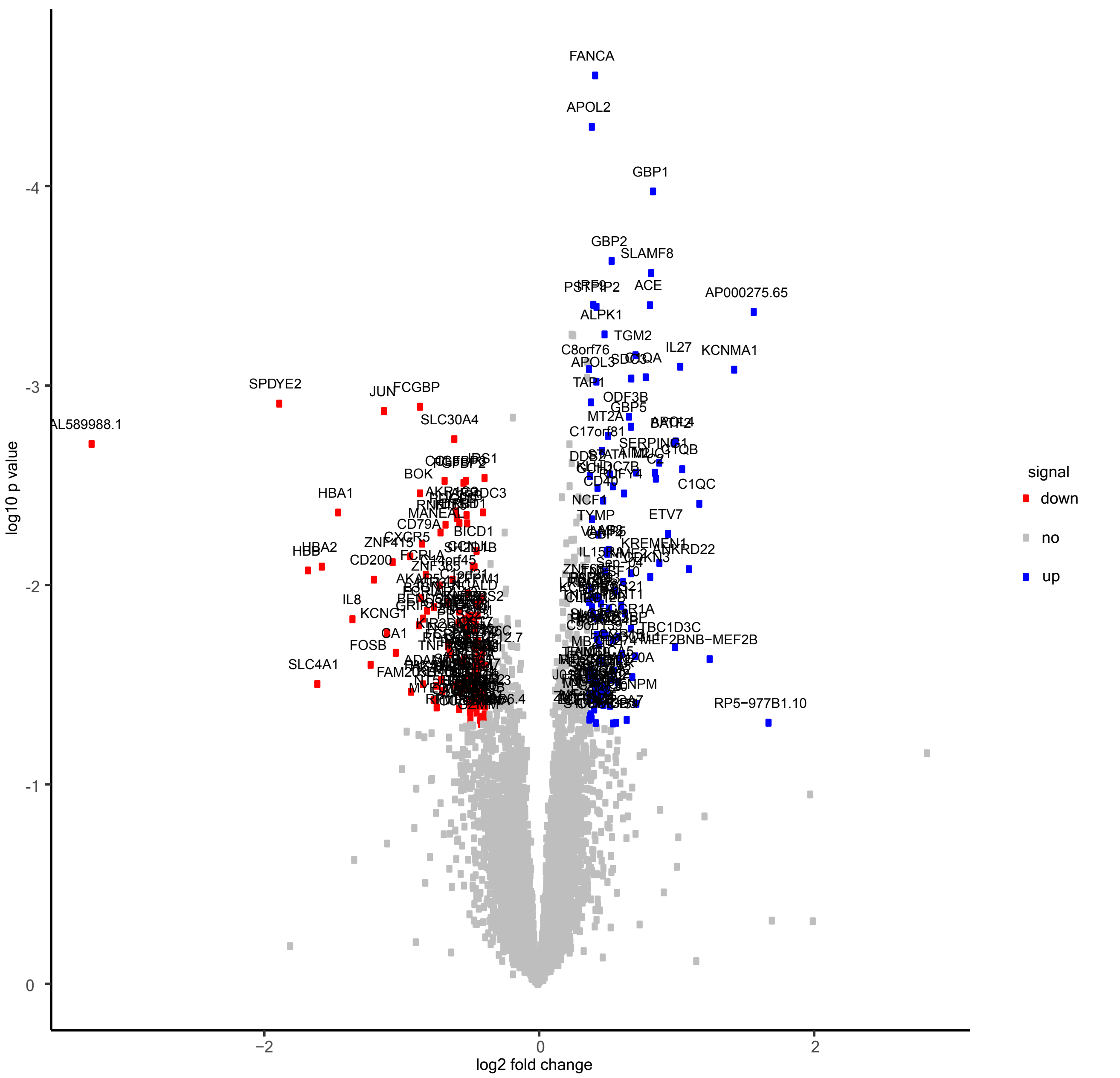

b

differentially expressed genes between pre-BCG1 and pre-BCG7

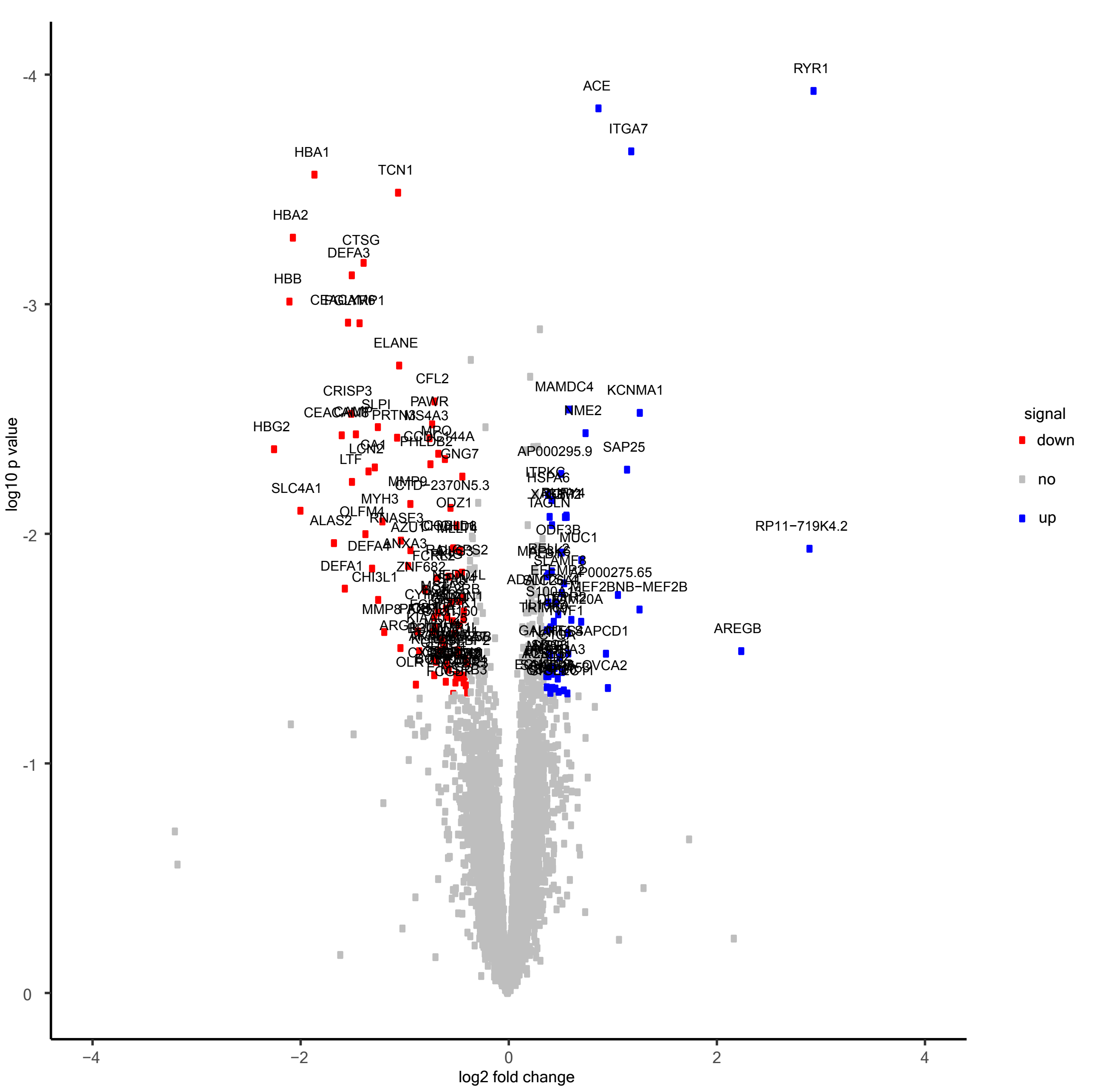

**Supplementary Table 1 - Inclusion and exclusion criteria for the Tribute study.**

| |
|---|
| **Inclusion criteria** |
| 1. Presence of high-grade Ta or T1 urothelial carcinoma of the bladder with or without CIS; tumor can be primary or recurrent |
| 2. Able to communicate in Dutch, and read and understand the patient information and informed consent form and fill out questionnaires |
| 3. Complete resection of (visible) tumor at start of BCG therapy as determined via re-TURT or negative cystoscopy/cytology (at most 6 weeks prior to BCG therapy initiation) |
| **Exclusion critera** |
| 1. Any previous intravesical BCG therapy |
| 2. Presence of primary CIS only |
| 3. Presence of histopathologically proven muscle-invasive urothelial carcinoma of the bladder at first or re-TUR surgical specimens |
| 4. Presence of tumor stage cN1 of cM1 as assessed by CT-thorax/abdomen |
| 5. Presence of any upper urinary tract tumors |
| 6. Histology subtype of resected tumor is not predominantly urothelial carcinoma |
| 7. Presence of another malignancy other than basal cell carcinoma of the skin or prostate cancer under active surveillance |
| 8. Presence of pregnancy or lactation |
| 9. Presence of active tuberculosis, any form of immunodeficiency (e.g. HIV + serology, transplant recipients) and/or any other contraindication of BCG therapy |
| 10. Patients who have received any systemic cytostatic agents within the last 3 months |
| 11. Patients younger than 18 and older than 90 years of age |
| 12. Patients with uncontrollable urinary tract infection |
| 13. Patients who are deemed unfit for study participation as assessed by their urologist |
| 14. Patients who receive any of the following: Methotrexate, Rituximab, Infliximab, Etanercept, other Anti-TNFα therapies, Anti IL-6 therapy, Anti IL-17 therapy or IFN-γ therapy |

**Supplementary Table 2 - Characteristics of patients included in the Tribute study.**

| **Gender** | ***n (%)*** |
|---|---|
| Male | 16 (94) |
| Female | 1 (6) |
| **Age at inclusion (median ± range)** | 67 ± 37 years |
| **Highest tumor stage at study inclusion[1]** | ***n (%)*** |
| Ta | 7 (41) |
| T1 | 10 (59) |
| **Highest tumor grade at study inclusion[1]** | ***n (%)*** |
| Grade 1 | 0 (0) |
| Grade 2a | 0 (0) |
| Grade 2b | 2 (12) |
| Grade 3 | 15 (88) |
| **Concomitant CIS at study inclusion** | ***n (%)*** |
| Yes | 6 (35) |
| No | 11 (65) |
| **Total number of BCG received in the year after start BCG** | ***n (%)*** |
| 5 or 6 | 2 (12) |
| 7 - 9 | 3 (18) |
| 10 - 12 | 5 (29) |
| 13 - 15 | 7 (41) |
| **Recurrence** | ***n (%)*** |
| Yes | 5 (29) |
| No | 12 (71) |
| **Reason for discontinuation with BCG or irregular BCG regimen** | ***n*** |
| Side effects | 4 |
| Tumor progression | 2 |
| BCG shortage | 1 |
| Other | 4 |
| **Clinical follow-up time** | ***months*** |
| Minimum | 21 |
| Maximum | 41 |

[1]The highest tumor stage/grade is reported as the highest stage or grade which was found in the pathology report at the transurethral resection of the bladder tumor (TURT) or re-TURT of study inclusion.

**Supplementary Table 3 - Overview of the concentration of circulating leukocytes in whole blood measured at different time points during BCG therapy in the Tribute study.**

Values are displayed as median ± range. Two-tailed matched-pairs Wilcoxon signed-rank test was used to determine statistical significant difference between time points. Statistical significance was accepted at p<0.05.

| Cell type | Time points and comparisons | | | | | | | | | | | | | | | |
|---|---|---|---|---|---|---|---|---|---|---|---|---|---|---|---|---|
| | pre-BCG1 n=17 | BCG 2 n=17 | p-value BCG2 : Pre-BCG1 | BCG 6 n=16 | p-value BCG 6 : Pre-BCG 1 | pre-BCG 7 n=14 | p-value Pre-BCG7 : Pre-BCG1 | BCG 9 n=12 | p-value BCG 9 : Pre-BCG 1 | pre-BCG 10 n=8 | p-value Pre-BCG10 : Pre-BCG1 | BCG 12 n=7 | p-value BCG12 : Pre-BCG1 | pre-BCG13 n=3 | p-value Pre-BCG13 : Pre-BCG1 | BCG 15 n=3 | p-value BCG15 : Pre-BCG1 |
| White blood cells (x10^9 /L) | 6.44 ± 6.66 | 7.15 ± 6.87 | **p<0.01** | 6.71 ± 6.29 | 0.35 | 6.98 ± 7.01 | 0.15 | 7.09 ± 6.72 | 0.42 | 7.10 ± 10.56 | 0.15 | 8.47 ± 6.30 | 0.08 | 6.63 ± 1.21 | 0.50 | 7.17 ± 2.33 | 0.50 |
| Neutrophils (x10^9 /L) | 3.78 ± 5.27 | 4.18 ± 5.81 | **p<0.01** | 4.18 ± 5.53 | 0.10 | 3.88 ± 6.72 | 0.28 | 4.04 ± 6.08 | 0.09 | 3.99 ± 9.29 | 0.15 | 5.8 ± 5.53 | 0.08 | 3.56 ± 0.80 | 0.75 | 4.33 ± 1.32 | 0.50 |
| Monocytes (x10^9 /L) | 0.64 ± 0.63 | 0.75 ± 0.75 | **p<0.01** | 0.68 ± 0.74 | 0.70 | 0.68 ± 0.64 | 0.15 | 0.72 ± 0.49 | 0.31 | 0.655 ± 1.57 | 0.64 | 0.65 ± 0.20 | 0.89 | 0.65 ± 0.06 | 0.75 | 0.73 ± 0.32 | 0.50 |
| Lymphocytes (x10^9 /L) | 1.98 ± 2.71 | 1.89 ± 3.10 | 0.35 | 1.89 ± 2.62 | 0.13 | 2.02 ± 2.52 | 0.73 | 1.67 ± 1.79 | 0.15 | 1.89 ± 0.90 | p>0.99 | 1.86 ± 0.82 | **p<0.05** | 2.47 ± 1.02 | 0.25 | 2.03 ± 0.85 | 0.25 |

**Supplementary Table 4 - Results from the ChIP-seq analysis.**

Comparisons in H3K4me3 signal were made between pre-BCG1 and any of the measured post-BCG time points (i.e., BCG6, pre-BCG7, BCG9 and pre-BCG10).

Candidate trained immunity genes are indicated in the final column.

| Genomic location | Comparison (up/down in H3K4me3 signal) | log2 fold change | p value | adjusted p value | Distance to gene (in bp) | Nearest Gene | Candidate trained immunity gene |
|---|---|---|---|---|---|---|---|
| chr10_77636776_77638793 | pre-BCG1_v_BCG6_up | 0,689051403 | 0,007152971 | 0,998975358 | 95266 | C10orf11 | |
| chr11_111512299_111512995 | pre-BCG1_v_BCG6_up | 0,654214908 | 0,002777022 | 0,998975358 | 39532 | SIK2 | |
| chr16_57416786_57417262 | pre-BCG1_v_BCG6_up | 0,738508555 | 0,01150535 | 0,998975358 | 10654 | CX3CL1 | |
| chr17_44911270_44911728 | pre-BCG1_v_BCG6_up | 0,804794584 | 0,005248445 | 0,998975358 | -15373 | WNT3 | |
| chr17_63848797_63849266 | pre-BCG1_v_BCG6_up | 1,004453929 | 0,010607428 | 0,998975358 | -291267 | AXIN2 | |
| chr17_68599032_68601980 | pre-BCG1_v_BCG6_up | 0,595314616 | 0,046093502 | 0,998975358 | 434830 | KCNJ2 | |
| chr19_44808740_44809365 | pre-BCG1_v_BCG6_up | 0,590164057 | 0,031538809 | 0,998975358 | 125 | ZNF235 | |
| chr2_218065225_218065862 | pre-BCG1_v_BCG6_up | 0,743442953 | 0,022465992 | 0,998975358 | -340757 | TNP1 | |
| chr2_38535872_38536457 | pre-BCG1_v_BCG6_up | 0,602418985 | 0,005357174 | 0,998975358 | 68239 | ATL2 | |
| chr2_47369090_47369828 | pre-BCG1_v_BCG6_up | 0,624761536 | 0,019328058 | 0,998975358 | 13058 | C2orf61 | |
| chr3_127915089_127915604 | pre-BCG1_v_BCG6_up | 0,755085202 | 0,011702195 | 0,998975358 | 43050 | EEFSEC | |
| chr3_9916678_9917495 | pre-BCG1_v_BCG6_up | 0,694519145 | 0,00221225 | 0,998975358 | 3108 | CIDEC | |
| chr7_6526535_6527062 | pre-BCG1_v_BCG6_up | 0,671134314 | 0,005140945 | 0,998975358 | -2926 | KDELR2 | |
| chr8_75407155_75407974 | pre-BCG1_v_BCG6_up | 0,582119322 | 0,002380173 | 0,998975358 | 144948 | GDAP1 | |
| chr8_76677782_76678320 | pre-BCG1_v_BCG6_up | 0,6499125 | 0,010108019 | 0,998975358 | 225947 | HNF4G | |
| chrX_25012752_25013765 | pre-BCG1_v_BCG6_up | 0,613390269 | 0,002318506 | 0,998975358 | 20806 | ARX | |
| chr1_26786255_26786966 | pre-BCG1_v_BCG6_down | -0,758714745 | 0,000637101 | 0,998975358 | -12399 | HMGN2 | |
| chr1_40388717_40389694 | pre-BCG1_v_BCG6_down | -0,770655267 | 0,004079169 | 0,998975358 | -21521 | MYCL | |
| chr10_15169051_15169800 | pre-BCG1_v_BCG6_down | -0,760960308 | 0,012395145 | 0,998975358 | 30045 | RPP38 | |
| chr10_8060651_8061174 | pre-BCG1_v_BCG6_down | -0,62168621 | 0,019991692 | 0,998975358 | -35743 | GATA3 | |
| chr10_96270807_96271460 | pre-BCG1_v_BCG6_down | -0,746021321 | 0,00105717 | 0,998975358 | -34440 | HELLS | |
| chr11_2988806_2989632 | pre-BCG1_v_BCG6_down | -0,651435773 | 0,024984251 | 0,998975358 | 24388 | NAP1L4 | |
| chr12_123075234_123076132 | pre-BCG1_v_BCG6_down | -0,587435259 | 0,024800866 | 0,998975358 | 63888 | KNTC1 | |
| chr12_48918053_48918675 | pre-BCG1_v_BCG6_down | -0,666184082 | 0,038377299 | 0,998975358 | -1051 | OR8S1 | |
| chr12_9675023_9675706 | pre-BCG1_v_BCG6_down | -0,789100749 | 0,025203784 | 0,998975358 | 85117 | KLRB1 | |

| | | | | | | |
|---|---|---|---|---|---|---|
| chr13_40984594_40985410 | pre-BCG1_v_BCG6_down | -0,60454567 | 0,011782629 | 0,998975358 | 255732 | FOXO1 |
| chr13_99215373_99216071 | pre-BCG1_v_BCG6_down | -0,752790041 | 0,005675127 | 0,998975358 | -41470 | STK24 |
| chr16_83805581_83806134 | pre-BCG1_v_BCG6_down | -0,602580499 | 0,038564865 | 0,998975358 | -35590 | HSBP1 |
| chr17_18186002_18186641 | pre-BCG1_v_BCG6_down | -0,772853983 | 0,012562824 | 0,998975358 | 21898 | MIEF2 |
| chr17_47737581_47738271 | pre-BCG1_v_BCG6_down | -0,618225314 | 0,048005259 | 0,998975358 | 17599 | SPOP |
| chr18_32771283_32771916 | pre-BCG1_v_BCG6_down | -0,666942798 | 0,042069708 | 0,998975358 | -49401 | ZNF397 |
| chr19_14583197_14584356 | pre-BCG1_v_BCG6_down | -0,639228815 | 0,04487689 | 0,998975358 | 2397 | PTGER1 |
| chr19_34391821_34392175 | pre-BCG1_v_BCG6_down | -1,440010467 | 0,012998197 | 0,998975358 | 102707 | KCTD15 |
| chr19_50418781_50420147 | pre-BCG1_v_BCG6_down | -0,726567574 | 0,000165907 | 0,998975358 | -12936 | ATF5 |
| chr19_6996859_6998063 | pre-BCG1_v_BCG6_down | -0,739060745 | 0,007235638 | 0,998975358 | -33128 | MBD3L5 |
| chr19_9362846_9363386 | pre-BCG1_v_BCG6_down | -0,756396915 | 0,02165982 | 0,998975358 | 1510 | OR7E24 |
| chr2_100141611_100142647 | pre-BCG1_v_BCG6_down | -0,583964804 | 0,012887171 | 0,998975358 | -35632 | REV1 |
| chr2_102419197_102419709 | pre-BCG1_v_BCG6_down | -0,617440627 | 0,014536048 | 0,998975358 | 104910 | MAP4K4 |
| chr2_143247185_143247952 | pre-BCG1_v_BCG6_down | -0,680016677 | 0,014482225 | 0,998975358 | -358299 | LRP1B |
| chr2_190359957_190360441 | pre-BCG1_v_BCG6_down | -0,585789115 | 0,037226339 | 0,998975358 | 54040 | WDR75 |
| chr20_44969409_44970473 | pre-BCG1_v_BCG6_down | -0,630627511 | 0,011993377 | 0,998975358 | 21872 | SLC35C2 |
| chr22_30940258_30940863 | pre-BCG1_v_BCG6_down | -0,769205896 | 0,02897983 | 0,998975358 | 2108 | SEC14L6 |
| chr22_31288619_31289191 | pre-BCG1_v_BCG6_down | -1,130939237 | 0,001578769 | 0,998975358 | 75282 | MORC2 |
| chr22_50469375_50470019 | pre-BCG1_v_BCG6_down | -0,661225179 | 0,036962161 | 0,998975358 | -18609 | IL17REL |
| chr3_108125146_108126412 | pre-BCG1_v_BCG6_down | -0,618515416 | 0,006136101 | 0,998975358 | 104447 | HHLA2 |
| chr3_195540047_195540679 | pre-BCG1_v_BCG6_down | -1,119650598 | 0,001262186 | 0,998975358 | -1215 | MUC4 |
| chr3_28292893_28294004 | pre-BCG1_v_BCG6_down | -0,592031587 | 0,01612779 | 0,998975358 | 10363 | CMC1 |
| chr4_69759047_69759990 | pre-BCG1_v_BCG6_down | -0,705055899 | 0,012000652 | 0,998975358 | 57990 | UGT2A3 |
| chr5_119333328_119334023 | pre-BCG1_v_BCG6_down | -0,673871289 | 0,029384374 | 0,998975358 | 368383 | FAM170A |
| chr5_138369920_138370524 | pre-BCG1_v_BCG6_down | -0,899239115 | 0,004222722 | 0,998975358 | -159243 | LRRTM2 |
| chr5_150846511_150847193 | pre-BCG1_v_BCG6_down | -0,716839438 | 0,002174625 | 0,998975358 | 19695 | SLC36A1 |
| chr5_56159746_56160675 | pre-BCG1_v_BCG6_down | -0,615806547 | 0,041983077 | 0,998975358 | -44876 | SETD9 |
| chr6_127899285_127901264 | pre-BCG1_v_BCG6_down | -0,624973191 | 0,008861184 | 0,998975358 | 1956 | C6orf58 |
| chr6_139839585_139840271 | pre-BCG1_v_BCG6_down | -0,604446506 | 0,027189014 | 0,998975358 | -144171 | CITED2 |
| chr6_158649979_158651155 | pre-BCG1_v_BCG6_down | -0,695646842 | 0,002252665 | 0,998975358 | 61183 | GTF2H5 |
| chr7_18495755_18496388 | pre-BCG1_v_BCG6_down | -0,816832621 | 0,004153717 | 0,998975358 | -39854 | HDAC9 |
| chr7_38362900_38363577 | pre-BCG1_v_BCG6_down | -0,977123545 | 0,007895706 | 0,998975358 | 145415 | STARD3NL |

| | | | | | | |
|---|---|---|---|---|---|---|
| chr8_11494415_11495264 | pre-BCG1_v_BCG6_down | -0,81473298 | 0,001306633 | 0,998975358 | -66873 | GATA4 |
| chr8_47360890_47361635 | pre-BCG1_v_BCG6_down | -0,749887182 | 0,0038632 | 0,998975358 | -811904 | SPIDR |
| chr9_130038378_130038894 | pre-BCG1_v_BCG6_down | -0,604842598 | 0,036413148 | 0,998975358 | 11831 | GARNL3 |
| chr9_35728125_35729024 | pre-BCG1_v_BCG6_down | -0,624533639 | 0,00549807 | 0,998975358 | -3757 | CREB3 |
| chrX_12909856_12910867 | pre-BCG1_v_BCG6_down | -0,617485179 | 0,012246177 | 0,998975358 | -14377 | TLR8 |
| chrX_136648164_136651743 | pre-BCG1_v_BCG6_down | -0,597976429 | 0,037736277 | 0,998975358 | 1653 | ZIC3 |
| chr1_6440247_6440965 | pre-BCG1_v_pre-BCG7_up | 0,83546548 | 0,002150024 | 0,999994721 | 5267 | ACOT7 |
| chr2_218065225_218065862 | pre-BCG1_v_pre-BCG7_up | 0,868386188 | 0,007333457 | 0,999994721 | -340757 | TNP1 |
| chr9_121736375_121736868 | pre-BCG1_v_pre-BCG7_up | 0,620003543 | 0,031364585 | 0,999994721 | 395123 | BRINP1 |
| chrX_65407078_65407688 | pre-BCG1_v_pre-BCG7_up | 0,63097322 | 0,035991197 | 0,999994721 | 24992 | HEPH |
| chr1_16514553_16514998 | pre-BCG1_v_pre-BCG7_down | -0,580738733 | 0,00848568 | 0,999994721 | 24328 | ARHGEF19 |
| chr1_167439282_167439705 | pre-BCG1_v_pre-BCG7_down | -0,660770461 | 0,038700354 | 0,999994721 | 48281 | CD247 |
| chr1_248404046_248404400 | pre-BCG1_v_pre-BCG7_down | -1,491606368 | 0,039297173 | 0,999994721 | 1992 | OR2M4 |
| chr1_44730816_44731405 | pre-BCG1_v_pre-BCG7_down | -0,617265555 | 0,011010381 | 0,999994721 | 51948 | DMAP1 |
| chr1_47179267_47179652 | pre-BCG1_v_pre-BCG7_down | -0,655516037 | 0,032840975 | 0,999994721 | 5276 | EFCAB14 |
| chr10_69228723_69229856 | pre-BCG1_v_pre-BCG7_down | -0,634107878 | 0,000606902 | 0,999994721 | 226637 | CTNNA3 |
| chr10_96270807_96271460 | pre-BCG1_v_pre-BCG7_down | -0,621199545 | 0,00580907 | 0,999994721 | -34440 | HELLS |
| chr10_96699935_96700761 | pre-BCG1_v_pre-BCG7_down | -0,665161093 | 0,018604444 | 0,999994721 | 1920 | CYP2C9 |
| chr12_131818687_131819254 | pre-BCG1_v_pre-BCG7_down | -0,597698825 | 0,026005192 | 0,999994721 | -376711 | SFSWAP |
| chr14_102544516_102545186 | pre-BCG1_v_pre-BCG7_down | -0,607150187 | 0,035060911 | 0,999994721 | 61172 | HSP90AA1 |
| chr14_23118617_23119253 | pre-BCG1_v_pre-BCG7_down | -0,666449245 | 0,03024756 | 0,999994721 | -15219 | OR6J1 |
| chr15_57300317_57302218 | pre-BCG1_v_pre-BCG7_down | -0,697848958 | 0,005520165 | 0,999994721 | 90447 | TCF12 |
| chr16_87490634_87491479 | pre-BCG1_v_pre-BCG7_down | -0,60412216 | 0,043686052 | 0,999994721 | 34594 | ZCCHC14 |
| chr17_18186002_18186641 | pre-BCG1_v_pre-BCG7_down | -0,680603968 | 0,02686072 | 0,999994721 | 21898 | MIEF2 |
| chr17_38967117_38967721 | pre-BCG1_v_pre-BCG7_down | -0,725315035 | 0,004828794 | 0,999994721 | -7958 | TMEM99 |
| chr17_60102370_60102793 | pre-BCG1_v_pre-BCG7_down | -0,613147573 | 0,017707941 | 0,999994721 | 40061 | MED13 |
| chr17_7838486_7839742 | pre-BCG1_v_pre-BCG7_down | -0,755766647 | 9,27E-05 | 0,549122361 | 3641 | CNTROB |
| chr19_14583197_14584356 | pre-BCG1_v_pre-BCG7_down | -0,646723082 | 0,0424442 | 0,999994721 | 2397 | PTGER1 |
| chr19_34391316_34391731 | pre-BCG1_v_pre-BCG7_down | -1,191724455 | 0,017510171 | 0,999994721 | 102233 | KCTD15 |
| chr19_34391821_34392175 | pre-BCG1_v_pre-BCG7_down | -1,333760569 | 0,020529106 | 0,999994721 | 102707 | KCTD15 |
| chr19_41877729_41878637 | pre-BCG1_v_pre-BCG7_down | -0,724618447 | 0,001746396 | 0,999994721 | -3946 | TMEM91 |
| chr19_50418781_50420147 | pre-BCG1_v_pre-BCG7_down | -0,866652017 | 7,73E-06 | 0,150338015 | -12936 | ATF5 |

| | | | | | | |
|---|---|---|---|---|---|---|
| chr19_7450321_7450984 | pre-BCG1_v_pre-BCG7_down | -1,133122444 | 0,027813742 | 0,999994721 | 36805 | ENSG00000263264 |
| chr19_9362846_9363386 | pre-BCG1_v_pre-BCG7_down | -0,64667456 | 0,047818914 | 0,999994721 | 1510 | OR7E24 |
| chr2_132416089_132416758 | pre-BCG1_v_pre-BCG7_down | -0,671188751 | 0,014675765 | 0,999994721 | -166429 | MZT2A |
| chr2_38600354_38601147 | pre-BCG1_v_pre-BCG7_down | -0,659593304 | 0,011498059 | 0,999994721 | 3653 | ATL2 |
| chr20_45097972_45098504 | pre-BCG1_v_pre-BCG7_down | -0,644586356 | 0,029180731 | 0,999994721 | 43955 | ZNF334 |
| chr20_60058430_60058967 | pre-BCG1_v_pre-BCG7_down | -0,589917859 | 0,029753883 | 0,999994721 | 231217 | CDH4 |
| chr22_10739623_10740127 | pre-BCG1_v_pre-BCG7_down | -1,214844819 | 0,018147771 | 0,999994721 | NONE | NONE |
| chr3_195540047_195540679 | pre-BCG1_v_pre-BCG7_down | -0,857411093 | 0,012256027 | 0,999994721 | -1215 | MUC4 |
| chr3_43996868_43997375 | pre-BCG1_v_pre-BCG7_down | -0,820227974 | 0,002891076 | 0,999994721 | 264760 | ABHD5 |
| chr4_114598892_114599748 | pre-BCG1_v_pre-BCG7_down | -0,956775355 | 0,000252984 | 0,81954027 | 82904 | CAMK2D |
| chr6_38715073_38715534 | pre-BCG1_v_pre-BCG7_down | -0,733232054 | 0,006110365 | 0,999994721 | 24406 | DNAH8 |
| chr8_24293961_24294531 | pre-BCG1_v_pre-BCG7_down | -0,603682306 | 0,031733059 | 0,999994721 | -4293 | ADAM7 |
| chr9_130038378_130038894 | pre-BCG1_v_pre-BCG7_down | -0,642805347 | 0,025783826 | 0,999994721 | 11831 | GARNL3 |
| chr9_35728125_35729024 | pre-BCG1_v_pre-BCG7_down | -0,674004515 | 0,002669947 | 0,999994721 | -3757 | CREB3 |
| chr9_41229458_41230737 | pre-BCG1_v_pre-BCG7_down | -1,060673596 | 0,043559406 | 0,999994721 | 97266 | SPATA31A4 |
| chr1_157993429_157994057 | pre-BCG1_v_BCG9_up | 0,696914991 | 0,002680689 | 0,242014136 | 30343 | KIRREL |
| chr1_16623844_16624356 | pre-BCG1_v_BCG9_up | 1,466729193 | 0,011485026 | 0,327924731 | 54849 | FBXO42 |
| chr1_16725115_16725553 | pre-BCG1_v_BCG9_up | 1,303118063 | 0,038490456 | 0,433902916 | 38585 | SPATA21 |
| chr1_167439282_167439705 | pre-BCG1_v_BCG9_up | 0,669323257 | 0,029654949 | 0,406580095 | 48281 | CD247 |
| chr1_247517760_247518647 | pre-BCG1_v_BCG9_up | 0,64328358 | 0,003111115 | 0,247622157 | -23159 | ZNF496 |
| chr1_50023617_50024078 | pre-BCG1_v_BCG9_up | 0,809239495 | 0,004900496 | 0,26789583 | 465737 | AGBL4 |
| chr1_52368383_52368812 | pre-BCG1_v_BCG9_up | 0,714817508 | 0,01140199 | 0,326392462 | -24121 | NRD1 |
| chr1_6440247_6440965 | pre-BCG1_v_BCG9_up | 0,661925934 | 0,017121536 | 0,357131651 | 5267 | ACOT7 |
| chr10_18140266_18140963 | pre-BCG1_v_BCG9_up | 0,696448971 | 0,014760934 | 0,342969644 | 42263 | MRC1 |
| chr10_48523908_48524321 | pre-BCG1_v_BCG9_up | 0,884174765 | 0,007464466 | 0,291925216 | -85139 | GDF10 |
| chr10_51074164_51075050 | pre-BCG1_v_BCG9_up | 1,070598112 | 0,001037404 | 0,194978227 | 56108 | PARG |
| chr10_69408912_69409384 | pre-BCG1_v_BCG9_up | 0,604722225 | 0,005301692 | 0,26789583 | 46779 | CTNNA3 |
| chr10_76163_76791 | pre-BCG1_v_BCG9_up | 0,597805161 | 0,001690453 | 0,206649859 | 18764 | TUBB8 |
| chr10_77636776_77638793 | pre-BCG1_v_BCG9_up | 0,675603544 | 0,008593409 | 0,303691088 | 95266 | C10orf11 |
| chr10_93600817_93601603 | pre-BCG1_v_BCG9_up | 0,720275938 | 0,011776204 | 0,327924731 | 43141 | TNKS2 |
| chr10_96374986_96375491 | pre-BCG1_v_BCG9_up | 0,611816243 | 0,00999409 | 0,31586199 | -68139 | CYP2C18 |
| chr10_99536791_99537421 | pre-BCG1_v_BCG9_up | 0,64714305 | 0,047965626 | 0,462208279 | -5397 | SFRP5 |

| | | | | | | | |
|---|---|---|---|---|---|---|---|
| chr11_129375636_129376399 | pre-BCG1_v_BCG9_up | 0,809249811 | 0,039057075 | 0,436127427 | 130183 | BARX2 | |
| chr11_57311205_57311772 | pre-BCG1_v_BCG9_up | 0,669137502 | 0,0044124 | 0,264296386 | 2510 | SMTNL1 | |
| chr11_9091129_9091919 | pre-BCG1_v_BCG9_up | 0,593183795 | 0,01092102 | 0,324090769 | 21622 | SCUBE2 | |
| chr12_108129230_108129631 | pre-BCG1_v_BCG9_up | 0,876948204 | 0,001463556 | 0,20205633 | 25506 | PRDM4 | |
| chr12_120368678_120369627 | pre-BCG1_v_BCG9_up | 0,687052935 | 0,020597858 | 0,375283061 | -54058 | CIT | |
| chr12_203473_203985 | pre-BCG1_v_BCG9_up | 0,881814335 | 0,001923152 | 0,219884136 | 27798 | IQSEC3 | |
| chr12_33439120_33439771 | pre-BCG1_v_BCG9_up | 0,738685843 | 0,01185274 | 0,328647216 | 153308 | SYT10 | |
| chr12_553478_554415 | pre-BCG1_v_BCG9_up | 1,07821576 | 0,020982764 | 0,375283061 | -15583 | B4GALNT3 | |
| chr12_9965209_9966762 | pre-BCG1_v_BCG9_up | 0,582169183 | 0,026797007 | 0,391619113 | -14091 | KLRF1 | |
| chr13_36431427_36432649 | pre-BCG1_v_BCG9_up | 0,621170094 | 0,000491653 | 0,15927111 | 273405 | DCLK1 | |
| chr13_36919663_36920530 | pre-BCG1_v_BCG9_up | 0,604666747 | 0,004740503 | 0,26789583 | 837 | SPG20 | |
| chr14_24314661_24315492 | pre-BCG1_v_BCG9_up | 0,654575787 | 0,015510165 | 0,34738072 | -107718 | DHRS4 | |
| chr15_20282402_20282955 | pre-BCG1_v_BCG9_up | 1,520223688 | 0,02040073 | 0,374620679 | NONE | NONE | |
| chr15_20348867_20349764 | pre-BCG1_v_BCG9_up | 1,46919324 | 0,010630255 | 0,32138843 | NONE | NONE | |
| chr15_51341595_51342117 | pre-BCG1_v_BCG9_up | 0,631708352 | 0,049777963 | 0,465008002 | 55617 | TNFAIP8L3 | |
| chr15_67825416_67825995 | pre-BCG1_v_BCG9_up | 0,593717592 | 0,020727804 | 0,375283061 | -9341 | MAP2K5 | |
| chr15_70853648_70854381 | pre-BCG1_v_BCG9_up | 0,787537646 | 0,003766888 | 0,264101006 | 201917 | UACA | |
| chr15_93072319_93073342 | pre-BCG1_v_BCG9_up | 0,608904603 | 0,023870548 | 0,381242269 | 126357 | FAM174B | |
| chr16_28505586_28508309 | pre-BCG1_v_BCG9_up | 0,621971477 | 0,02243421 | 0,375911088 | -93 | CLN3 | |
| chr16_3112406_3113051 | pre-BCG1_v_BCG9_up | 0,716089086 | 0,002706427 | 0,24241855 | -2656 | IL32 | Yes |
| chr16_68356168_68356928 | pre-BCG1_v_BCG9_up | 0,621402042 | 0,042897326 | 0,447569633 | 11511 | PRMT7 | |
| chr16_83968176_83968703 | pre-BCG1_v_BCG9_up | 0,605588733 | 0,023939651 | 0,381482232 | -14232 | OSGIN1 | |
| chr17_32927753_32928522 | pre-BCG1_v_BCG9_up | 0,800033905 | 0,004281337 | 0,264296386 | 20370 | TMEM132E | |
| chr17_43231538_43232635 | pre-BCG1_v_BCG9_up | 0,824499503 | 0,015605312 | 0,34738072 | -5980 | HEXIM2 | |
| chr17_44308153_44309186 | pre-BCG1_v_BCG9_up | 0,730001858 | 0,002421425 | 0,237208728 | -38504 | KANSL1 | |
| chr17_44911270_44911728 | pre-BCG1_v_BCG9_up | 0,867666567 | 0,003057146 | 0,247622157 | -15373 | WNT3 | |
| chr17_48576639_48577165 | pre-BCG1_v_BCG9_up | 0,785418302 | 0,025111638 | 0,384932891 | -8843 | MYCBPAP | |
| chr17_54900444_54900960 | pre-BCG1_v_BCG9_up | 0,774179569 | 0,003996913 | 0,264101006 | 10555 | C17orf67 | |
| chr17_57833179_57833561 | pre-BCG1_v_BCG9_up | 0,614435985 | 0,014846127 | 0,34352877 | 48544 | VMP1 | |
| chr17_74861124_74861929 | pre-BCG1_v_BCG9_up | 0,692798883 | 0,025832308 | 0,387850306 | -7202 | MGAT5B | |
| chr18_12777022_12777914 | pre-BCG1_v_BCG9_up | 0,59774619 | 0,025862555 | 0,387850306 | -74695 | CEP76 | |
| chr18_28176755_28178224 | pre-BCG1_v_BCG9_up | 0,591838187 | 0,00145541 | 0,20205633 | 445217 | DSC3 | |

| | | | | | | |
|---|---|---|---|---|---|---|
| chr18_5890697_5891242 | pre-BCG1_v_BCG9_up | 0,7488732 | 0,012392256 | 0,333977084 | 4984 | TMEM200C |
| chr19_1000174_1000965 | pre-BCG1_v_BCG9_up | 0,915505304 | 1,95E-05 | 0,037885069 | 152 | GRIN3B |
| chr19_17794772_17795540 | pre-BCG1_v_BCG9_up | 0,765614172 | 0,022517013 | 0,37611534 | 3852 | UNC13A |
| chr19_18789874_18790435 | pre-BCG1_v_BCG9_up | 0,655269743 | 0,046525222 | 0,457647131 | -4333 | CRTC1 |
| chr19_19224765_19225533 | pre-BCG1_v_BCG9_up | 0,591169806 | 0,003067576 | 0,247622157 | 24153 | TMEM161A |
| chr19_22421606_22422542 | pre-BCG1_v_BCG9_up | 0,929597002 | 0,033701486 | 0,422463275 | -42321 | ZNF676 |
| chr19_2720891_2721772 | pre-BCG1_v_BCG9_up | 0,6207245 | 0,001838641 | 0,212724142 | 84 | DIRAS1 |
| chr19_35040686_35041398 | pre-BCG1_v_BCG9_up | 0,628484742 | 0,009191411 | 0,308925674 | 68499 | WTIP |
| chr19_3576422_3577238 | pre-BCG1_v_BCG9_up | 0,72403725 | 0,001309749 | 0,20205633 | 3894 | HMG20B |
| chr19_37817138_37817866 | pre-BCG1_v_BCG9_up | 0,806825085 | 0,024553392 | 0,384016298 | -8078 | HKR1 |
| chr19_44808740_44809365 | pre-BCG1_v_BCG9_up | 0,628527922 | 0,024096132 | 0,382253164 | 125 | ZNF235 |
| chr19_53979761_53980566 | pre-BCG1_v_BCG9_up | 0,596906904 | 0,023298467 | 0,380061443 | 9175 | ZNF813 |
| chr19_53982086_53983094 | pre-BCG1_v_BCG9_up | 0,5812511 | 0,001131143 | 0,201706747 | 11601 | ZNF813 |
| chr19_7917319_7918073 | pre-BCG1_v_BCG9_up | 0,590539367 | 0,019537726 | 0,372673973 | 6267 | EVI5L |
| chr2_127422914_127423866 | pre-BCG1_v_BCG9_up | 0,62860734 | 0,006245432 | 0,275891965 | 9881 | GYPC |
| chr2_127674657_127675223 | pre-BCG1_v_BCG9_up | 0,581170943 | 0,034744697 | 0,426985488 | 189991 | BIN1 |
| chr2_143936841_143937586 | pre-BCG1_v_BCG9_up | 0,65935479 | 0,015524896 | 0,34738072 | 50331 | ARHGAP15 |
| chr2_149038517_149039089 | pre-BCG1_v_BCG9_up | 0,726993363 | 0,01559817 | 0,34738072 | 260223 | MBD5 |
| chr2_176122575_176123082 | pre-BCG1_v_BCG9_up | 0,765413875 | 0,000875083 | 0,183834761 | -73494 | ATP5G3 |
| chr2_207681283_207681814 | pre-BCG1_v_BCG9_up | 0,756932957 | 0,048970717 | 0,464936962 | -51278 | MDH1B |
| chr2_23386424_23387052 | pre-BCG1_v_BCG9_up | 0,709027327 | 0,032961634 | 0,420390092 | -221350 | KLHL29 |
| chr2_45169015_45169979 | pre-BCG1_v_BCG9_up | 0,590456205 | 0,046787605 | 0,457781598 | 595 | SIX3 |
| chr2_66445405_66446591 | pre-BCG1_v_BCG9_up | 0,62664198 | 0,005421537 | 0,26789583 | -216534 | MEIS1 |
| chr2_88452012_88452886 | pre-BCG1_v_BCG9_up | 0,643657594 | 0,010211674 | 0,320327238 | -18530 | THNSL2 |
| chr20_32031166_32031739 | pre-BCG1_v_BCG9_up | 0,702688365 | 0,015524423 | 0,34738072 | 245 | SNTA1 |
| chr20_6122549_6123226 | pre-BCG1_v_BCG9_up | 0,655804028 | 0,001597815 | 0,204320535 | -18697 | FERMT1 |
| chr21_10327772_10328209 | pre-BCG1_v_BCG9_up | 1,072454777 | 0,026678946 | 0,391466617 | 662891 | TPTE |
| chr21_10355286_10355883 | pre-BCG1_v_BCG9_up | 1,375486898 | 0,013706235 | 0,335128206 | 635297 | TPTE |
| chr21_14063558_14064997 | pre-BCG1_v_BCG9_up | 1,315132933 | 0,030025485 | 0,407545632 | -918220 | POTED |
| chr21_33027133_33027801 | pre-BCG1_v_BCG9_up | 0,662908488 | 0,047562153 | 0,461310162 | -4468 | SOD1 |
| chr21_41316555_41317477 | pre-BCG1_v_BCG9_up | 1,089057429 | 0,000832501 | 0,183834761 | 77773 | PCP4 |
| chr21_8234012_8234860 | pre-BCG1_v_BCG9_up | 1,124627571 | 0,031807823 | 0,415644346 | NONE | NONE |

| | | | | | | |
|---|---|---|---|---|---|---|
| chr22_10731220_10731692 | pre-BCG1_v_BCG9_up | 1,596988656 | 0,011732325 | 0,327924731 | NONE | NONE |
| chr22_19760571_19761219 | pre-BCG1_v_BCG9_up | 0,604852454 | 0,017920949 | 0,363081267 | 16669 | TBX1 |
| chr22_29314846_29315812 | pre-BCG1_v_BCG9_up | 0,627719164 | 0,005893984 | 0,270438199 | 35749 | ZNRF3 |
| chr3_127915089_127915604 | pre-BCG1_v_BCG9_up | 0,901281312 | 0,00289325 | 0,244645504 | 43050 | EEFSEC |
| chr3_129343844_129344505 | pre-BCG1_v_BCG9_up | 0,587395417 | 0,020757667 | 0,375283061 | -18514 | PLXND1 |
| chr3_130093960_130094722 | pre-BCG1_v_BCG9_up | 0,727586564 | 0,002347473 | 0,237208728 | 29982 | COL6A5 |
| chr3_134374020_134374815 | pre-BCG1_v_BCG9_up | 0,607017141 | 0,023845731 | 0,381159114 | -4554 | KY |
| chr3_139934997_139935443 | pre-BCG1_v_BCG9_up | 0,642974065 | 0,024902901 | 0,384040277 | 281193 | CLSTN2 |
| chr3_141051071_141051620 | pre-BCG1_v_BCG9_up | 0,728896553 | 0,023231625 | 0,380061443 | -55308 | ZBTB38 |
| chr3_30894470_30894938 | pre-BCG1_v_BCG9_up | 0,687648774 | 0,00592483 | 0,270438199 | 41553 | GADL1 |
| chr3_43995119_43995603 | pre-BCG1_v_BCG9_up | 0,6645682 | 0,025248709 | 0,386159097 | 262999 | ABHD5 |
| chr3_46565820_46566396 | pre-BCG1_v_BCG9_up | 0,688428293 | 0,015788242 | 0,348936879 | 27127 | RTP3 |
| chr3_53494172_53494923 | pre-BCG1_v_BCG9_up | 0,601742586 | 0,010599033 | 0,32138843 | -34528 | CACNA1D |
| chr3_9916678_9917495 | pre-BCG1_v_BCG9_up | 0,794472756 | 0,000527765 | 0,165454372 | 3108 | CIDEC |
| chr4_118033227_118033849 | pre-BCG1_v_BCG9_up | 0,580366785 | 0,03786915 | 0,433147183 | -26802 | TRAM1L1 |
| chr4_182448678_182449338 | pre-BCG1_v_BCG9_up | 0,620179297 | 0,03795948 | 0,433147183 | -715574 | TENM3 |
| chr4_61200308_61201490 | pre-BCG1_v_BCG9_up | 0,752711427 | 0,000591506 | 0,174952403 | -866961 | LPHN3 |
| chr4_82561577_82562345 | pre-BCG1_v_BCG9_up | 0,676972293 | 0,00140907 | 0,20205633 | -168892 | RASGEF1B |
| chr4_8269566_8270406 | pre-BCG1_v_BCG9_up | 0,586413764 | 0,019317851 | 0,371763444 | -1506 | HTRA3 |
| chr5_120465493_120465806 | pre-BCG1_v_BCG9_up | 0,703946965 | 0,028883124 | 0,40407839 | 665631 | PRR16 |
| chr5_135578638_135579241 | pre-BCG1_v_BCG9_up | 0,581065943 | 0,048632436 | 0,464076546 | 110404 | SMAD5 |
| chr5_147509584_147510161 | pre-BCG1_v_BCG9_up | 0,837101693 | 0,023093127 | 0,380061443 | -39423 | SPINK14 |
| chr5_179343933_179345711 | pre-BCG1_v_BCG9_up | 0,724076599 | 0,0107685 | 0,32400516 | -9963 | TBC1D9B |
| chr5_55176249_55176797 | pre-BCG1_v_BCG9_up | 0,635597392 | 0,045222217 | 0,453852092 | 29189 | IL31RA |
| chr5_88680214_88680682 | pre-BCG1_v_BCG9_up | 0,75692087 | 0,000662442 | 0,180466765 | -480549 | MEF2C |
| chr6_104941012_104941716 | pre-BCG1_v_BCG9_up | 0,628347827 | 0,004613939 | 0,26789583 | 366430 | HACE1 |
| chr6_147508112_147508790 | pre-BCG1_v_BCG9_up | 0,637603366 | 0,04087647 | 0,442075782 | -17218 | STXBP5 |
| chr6_151325412_151325883 | pre-BCG1_v_BCG9_up | 0,651859948 | 0,029581631 | 0,406580095 | 138963 | MTHFD1L |
| chr6_159168949_159170125 | pre-BCG1_v_BCG9_up | 0,722453195 | 0,000644356 | 0,180466765 | 70907 | EZR |
| chr7_102279639_102280821 | pre-BCG1_v_BCG9_up | 0,663762491 | 0,018657892 | 0,369928459 | 3008 | UPK3BL |
| chr7_116671796_116672546 | pre-BCG1_v_BCG9_up | 0,646015598 | 0,001942548 | 0,22080295 | 78790 | ST7 |
| chr7_130491112_130491571 | pre-BCG1_v_BCG9_up | 0,745652916 | 0,006499564 | 0,28068943 | -72454 | KLF14 |

| | | | | | | |
|---|---|---|---|---|---|---|
| chr7_130491941_130492469 | pre-BCG1_v_BCG9_up | 0,84489264 | 0,004879805 | 0,26789583 | -73317 | KLF14 |
| chr7_20786734_20787201 | pre-BCG1_v_BCG9_up | 0,637094634 | 0,030901736 | 0,412221444 | 39537 | SP8 |
| chr7_21542834_21543348 | pre-BCG1_v_BCG9_up | 0,626185715 | 0,003583466 | 0,264101006 | -39742 | DNAH11 |
| chr7_33904448_33905202 | pre-BCG1_v_BCG9_up | 0,643791831 | 0,030802065 | 0,411908982 | -39698 | BMPER |
| chr7_35971340_35971856 | pre-BCG1_v_BCG9_up | 0,645722934 | 0,048458775 | 0,463839356 | 130971 | SEPT7 |
| chr7_38630698_38631640 | pre-BCG1_v_BCG9_up | 0,682129889 | 0,004535333 | 0,266324081 | 39998 | AMPH |
| chr7_64463539_64464431 | pre-BCG1_v_BCG9_up | 0,630761844 | 0,017399399 | 0,358634266 | 3046 | ERV3-1 |
| chr7_6526535_6527062 | pre-BCG1_v_BCG9_up | 0,88590328 | 0,000258877 | 0,10938679 | -2926 | KDELR2 |
| chr7_82443268_82443808 | pre-BCG1_v_BCG9_up | 0,591024602 | 0,044028786 | 0,449946482 | 348708 | PCLO |
| chr8_103501338_103501855 | pre-BCG1_v_BCG9_up | 0,667993108 | 0,037886527 | 0,433147183 | -62203 | ODF1 |
| chr8_141366083_141367236 | pre-BCG1_v_BCG9_up | 0,598766793 | 0,033286019 | 0,420390092 | 102018 | TRAPPC9 |
| chr8_21787326_21788103 | pre-BCG1_v_BCG9_up | 0,589655644 | 0,029141762 | 0,405170543 | 10533 | XPO7 |
| chr8_76677782_76678320 | pre-BCG1_v_BCG9_up | 0,762649013 | 0,002873563 | 0,244645504 | 225947 | HNF4G |
| chr9_113154430_113154996 | pre-BCG1_v_BCG9_up | 0,678113351 | 0,017622434 | 0,360554994 | -54625 | TXNDC8 |
| chr9_121599254_121599823 | pre-BCG1_v_BCG9_up | 0,628292951 | 0,021876158 | 0,375283061 | 532206 | BRINP1 |
| chr9_121736375_121736868 | pre-BCG1_v_BCG9_up | 0,838147983 | 0,004027901 | 0,264296386 | 395123 | BRINP1 |
| chr9_130938798_130939748 | pre-BCG1_v_BCG9_up | 0,605288038 | 0,026837987 | 0,391790751 | 16734 | C9orf16 |
| chr9_137053642_137054370 | pre-BCG1_v_BCG9_up | 0,654073889 | 0,018884275 | 0,369928459 | 52796 | WDR5 |
| chr9_4490073_4491116 | pre-BCG1_v_BCG9_up | 1,041047554 | 0,000687068 | 0,180466765 | 151 | SLC1A1 |
| chr9_8857297_8858141 | pre-BCG1_v_BCG9_up | 0,669988044 | 0,005019625 | 0,26789583 | NONE | NONE |
| chr9_93131329_93131827 | pre-BCG1_v_BCG9_up | 0,817309674 | 0,009981916 | 0,31586199 | 273808 | DIRAS2 |
| chr9_97848484_97849321 | pre-BCG1_v_BCG9_up | 0,631771814 | 0,006825443 | 0,284780812 | 231081 | FANCC |
| chrX_65407078_65407688 | pre-BCG1_v_BCG9_up | 1,29251962 | 1,61E-05 | 0,037885069 | 24992 | HEPH |
| chrY_11307334_11308401 | pre-BCG1_v_BCG9_up | 1,346524165 | 0,014501798 | 0,340895802 | NONE | NONE |
| chrY_11322380_11322990 | pre-BCG1_v_BCG9_up | 1,208813916 | 0,040049872 | 0,440548595 | NONE | NONE |
| chrY_11331928_11334400 | pre-BCG1_v_BCG9_up | 1,110976485 | 0,033935189 | 0,423906348 | NONE | NONE |
| chrY_56850100_56851308 | pre-BCG1_v_BCG9_up | 1,094258712 | 0,028958355 | 0,404355991 | NONE | NONE |
| chrY_56855130_56856102 | pre-BCG1_v_BCG9_up | 1,224352727 | 0,019675992 | 0,374209644 | NONE | NONE |
| chr1_158830992_158837305 | pre-BCG1_v_BCG9_down | -1,005472938 | 0,000826299 | 0,183834761 | 33042 | MNDA |
| chr1_158930786_158934314 | pre-BCG1_v_BCG9_down | -0,888955163 | 0,000867255 | 0,183834761 | 31208 | PYHIN1 |
| chr1_16514553_16514998 | pre-BCG1_v_BCG9_down | -0,686907861 | 0,003234611 | 0,250482609 | 24328 | ARHGEF19 |
| chr1_169584731_169585422 | pre-BCG1_v_BCG9_down | -1,008851354 | 0,000200917 | 0,092283586 | 14273 | SELP |

| | | | | | | |
|---|---|---|---|---|---|---|
| chr1_192158370_192161157 | pre-BCG1_v_BCG9_down | -1,087905639 | 0,000293885 | 0,11900522 | 32177 | RGS18 |
| chr1_198638405_198644441 | pre-BCG1_v_BCG9_down | -0,977196085 | 0,000626305 | 0,179021931 | 33253 | PTPRC |
| chr1_241531388_241532998 | pre-BCG1_v_BCG9_down | -0,658361993 | 8,76E-05 | 0,061244017 | -11663 | RGS7 |
| chr1_245953572_245954431 | pre-BCG1_v_BCG9_down | -0,944227794 | 6,91E-05 | 0,053743783 | 635715 | KIF26B |
| chr1_44730816_44731405 | pre-BCG1_v_BCG9_down | -0,667614992 | 0,00781591 | 0,294608417 | 51948 | DMAP1 |
| chr1_78003996_78005999 | pre-BCG1_v_BCG9_down | -0,622532517 | 0,000459754 | 0,154073039 | 143345 | ZZZ3 |
| chr1_84144173_84146537 | pre-BCG1_v_BCG9_down | -1,18962894 | 5,22E-05 | 0,049776798 | 319460 | TTLL7 |
| chr1_89069767_89070467 | pre-BCG1_v_BCG9_down | -0,667801267 | 0,007708476 | 0,29374227 | -79788 | PKN2 |
| chr10_133278153_133278741 | pre-BCG1_v_BCG9_down | -0,631809158 | 0,017391345 | 0,358634266 | -469508 | PPP2R2D |
| chr10_3828579_3829322 | pre-BCG1_v_BCG9_down | -0,657185768 | 0,0127841 | 0,334936875 | -1484 | KLF6 |
| chr10_63264732_63269666 | pre-BCG1_v_BCG9_down | -0,722467206 | 0,000594066 | 0,174952403 | -53991 | TMEM26 |
| chr11_66100522_66101122 | pre-BCG1_v_BCG9_down | -0,581296235 | 0,049934861 | 0,465008002 | 3178 | RIN1 |
| chr12_10386640_10390138 | pre-BCG1_v_BCG9_down | -1,284012033 | 4,15E-05 | 0,044708307 | 22985 | GABARAPL1 |
| chr12_10407631_10409685 | pre-BCG1_v_BCG9_down | -0,646779721 | 0,004169511 | 0,264296386 | 43254 | GABARAPL1 |
| chr12_118358663_118359778 | pre-BCG1_v_BCG9_down | -0,992610273 | 0,00017394 | 0,087424315 | 47567 | KSR2 |
| chr12_123075234_123076132 | pre-BCG1_v_BCG9_down | -0,737987159 | 0,005688876 | 0,270438199 | 63888 | KNTC1 |
| chr12_14385600_14386559 | pre-BCG1_v_BCG9_down | -1,441325756 | 4,37E-05 | 0,044708307 | -132536 | ATF7IP |
| chr12_57768905_57770049 | pre-BCG1_v_BCG9_down | -2,205469694 | 0,005650127 | 0,270438199 | 55311 | R3HDM2 |
| chr12_57771011_57771402 | pre-BCG1_v_BCG9_down | -1,845770782 | 0,019279017 | 0,371579305 | 53581 | R3HDM2 |
| chr12_68158844_68159804 | pre-BCG1_v_BCG9_down | -0,751003093 | 0,027173491 | 0,394186417 | 116823 | DYRK2 |
| chr12_8513426_8514868 | pre-BCG1_v_BCG9_down | -0,934675453 | 0,000900626 | 0,183834761 | -94375 | CLEC6A |
| chr12_8989434_8991604 | pre-BCG1_v_BCG9_down | -0,920820825 | 0,000403518 | 0,140056889 | 15451 | A2ML1 |
| chr12_9827324_9830544 | pre-BCG1_v_BCG9_down | -0,700076469 | 0,011387473 | 0,326392462 | 6625 | CLEC2D |
| chr12_9864997_9870634 | pre-BCG1_v_BCG9_down | -0,819628686 | 4,00E-05 | 0,044708307 | 18079 | CLECL1 |
| chr13_40978878_40983360 | pre-BCG1_v_BCG9_down | -0,633897378 | 0,002339089 | 0,237208728 | 259615 | FOXO1 |
| chr13_41016040_41019586 | pre-BCG1_v_BCG9_down | -0,715707753 | 0,000116101 | 0,073766482 | 222921 | FOXO1 |
| chr13_49109963_49111218 | pre-BCG1_v_BCG9_down | -0,684627304 | 0,010621443 | 0,32138843 | -3222 | RCBTB2 |
| chr13_99256598_99259125 | pre-BCG1_v_BCG9_down | -0,899393353 | 5,78E-06 | 0,02808669 | -83610 | STK24 |
| chr14_60184086_60184518 | pre-BCG1_v_BCG9_down | -0,609980056 | 0,013189727 | 0,335128206 | 153382 | RTN1 |
| chr14_88004823_88007605 | pre-BCG1_v_BCG9_down | -0,662328417 | 3,75E-06 | 0,024327429 | 453401 | GALC |
| chr17_31317766_31321936 | pre-BCG1_v_BCG9_down | -0,731052707 | 0,000686461 | 0,180466765 | 964 | SPACA3 |
| chr17_46361441_46361887 | pre-BCG1_v_BCG9_down | -0,746483283 | 0,00490629 | 0,26789583 | 145888 | SKAP1 |

| | | | | | | |
|---|---|---|---|---|---|---|
| chr18_2980276_2982873 | pre-BCG1_v_BCG9_down | -0,585071976 | 0,003941198 | 0,264101006 | 30370 | LPIN2 |
| chr2_119929630_119931996 | pre-BCG1_v_BCG9_down | -1,232944842 | 3,33E-05 | 0,044708307 | -14348 | C1QL2 |
| chr2_142877514_142879767 | pre-BCG1_v_BCG9_down | -0,81069971 | 0,000869498 | 0,183834761 | 10629 | LRP1B |
| chr2_143128594_143130833 | pre-BCG1_v_BCG9_down | -1,182886117 | 4,34E-05 | 0,044708307 | -240444 | LRP1B |
| chr2_157435848_157436462 | pre-BCG1_v_BCG9_down | -0,707536247 | 0,013776171 | 0,335128206 | 144202 | GPD2 |
| chr2_159515626_159516942 | pre-BCG1_v_BCG9_down | -0,735762245 | 5,38E-05 | 0,049776798 | -135545 | DAPL1 |
| chr2_191148769_191149446 | pre-BCG1_v_BCG9_down | -1,027011355 | 0,001639788 | 0,204647596 | 35663 | HIBCH |
| chr2_213149684_213150692 | pre-BCG1_v_BCG9_down | -0,583018632 | 0,024454166 | 0,384016298 | 253377 | ERBB4 |
| chr2_38600354_38601147 | pre-BCG1_v_BCG9_down | -0,620559337 | 0,020050312 | 0,3743268 | 3653 | ATL2 |
| chr2_71417000_71417726 | pre-BCG1_v_BCG9_down | -0,626200131 | 0,003366608 | 0,255079978 | 36850 | PAIP2B |
| chr2_98660684_98663507 | pre-BCG1_v_BCG9_down | -0,667333981 | 0,033440961 | 0,421346713 | -41559 | VWA3B |
| chr21_14543894_14546545 | pre-BCG1_v_BCG9_down | -1,135198724 | 0,000617716 | 0,179021931 | -437278 | POTED |
| chr21_24428544_24429525 | pre-BCG1_v_BCG9_down | -0,598169548 | 0,002268601 | 0,235801044 | NONE | NONE |
| chr3_108822618_108825023 | pre-BCG1_v_BCG9_down | -0,623638668 | 0,040083369 | 0,440667668 | 13168 | MORC1 |
| chr3_151201296_151203233 | pre-BCG1_v_BCG9_down | -0,610098525 | 0,015227304 | 0,346550342 | -25768 | IGSF10 |
| chr3_152243322_152246923 | pre-BCG1_v_BCG9_down | -0,944695241 | 4,27E-05 | 0,044708307 | -186344 | TMEM14E |
| chr3_16888067_16889077 | pre-BCG1_v_BCG9_down | -0,674705656 | 0,01294649 | 0,335128206 | -37880 | PLCL2 |
| chr3_28292893_28294004 | pre-BCG1_v_BCG9_down | -0,767515595 | 0,002352496 | 0,237208728 | 10363 | CMC1 |
| chr4_147057356_147057873 | pre-BCG1_v_BCG9_down | -0,706427831 | 0,006373483 | 0,278385166 | -39266 | LSM6 |
| chr4_39030525_39032408 | pre-BCG1_v_BCG9_down | -0,94485086 | 4,23E-05 | 0,044708307 | 2574 | TMEM156 |
| chr5_58491289_58493150 | pre-BCG1_v_BCG9_down | -0,94110653 | 0,000720142 | 0,181618481 | 613353 | RAB3C |
| chr5_95080139_95081875 | pre-BCG1_v_BCG9_down | -0,645728733 | 9,14E-05 | 0,061244017 | 14128 | RHOBTB3 |
| chr5_96896313_96896924 | pre-BCG1_v_BCG9_down | -0,825441519 | 0,001528131 | 0,20205633 | -377655 | RIOK2 |
| chr6_127899285_127901264 | pre-BCG1_v_BCG9_down | -0,775104355 | 0,001325887 | 0,20205633 | 1956 | C6orf58 |
| chr6_156781504_156782711 | pre-BCG1_v_BCG9_down | -1,278193158 | 1,27E-05 | 0,037885069 | -316955 | ARID1B |
| chr6_158649979_158651155 | pre-BCG1_v_BCG9_down | -0,649509238 | 0,005124786 | 0,26789583 | 61183 | GTF2H5 |
| chr7_116027824_116031041 | pre-BCG1_v_BCG9_down | -0,657606305 | 0,007801748 | 0,294608417 | -110011 | CAV2 |
| chr7_148962607_148964251 | pre-BCG1_v_BCG9_down | -0,636625086 | 3,58E-05 | 0,044708307 | 4167 | ZNF783 |
| chr7_33039359_33040256 | pre-BCG1_v_BCG9_down | -0,791493562 | 0,00285903 | 0,244645504 | 42791 | FKBP9 |
| chr7_38250186_38251605 | pre-BCG1_v_BCG9_down | -1,204282607 | 0,003985543 | 0,264101006 | 33072 | STARD3NL |
| chr7_38251676_38253400 | pre-BCG1_v_BCG9_down | -0,79144902 | 0,024428606 | 0,384016298 | 34714 | STARD3NL |
| chr7_38332375_38336911 | pre-BCG1_v_BCG9_down | -0,626278968 | 0,001668702 | 0,206589598 | 116819 | STARD3NL |

| | | | | | | | |
|---|---|---|---|---|---|---|---|
| chr7_80638066_80642175 | pre-BCG1_v_BCG9_down | -1,012602461 | 0,000907966 | 0,183834761 | -91454 | SEMA3C | |
| chr9_121121259_121122937 | pre-BCG1_v_BCG9_down | -0,656774721 | 0,001496268 | 0,20205633 | 655448 | TLR4 | |
| chr9_122900911_122901710 | pre-BCG1_v_BCG9_down | -1,323583055 | 1,93E-05 | 0,037885069 | 441125 | CDK5RAP2 | |
| chr9_36143729_36145208 | pre-BCG1_v_BCG9_down | -0,67990964 | 0,000300583 | 0,119233323 | 7727 | GLIPR2 | |
| chrX_79144638_79147652 | pre-BCG1_v_BCG9_down | -0,818285055 | 0,002161955 | 0,230639024 | -131596 | TBX22 | |
| chr1_159945492_159946025 | pre-BCG1_v_pre-BCG10_up | 0,599766636 | 0,03209691 | 0,62614993 | -21715 | SLAMF9 | |
| chr1_239386557_239387345 | pre-BCG1_v_pre-BCG10_up | 0,714345994 | 0,000936275 | 0,544998581 | -405422 | CHRM3 | |
| chr1_247517760_247518647 | pre-BCG1_v_pre-BCG10_up | 0,693538699 | 0,002543117 | 0,548740748 | -23159 | ZNF496 | |
| chr1_46203088_46203819 | pre-BCG1_v_pre-BCG10_up | 0,613732358 | 0,000470679 | 0,508254413 | 12867 | IPP | |
| chr1_6440247_6440965 | pre-BCG1_v_pre-BCG10_up | 0,734187564 | 0,011547996 | 0,58050417 | 5267 | ACOT7 | |
| chr1_66751932_66752852 | pre-BCG1_v_pre-BCG10_up | 0,606981058 | 0,002828448 | 0,548740748 | -247573 | SGIP1 | |
| chr1_85576536_85577452 | pre-BCG1_v_pre-BCG10_up | 0,655845049 | 0,002426532 | 0,544998581 | 48990 | WDR63 | |
| chr10_13891394_13892137 | pre-BCG1_v_pre-BCG10_up | 0,676195399 | 0,030766331 | 0,623045375 | 262839 | PRPF18 | |
| chr10_18140266_18140963 | pre-BCG1_v_pre-BCG10_up | 0,608992923 | 0,043435039 | 0,644410661 | 42263 | MRC1 | |
| chr10_26391595_26392685 | pre-BCG1_v_pre-BCG10_up | 0,589421725 | 0,008759807 | 0,548740748 | -113096 | GAD2 | |
| chr10_48523908_48524321 | pre-BCG1_v_pre-BCG10_up | 0,913121803 | 0,008657688 | 0,548740748 | -85139 | GDF10 | |
| chr10_51074164_51075050 | pre-BCG1_v_pre-BCG10_up | 0,778980423 | 0,022794524 | 0,599856373 | 56108 | PARG | |
| chr10_77636776_77638793 | pre-BCG1_v_pre-BCG10_up | 0,762577172 | 0,004933595 | 0,548740748 | 95266 | C10orf11 | |
| chr10_93600817_93601603 | pre-BCG1_v_pre-BCG10_up | 0,773266827 | 0,010191534 | 0,571815561 | 43141 | TNKS2 | |
| chr10_99536791_99537421 | pre-BCG1_v_pre-BCG10_up | 0,761423888 | 0,026562141 | 0,604315032 | -5397 | SFRP5 | |
| chr11_111512299_111512995 | pre-BCG1_v_pre-BCG10_up | 0,614746898 | 0,008608361 | 0,548740748 | 39532 | SIK2 | |
| chr11_117186147_117187417 | pre-BCG1_v_pre-BCG10_up | 0,914102579 | 0,003393023 | 0,548740748 | 190 | BACE1 | |
| chr11_118621218_118621750 | pre-BCG1_v_pre-BCG10_up | 0,645311212 | 0,025950607 | 0,60063625 | 40361 | DDX6 | |
| chr11_129375636_129376399 | pre-BCG1_v_pre-BCG10_up | 1,063460078 | 0,00940657 | 0,552373103 | 130183 | BARX2 | |
| chr11_46277608_46278138 | pre-BCG1_v_pre-BCG10_up | 1,032780291 | 0,00715079 | 0,548740748 | -21339 | CREB3L1 | |
| chr11_72589953_72590556 | pre-BCG1_v_pre-BCG10_up | 0,686115477 | 0,0366869 | 0,627714142 | 64902 | ATG16L2 | Yes |
| chr11_9091129_9091919 | pre-BCG1_v_pre-BCG10_up | 0,682012912 | 0,005419495 | 0,548740748 | 21622 | SCUBE2 | |
| chr11_94740503_94741257 | pre-BCG1_v_pre-BCG10_up | 0,733804523 | 0,000681149 | 0,544998581 | -17542 | KDM4E | Yes |
| chr12_125186183_125186787 | pre-BCG1_v_pre-BCG10_up | 0,654285814 | 0,026638806 | 0,604315032 | 161908 | SCARB1 | |
| chr12_14216420_14217014 | pre-BCG1_v_pre-BCG10_up | 0,587072436 | 0,048509191 | 0,649412094 | -83664 | GRIN2B | |
| chr12_33439120_33439771 | pre-BCG1_v_pre-BCG10_up | 0,826087165 | 0,007414729 | 0,548740748 | 153308 | SYT10 | |
| chr12_55681472_55682478 | pre-BCG1_v_pre-BCG10_up | 0,830076471 | 0,020919145 | 0,595272408 | 7041 | OR6C6 | |

| | | | | | | | |
|---|---|---|---|---|---|---|---|
| chr12_58920109_58920638 | pre-BCG1_v_pre-BCG10_up | 0,980021901 | 0,007102775 | 0,548740748 | 393929 | LRIG3 | |
| chr12_77324798_77325271 | pre-BCG1_v_pre-BCG10_up | 0,788325855 | 0,017852725 | 0,594721648 | -52195 | CSRP2 | |
| chr12_88579703_88580716 | pre-BCG1_v_pre-BCG10_up | 0,682296286 | 0,001843994 | 0,544998581 | 44137 | TMTC3 | |
| chr13_36919663_36920530 | pre-BCG1_v_pre-BCG10_up | 0,640752542 | 0,004455649 | 0,548740748 | 837 | SPG20 | |
| chr14_88322950_88323848 | pre-BCG1_v_pre-BCG10_up | 0,590380833 | 0,039049434 | 0,635425224 | 136216 | GALC | |
| chr15_29821480_29822596 | pre-BCG1_v_pre-BCG10_up | 0,61923345 | 0,004616746 | 0,548740748 | 40889 | FAM189A1 | |
| chr15_51093837_51095099 | pre-BCG1_v_pre-BCG10_up | 0,776856216 | 0,00045586 | 0,508254413 | -36463 | SPPL2A | |
| chr15_68781658_68782061 | pre-BCG1_v_pre-BCG10_up | 0,601982932 | 0,04573435 | 0,644410661 | -57368 | ITGA11 | |
| chr15_93072319_93073342 | pre-BCG1_v_pre-BCG10_up | 0,580935315 | 0,040905809 | 0,640955277 | 126357 | FAM174B | |
| chr16_28505586_28508309 | pre-BCG1_v_pre-BCG10_up | 0,692492995 | 0,015723742 | 0,594721648 | -93 | CLN3 | |
| chr16_3112406_3113051 | pre-BCG1_v_pre-BCG10_up | 0,62487336 | 0,01372673 | 0,593059077 | -2656 | IL32 | Yes |
| chr16_3364992_3365653 | pre-BCG1_v_pre-BCG10_up | 0,629562943 | 0,019893063 | 0,594721648 | -9678 | TIGD7 | |
| chr16_57998320_57999172 | pre-BCG1_v_pre-BCG10_up | 0,821878644 | 0,001850081 | 0,544998581 | 6270 | CNGB1 | |
| chr16_66517060_66517572 | pre-BCG1_v_pre-BCG10_up | 0,727188274 | 0,017830062 | 0,594721648 | 56116 | BEAN1 | |
| chr16_66578946_66579630 | pre-BCG1_v_pre-BCG10_up | 0,702035549 | 0,012911627 | 0,586843988 | 5027 | TK2 | |
| chr16_83968176_83968703 | pre-BCG1_v_pre-BCG10_up | 0,59445605 | 0,03642109 | 0,626474973 | -14232 | OSGIN1 | |
| chr16_85169302_85169957 | pre-BCG1_v_pre-BCG10_up | 0,638125394 | 0,023936899 | 0,60063625 | -23516 | FAM92B | |
| chr16_87945228_87945849 | pre-BCG1_v_pre-BCG10_up | 0,623762473 | 0,012506126 | 0,584685955 | 24583 | CA5A | |
| chr17_19579663_19580481 | pre-BCG1_v_pre-BCG10_up | 0,674243969 | 0,022639442 | 0,599856373 | 28008 | ALDH3A2 | |
| chr17_44911270_44911728 | pre-BCG1_v_pre-BCG10_up | 0,967468488 | 0,00148077 | 0,544998581 | -15373 | WNT3 | |
| chr17_47423381_47423996 | pre-BCG1_v_pre-BCG10_up | 0,638155876 | 0,01747399 | 0,594721648 | 16146 | ZNF652 | |
| chr17_68599032_68601980 | pre-BCG1_v_pre-BCG10_up | 0,987211884 | 0,001664434 | 0,544998581 | 434830 | KCNJ2 | |
| chr17_74861124_74861929 | pre-BCG1_v_pre-BCG10_up | 0,743302875 | 0,022396088 | 0,599856373 | -7202 | MGAT5B | |
| chr17_78258513_78259494 | pre-BCG1_v_pre-BCG10_up | 1,152954686 | 0,000148589 | 0,433052116 | 24337 | RNF213 | |
| chr18_68721796_68722330 | pre-BCG1_v_pre-BCG10_up | 0,615846797 | 0,024985644 | 0,60063625 | -404083 | GTSCR1 | |
| chr19_1000174_1000965 | pre-BCG1_v_pre-BCG10_up | 0,617785294 | 0,006606671 | 0,548740748 | 152 | GRIN3B | |
| chr19_1154578_1155306 | pre-BCG1_v_pre-BCG10_up | 0,814920222 | 0,015665864 | 0,594721648 | 19340 | SBNO2 | |
| chr19_18021848_18022310 | pre-BCG1_v_pre-BCG10_up | 0,624360238 | 0,046497421 | 0,645206643 | -23826 | CCDC124 | |
| chr19_2610845_2611516 | pre-BCG1_v_pre-BCG10_up | 0,613873657 | 8,27E-05 | 0,401982801 | 91526 | GNG7 | |
| chr19_2720891_2721772 | pre-BCG1_v_pre-BCG10_up | 0,602782008 | 0,004274399 | 0,548740748 | 84 | DIRAS1 | |
| chr19_35673137_35673721 | pre-BCG1_v_pre-BCG10_up | 0,683554058 | 0,010375829 | 0,573432612 | 27753 | FXYD5 | |
| chr19_36421332_36422109 | pre-BCG1_v_pre-BCG10_up | 0,708318616 | 0,047638346 | 0,646048229 | -4731 | LRFN3 | |

| | | | | | | |
|---|---|---|---|---|---|---|
| chr19_44808740_44809365 | pre-BCG1_v_pre-BCG10_up | 0,598885496 | 0,042168133 | 0,642840791 | 125 | ZNF235 |
| chr19_51018636_51019291 | pre-BCG1_v_pre-BCG10_up | 0,745609755 | 0,008072415 | 0,548740748 | -1822 | ASPDH |
| chr19_53982086_53983094 | pre-BCG1_v_pre-BCG10_up | 0,581038526 | 0,002093945 | 0,544998581 | 11601 | ZNF813 |
| chr19_7645454_7647007 | pre-BCG1_v_pre-BCG10_up | 0,70531429 | 0,002398524 | 0,544998581 | -14557 | CAMSAP3 |
| chr19_7917319_7918073 | pre-BCG1_v_pre-BCG10_up | 0,629756327 | 0,018023732 | 0,594721648 | 6267 | EVI5L |
| chr2_130963650_130964280 | pre-BCG1_v_pre-BCG10_up | 0,859778076 | 0,018443233 | 0,594721648 | -7931 | TUBA3E |
| chr2_176122575_176123082 | pre-BCG1_v_pre-BCG10_up | 0,747030181 | 0,002124942 | 0,544998581 | -73494 | ATP5G3 |
| chr2_199463129_199463701 | pre-BCG1_v_pre-BCG10_up | 0,611853714 | 0,012444525 | 0,584261418 | 793989 | PLCL1 |
| chr2_207681283_207681814 | pre-BCG1_v_pre-BCG10_up | 1,088225631 | 0,006846216 | 0,548740748 | -51278 | MDH1B |
| chr2_218065225_218065862 | pre-BCG1_v_pre-BCG10_up | 0,726303877 | 0,037271135 | 0,630464383 | -340757 | TNP1 |
| chr2_222671267_222672315 | pre-BCG1_v_pre-BCG10_up | 0,589751762 | 0,001404558 | 0,544998581 | -234781 | EPHA4 |
| chr2_38535872_38536457 | pre-BCG1_v_pre-BCG10_up | 0,629447023 | 0,006482044 | 0,548740748 | 68239 | ATL2 |
| chr2_47369090_47369828 | pre-BCG1_v_pre-BCG10_up | 0,75061791 | 0,008096932 | 0,548740748 | 13058 | C2orf61 |
| chr2_56183678_56184972 | pre-BCG1_v_pre-BCG10_up | 0,637187569 | 0,014171787 | 0,594721648 | -33051 | EFEMP1 |
| chr2_73981102_73981611 | pre-BCG1_v_pre-BCG10_up | 0,944428154 | 0,002599148 | 0,548740748 | -16844 | TPRKB |
| chr20_1184994_1185901 | pre-BCG1_v_pre-BCG10_up | 0,607463888 | 0,014259035 | 0,594721648 | -20331 | TMEM74B |
| chr20_32031166_32031739 | pre-BCG1_v_pre-BCG10_up | 0,655006331 | 0,031904816 | 0,62614993 | 245 | SNTA1 |
| chr20_38164942_38165424 | pre-BCG1_v_pre-BCG10_up | 0,797586989 | 0,015602204 | 0,594721648 | 574205 | DHX35 |
| chr22_19723411_19724251 | pre-BCG1_v_pre-BCG10_up | 0,721987369 | 0,040840498 | 0,640955277 | 13363 | GP1BB |
| chr22_38958866_38960002 | pre-BCG1_v_pre-BCG10_up | 0,628424025 | 0,046106616 | 0,645193875 | 6857 | DMC1 |
| chr3_12004155_12004931 | pre-BCG1_v_pre-BCG10_up | 0,630614838 | 0,021619581 | 0,597025879 | -116150 | TAMM41 |
| chr3_127915089_127915604 | pre-BCG1_v_pre-BCG10_up | 0,648151194 | 0,044203621 | 0,644410661 | 43050 | EEFSEC |
| chr3_138434348_138435340 | pre-BCG1_v_pre-BCG10_up | 0,808285111 | 0,000640263 | 0,544998581 | 107393 | FAIM |
| chr3_141051071_141051620 | pre-BCG1_v_pre-BCG10_up | 0,970285492 | 0,003790667 | 0,548740748 | -55308 | ZBTB38 |
| chr3_46565820_46566396 | pre-BCG1_v_pre-BCG10_up | 0,711577826 | 0,017778708 | 0,594721648 | 27127 | RTP3 |
| chr3_46926648_46927199 | pre-BCG1_v_pre-BCG10_up | 0,963219541 | 0,015383776 | 0,594721648 | 2077 | PTH1R |
| chr3_9916678_9917495 | pre-BCG1_v_pre-BCG10_up | 1,169343007 | 9,51E-07 | 0,018489064 | 3108 | CIDEC |
| chr4_165378790_165379636 | pre-BCG1_v_pre-BCG10_up | 0,709273799 | 0,001315938 | 0,544998581 | -74011 | MARCHF1 |
| chr5_120465493_120465806 | pre-BCG1_v_pre-BCG10_up | 0,787283331 | 0,019980349 | 0,594721648 | 665631 | PRR16 |
| chr5_149271493_149272437 | pre-BCG1_v_pre-BCG10_up | 0,589386543 | 0,016270069 | 0,594721648 | 52391 | PDE6A |
| chr5_31854616_31855318 | pre-BCG1_v_pre-BCG10_up | 0,655898918 | 0,024276151 | 0,60063625 | 215450 | PDZD2 |
| chr5_88680214_88680682 | pre-BCG1_v_pre-BCG10_up | 0,633687915 | 0,007283208 | 0,548740748 | -480549 | MEF2C |

| | | | | | | |
|---|---|---|---|---|---|---|
| chr6_10523873_10524547 | pre-BCG1_v_pre-BCG10_up | 0,705498004 | 0,001970618 | 0,544998581 | -4379 | GCNT2 |
| chr6_147508112_147508790 | pre-BCG1_v_pre-BCG10_up | 0,749886735 | 0,022300569 | 0,599856373 | -17218 | STXBP5 |
| chr6_159168949_159170125 | pre-BCG1_v_pre-BCG10_up | 0,63725387 | 0,004409672 | 0,548740748 | 70907 | EZR |
| chr7_130491941_130492469 | pre-BCG1_v_pre-BCG10_up | 0,767070171 | 0,015581388 | 0,594721648 | -73317 | KLF14 |
| chr7_16850822_16851660 | pre-BCG1_v_pre-BCG10_up | 0,616428428 | 0,008906597 | 0,548740748 | -6537 | AGR2 |
| chr7_28957866_28958469 | pre-BCG1_v_pre-BCG10_up | 0,795621885 | 0,034791568 | 0,626474973 | -275860 | CHN2 |
| chr7_35971340_35971856 | pre-BCG1_v_pre-BCG10_up | 0,854108682 | 0,012544774 | 0,584685955 | 130971 | SEPT7 |
| chr7_39609558_39610097 | pre-BCG1_v_pre-BCG10_up | 0,692997094 | 0,039287252 | 0,636254061 | 3853 | YAE1D1 |
| chr7_39833152_39833706 | pre-BCG1_v_pre-BCG10_up | 0,595086104 | 0,016168829 | 0,594721648 | -156207 | CDK13 |
| chr7_64463539_64464431 | pre-BCG1_v_pre-BCG10_up | 0,710418485 | 0,012356707 | 0,583420865 | 3046 | ERV3-1 |
| chr7_6526535_6527062 | pre-BCG1_v_pre-BCG10_up | 0,795607526 | 0,001981361 | 0,544998581 | -2926 | KDELR2 |
| chr7_73624006_73624641 | pre-BCG1_v_pre-BCG10_up | 0,783004917 | 0,008285905 | 0,548740748 | 576 | LAT2 |
| chr7_76281323_76281940 | pre-BCG1_v_pre-BCG10_up | 0,666800073 | 0,022398838 | 0,599856373 | -25074 | POMZP3 |
| chr8_103499318_103500979 | pre-BCG1_v_pre-BCG10_up | 0,594539971 | 0,000443046 | 0,508254413 | -63651 | ODF1 |
| chr8_121639626_121640067 | pre-BCG1_v_pre-BCG10_up | 0,722472369 | 0,036094072 | 0,626474973 | 182181 | MTBP |
| chr8_144332010_144332542 | pre-BCG1_v_pre-BCG10_up | 0,777658363 | 0,020901145 | 0,595272408 | 3178 | ZFP41 |
| chr8_21787326_21788103 | pre-BCG1_v_pre-BCG10_up | 0,847886751 | 0,002772556 | 0,548740748 | 10533 | XPO7 |
| chr8_38974112_38974864 | pre-BCG1_v_pre-BCG10_up | 0,640222771 | 0,013688614 | 0,593059077 | 9320 | ADAM32 |
| chr8_76677782_76678320 | pre-BCG1_v_pre-BCG10_up | 0,694797558 | 0,010430628 | 0,574334587 | 225947 | HNF4G |
| chr8_79611867_79612383 | pre-BCG1_v_pre-BCG10_up | 0,742441311 | 0,013633655 | 0,593059077 | 33843 | ZC2HC1A |
| chr9_121736375_121736868 | pre-BCG1_v_pre-BCG10_up | 0,775484482 | 0,011635617 | 0,58050417 | 395123 | BRINP1 |
| chr9_122213608_122214358 | pre-BCG1_v_pre-BCG10_up | 0,656148186 | 0,000822974 | 0,544998581 | -82238 | BRINP1 |
| chr9_130938798_130939748 | pre-BCG1_v_pre-BCG10_up | 0,75822217 | 0,008264262 | 0,548740748 | 16734 | C9orf16 |
| chr9_38423910_38424641 | pre-BCG1_v_pre-BCG10_up | 0,852779073 | 0,015019841 | 0,594721648 | 168 | IGFBPL1 |
| chr9_41235084_41235631 | pre-BCG1_v_pre-BCG10_up | 0,635721704 | 0,035660341 | 0,626474973 | 92006 | SPATA31A4 |
| chr9_4490073_4491116 | pre-BCG1_v_pre-BCG10_up | 0,730455972 | 0,023552036 | 0,60063625 | 151 | SLC1A1 |
| chr9_93131329_93131827 | pre-BCG1_v_pre-BCG10_up | 0,796112484 | 0,017623562 | 0,594721648 | 273808 | DIRAS2 |
| chrX_11427265_11428190 | pre-BCG1_v_pre-BCG10_up | 0,648700614 | 0,005955701 | 0,548740748 | 116195 | AMELX |
| chrX_134990818_134991429 | pre-BCG1_v_pre-BCG10_up | 0,595369647 | 0,031661952 | 0,625963685 | 65098 | MMGT1 |
| chrX_65407078_65407688 | pre-BCG1_v_pre-BCG10_up | 0,735682222 | 0,022700228 | 0,599856373 | 24992 | HEPH |
| chr1_117001991_117002583 | pre-BCG1_v_pre-BCG10_down | -0,790505144 | 0,032139784 | 0,62614993 | 85798 | ATP1A1 |
| chr1_158930786_158934314 | pre-BCG1_v_pre-BCG10_down | -0,594576303 | 0,033768811 | 0,626474973 | 31208 | PYHIN1 |

| | | | | | | |
|---|---|---|---|---|---|---|
| chr1_198638405_198644441 | pre-BCG1_v_pre-BCG10_down | -0,634205202 | 0,034648769 | 0,626474973 | 33253 | PTPRC |
| chr1_40388717_40389694 | pre-BCG1_v_pre-BCG10_down | -0,659162379 | 0,020558266 | 0,594804894 | -21521 | MYCL |
| chr1_84144173_84146537 | pre-BCG1_v_pre-BCG10_down | -0,770703574 | 0,012366589 | 0,583420865 | 319460 | TTLL7 |
| chr10_6140817_6141751 | pre-BCG1_v_pre-BCG10_down | -0,596492092 | 0,023867073 | 0,60063625 | 9975 | RBM17 |
| chr10_96270807_96271460 | pre-BCG1_v_pre-BCG10_down | -0,759101416 | 0,00230029 | 0,544998581 | -34440 | HELLS |
| chr11_33197863_33198328 | pre-BCG1_v_pre-BCG10_down | -0,625573562 | 0,047875917 | 0,646675302 | -15085 | CSTF3 |
| chr12_10386640_10390138 | pre-BCG1_v_pre-BCG10_down | -0,650107969 | 0,047313327 | 0,645879721 | 22985 | GABARAPL1 |
| chr12_118358663_118359778 | pre-BCG1_v_pre-BCG10_down | -0,618503236 | 0,025083125 | 0,60063625 | 47567 | KSR2 |
| chr12_14385600_14386559 | pre-BCG1_v_pre-BCG10_down | -0,753641639 | 0,036185825 | 0,626474973 | -132536 | ATF7IP |
| chr12_15546470_15547122 | pre-BCG1_v_pre-BCG10_down | -0,599186683 | 0,032939295 | 0,62614993 | 71465 | PTPRO |
| chr12_31251921_31252469 | pre-BCG1_v_pre-BCG10_down | -0,620745497 | 0,024731492 | 0,60063625 | 25416 | DDX11 |
| chr12_8989434_8991604 | pre-BCG1_v_pre-BCG10_down | -0,791524434 | 0,003796001 | 0,548740748 | 15451 | A2ML1 |
| chr13_98570871_98571394 | pre-BCG1_v_pre-BCG10_down | -0,835667174 | 0,002874598 | 0,548740748 | -34796 | IPO5 |
| chr16_87490634_87491479 | pre-BCG1_v_pre-BCG10_down | -0,68202229 | 0,033927276 | 0,626474973 | 34594 | ZCCHC14 |
| chr17_18186002_18186641 | pre-BCG1_v_pre-BCG10_down | -0,656352758 | 0,046284813 | 0,645206643 | 21898 | MIEF2 |
| chr17_7851500_7851938 | pre-BCG1_v_pre-BCG10_down | -0,774987418 | 0,017748352 | 0,594721648 | 16246 | CNTROB |
| chr19_50418781_50420147 | pre-BCG1_v_pre-BCG10_down | -0,617572579 | 0,003075677 | 0,548740748 | -12936 | ATF5 |
| chr19_6996859_6998063 | pre-BCG1_v_pre-BCG10_down | -0,714203377 | 0,016982617 | 0,594721648 | -33128 | MBD3L5 |
| chr2_142877514_142879767 | pre-BCG1_v_pre-BCG10_down | -0,61281026 | 0,016816174 | 0,594721648 | 10629 | LRP1B |
| chr2_143128594_143130833 | pre-BCG1_v_pre-BCG10_down | -0,651792349 | 0,031347673 | 0,623648633 | -240444 | LRP1B |
| chr2_143248012_143248756 | pre-BCG1_v_pre-BCG10_down | -0,802845886 | 0,022592999 | 0,599856373 | -359114 | LRP1B |
| chr2_159515626_159516942 | pre-BCG1_v_pre-BCG10_down | -0,606399224 | 0,001577781 | 0,544998581 | -135545 | DAPL1 |
| chr2_203706607_203710225 | pre-BCG1_v_pre-BCG10_down | -0,588873166 | 0,033385114 | 0,626474973 | 27949 | ICA1L |
| chr21_14543894_14546545 | pre-BCG1_v_pre-BCG10_down | -0,738720884 | 0,033879481 | 0,626474973 | -437278 | POTED |
| chr21_17793651_17794173 | pre-BCG1_v_pre-BCG10_down | -0,692761368 | 0,021596394 | 0,597025879 | 691568 | USP25 |
| chr3_28292893_28294004 | pre-BCG1_v_pre-BCG10_down | -0,583330429 | 0,027024569 | 0,607256117 | 10363 | CMC1 |
| chr4_39030525_39032408 | pre-BCG1_v_pre-BCG10_down | -0,687976873 | 0,004429759 | 0,548740748 | 2574 | TMEM156 |
| chr5_150846511_150847193 | pre-BCG1_v_pre-BCG10_down | -0,615021289 | 0,015149072 | 0,594721648 | 19695 | SLC36A1 |
| chr5_58491289_58493150 | pre-BCG1_v_pre-BCG10_down | -0,741564163 | 0,011261264 | 0,5762077 | 613353 | RAB3C |
| chr5_96896313_96896924 | pre-BCG1_v_pre-BCG10_down | -0,722226679 | 0,008494528 | 0,548740748 | -377655 | RIOK2 |
| chr6_127899285_127901264 | pre-BCG1_v_pre-BCG10_down | -0,674495853 | 0,007816329 | 0,548740748 | 1956 | C6orf58 |
| chr6_156781504_156782711 | pre-BCG1_v_pre-BCG10_down | -0,823521896 | 0,007097316 | 0,548740748 | -316955 | ARID1B |

| | | | | | | |
|---|---|---|---|---|---|---|
| chr6_158649979_158651155 | pre-BCG1_v_pre-BCG10_down | -0,581363151 | 0,017123118 | 0,594721648 | 61183 | GTF2H5 |
| chr7_33039359_33040256 | pre-BCG1_v_pre-BCG10_down | -0,789710249 | 0,005050776 | 0,548740748 | 42791 | FKBP9 |
| chr7_38251676_38253400 | pre-BCG1_v_pre-BCG10_down | -0,894214286 | 0,015445648 | 0,594721648 | 34714 | STARD3NL |
| chr7_38302919_38304101 | pre-BCG1_v_pre-BCG10_down | -0,804916855 | 0,002467339 | 0,544998581 | 85686 | STARD3NL |
| chr8_23230613_23231216 | pre-BCG1_v_pre-BCG10_down | -0,724729047 | 0,018123245 | 0,594721648 | 30838 | LOXL2 |
| chr9_130038378_130038894 | pre-BCG1_v_pre-BCG10_down | -0,757030665 | 0,015427617 | 0,594721648 | 11831 | GARNL3 |
| chrX_131812614_131813431 | pre-BCG1_v_pre-BCG10_down | -0,604922928 | 0,001585819 | 0,544998581 | -239305 | MBNL3 |
| chrX_54921055_54921989 | pre-BCG1_v_pre-BCG10_down | -0,676063086 | 0,005176935 | 0,548740748 | -25722 | TRO |

**Supplementary Table 5 - List of selected candidate trained immunity genes.**

| Gene name | Mechanism/Pathway important for trained immunity (TI) | Reference for trained immunity |
|---|---|---|
| AIM2 | Inflammasome signaling | |
| ASC/PYCARD/CARD5 | Inflammasome signaling | Arts, R.J.W. et al., BCG Vaccination Protects against Experimental Viral Infection in Humans through the Induction of Cytokines Associated with Trained Immunity. Cell host & microbe 23, 2018. |
| ATG16L1 | Autophagy | |
| ATG16L2 | Autophagy | |
| ATG2B | Autophagy | Buffen, K. et al., Autophagy controls BCG-induced trained immunity and the response to intravesical BCG therapy for bladder cancer. PLoS pathogens 10, 2014. |
| ATG5 | Autophagy | |
| ATG7 | Autophagy | |
| CASP1 | Inflammasome signaling | |
| CASP5 | Inflammasome signaling | |
| EHMT2 | Epigenetic enzyme | Mourits, V.P. et al., Lysine methyltransferase G9a is an important modulator of trained immunity. Clinical & translational immunology 10, 2021. |
| GBP1 | Inflammasome signaling | |
| GBP2 | Inflammasome signaling | |
| GBP3 | Inflammasome signaling | |
| GBP4 | Inflammasome signaling | |
| GBP5 | Inflammasome signaling | |
| HK2 | Glycolysis | Arts, R.J.W. et al., Immunometabolic Pathways in BCG-Induced Trained Immunity. Cell reports 17, 2016. |
| HNF1A | Myelopoiesis | Cirovic, B. et al., BCG Vaccination in Humans Elicits Trained Immunity via the Hematopoietic Progenitor Compartment. Cell host & microbe 28, 2020. |
| HNF1B | Myelopoiesis | Cirovic, B. et al., BCG Vaccination in Humans Elicits Trained Immunity via the Hematopoietic Progenitor Compartment. Cell host & microbe 28, 2020. |
| IL18 | TI cytokine | |

| | | |
|---|---|---|
| IL1B | TI cytokine | Arts, R.J.W. et al., BCG Vaccination Protects against Experimental Viral Infection in Humans through the Induction of Cytokines Associated with Trained Immunity. Cell host & microbe 23, 2018. |
| IL1RN | TI cytokine | |
| IL32 | TI cytokine | Dos Santos, J.C. et al., β-Glucan-Induced Trained Immunity Protects against Leishmania braziliensis Infection: a Crucial Role for IL-32. Cell reports 28, 2019. |
| IL6 | TI cytokine | |
| KDM4A | Epigenetic enzyme | Moorlag, S. et al., An integrative genomics approach identifies KDM4 as a modulator of trained immunity. Eur J Immunol 2021. |
| KDM4B | Epigenetic enzyme | Moorlag, S. et al., An integrative genomics approach identifies KDM4 as a modulator of trained immunity. Eur J Immunol 2021. |
| KDM4C | Epigenetic enzyme | Moorlag, S. et al., An integrative genomics approach identifies KDM4 as a modulator of trained immunity. Eur J Immunol 2021. |
| KDM4D | Epigenetic enzyme | Moorlag, S. et al., An integrative genomics approach identifies KDM4 as a modulator of trained immunity. Eur J Immunol 2021. |
| KDM4E | Epigenetic enzyme | Moorlag, S. et al., An integrative genomics approach identifies KDM4 as a modulator of trained immunity. Eur J Immunol 2021. |
| LAMP2 | Autophagy | |
| NLRP3 | Inflammasome signaling | |
| NOD2/CARD15 | Autophagy | |
| PFKP | Glycolysis | Arts, R.J.W. et al., Immunometabolic Pathways in BCG-Induced Trained Immunity. Cell reports 17, 2016. |
| RIPK2 | Autophagy | |
| TNF | TI cytokine | |

**Supplementary Table 6 - Table with the differentially expressed genes between pre-BCG1 and BCG6.**
This table shows the differentially expressed genes at p(unadjusted)<0.05 between pre-BCG1 and BCG6.
Candidate trained immunity genes are indicated in the final column.

| Gene name | log2 Fold Change | p value | adjusted p value | Candidate trained immunity gene |
|---|---|---|---|---|
| FANCA | 0,416164968 | 2,79E-05 | 0,019944468 | |
| APOL2 | 0,392119005 | 5,05E-05 | 0,030298792 | |
| GBP1 | 0,837182252 | 0,000106425 | 0,052781665 | Yes |
| GBP2 | 0,535894975 | 0,000237511 | 0,114113007 | Yes |
| SLAMF8 | 0,824175426 | 0,000273093 | 0,129189933 | |
| ACE | 0,814849641 | 0,000395597 | 0,176293017 | |
| IRF9 | 0,403102363 | 0,000393838 | 0,176293017 | |
| PSTPIP2 | 0,425863886 | 0,000403355 | 0,177182411 | |
| AP000275.65 | 1,569202645 | 0,000428268 | 0,182900266 | |
| ALPK1 | 0,484670341 | 0,000554252 | 0,227017741 | |
| TGM2 | 0,709969574 | 0,000704876 | 0,277874771 | |
| IL27 | 1,035307566 | 0,000805327 | 0,305716094 | |
| KCNMA1 | 1,427699707 | 0,000833368 | 0,308737816 | |
| C8orf76 | 0,37165149 | 0,000827852 | 0,308737816 | |
| C1QA | 0,784136216 | 0,000909738 | 0,326236012 | |
| SDC3 | 0,679010653 | 0,000923039 | 0,326236012 | |
| APOL3 | 0,425167377 | 0,000956946 | 0,330619403 | |
| TAP1 | 0,38678848 | 0,001215249 | 0,406170553 | |
| SPDYE2 | -1,881858553 | 0,001232919 | 0,407645542 | |
| FCGBP | -0,858008712 | 0,001278171 | 0,418111355 | |
| JUN | -1,119329459 | 0,001344874 | 0,435300235 | |
| ODF3B | 0,662251968 | 0,001434222 | 0,459065053 | |
| GBP5 | 0,676898788 | 0,00160993 | 0,505140205 | Yes |
| MT2A | 0,510167379 | 0,001789389 | 0,544771425 | |
| SLC30A4 | -0,608250308 | 0,001857706 | 0,560025368 | |
| APOL4 | 1,002388629 | 0,001906898 | 0,565358661 | |
| BATF2 | 0,987772806 | 0,001937874 | 0,566131453 | |
| AL589988.1 | -3,24698225 | 0,001963859 | 0,566131453 | |
| C17orf81 | 0,465103748 | 0,002130619 | 0,601049521 | |
| SERPING1 | 0,882433958 | 0,002439069 | 0,678374816 | |
| C1QB | 1,049365496 | 0,00262936 | 0,709212181 | |
| MUC1 | 0,851786943 | 0,002740676 | 0,720282322 | |
| AIM2 | 0,713404476 | 0,002734595 | 0,720282322 | Yes |
| STAT1 | 0,523590248 | 0,0027902 | 0,727083529 | |
| DDB2 | 0,378676649 | 0,002844875 | 0,735101394 | |
| C2 | 0,858142626 | 0,002932122 | 0,745122465 | |
| IRS1 | -0,388456714 | 0,002909691 | 0,745122465 | |
| IGFBP3 | -0,524377716 | 0,003006993 | 0,751997231 | |
| CCL3 | -0,678866553 | 0,003008087 | 0,751997231 | |
| FGFBP2 | -0,541669259 | 0,003077319 | 0,763100596 | |

| | | | | |
|---|---|---|---|---|
| KLHDC7B | 0,545207163 | 0,003205436 | 0,764061534 | |
| GCH1 | 0,433329008 | 0,003261928 | 0,771546393 | |
| RUFY4 | 0,626651041 | 0,003478501 | 0,808708187 | |
| BOK | -0,857008397 | 0,003474398 | 0,808708187 | |
| CD40 | 0,47463582 | 0,003779509 | 0,842145826 | |
| C1QC | 1,173979553 | 0,003921315 | 0,867457017 | |
| ARRDC3 | -0,399533063 | 0,004330665 | 0,931971002 | |
| AKR1C3 | -0,596546359 | 0,004276409 | 0,931971002 | |
| HBA1 | -1,453582641 | 0,004334185 | 0,931971002 | |
| IL2RB | -0,519396504 | 0,004467671 | 0,954002865 | |
| PDGFD | -0,590699808 | 0,004618538 | 0,972708439 | |
| NCF1 | 0,39272474 | 0,00468607 | 0,980217368 | |
| RP5-977B1.10 | 1,677033087 | 0,049074668 | 1 | |
| MEF2BNB-MEF2B | 1,248862831 | 0,023548968 | 1 | |
| ANKRD22 | 1,09761395 | 0,008332145 | 1 | |
| TBC1D3C | 0,996891295 | 0,020485474 | 1 | |
| ETV7 | 0,947300542 | 0,00555 | 1 | |
| KREMEN1 | 0,883521368 | 0,0077558 | 1 | |
| CDKN3 | 0,816848738 | 0,009116359 | 1 | |
| CENPM | 0,715971748 | 0,039455664 | 1 | |
| EXOC3L1 | 0,707652082 | 0,022797635 | 1 | |
| FAM20A | 0,686287983 | 0,029008276 | 1 | |
| FCGR1A | 0,677957968 | 0,016560352 | 1 | |
| NME2 | 0,676923434 | 0,008742964 | 1 | |
| ITGA7 | 0,645611823 | 0,04749242 | 1 | |
| TNNT1 | 0,634606854 | 0,013908199 | 1 | |
| SEPTIN4 | 0,619245911 | 0,009684948 | 1 | |
| FCGR1B | 0,610630851 | 0,022187444 | 1 | |
| PRSS21 | 0,606527046 | 0,012742105 | 1 | |
| CD274 | 0,589720504 | 0,023818374 | 1 | |
| LGALS3BP | 0,58540446 | 0,018172071 | 1 | |
| GPER | 0,580884788 | 0,030727915 | 1 | |
| HMGB3 | 0,568119229 | 0,049107076 | 1 | |
| TNFSF10 | 0,56185307 | 0,010725483 | 1 | |
| CDCA5 | 0,552782326 | 0,02692685 | 1 | |
| CCDC154 | 0,545911996 | 0,049495278 | 1 | |
| PLA2G4B | 0,544689511 | 0,018914299 | 1 | |
| ZNF222 | 0,524210862 | 0,029833271 | 1 | |
| ZNF20 | 0,523429103 | 0,040410492 | 1 | |
| EPSTI1 | 0,515936225 | 0,013113973 | 1 | |
| LAP3 | 0,514206615 | 0,006674993 | 1 | |
| GBP4 | 0,50391882 | 0,006976171 | 1 | Yes |
| VAMP5 | 0,501162477 | 0,006757253 | 1 | |
| SLC6A12 | 0,500340913 | 0,033414764 | 1 | |
| C15orf52 | 0,4951153 | 0,035068023 | 1 | |

| | | | |
|---|---|---|---|
| MARCO | 0,491052503 | 0,017908252 | 1 |
| IL15RA | 0,487243844 | 0,008562457 | 1 |
| CDCA7 | 0,486726514 | 0,033914711 | 1 |
| SLC26A1 | 0,479122701 | 0,017501327 | 1 |
| CASP5 | 0,476573284 | 0,039961715 | 1 |
| MB21D1 | 0,465239875 | 0,02401357 | 1 |
| SAMD4A | 0,464845301 | 0,032420175 | 1 |
| SLC31A2 | 0,459762196 | 0,035448672 | 1 |
| WARS | 0,458540321 | 0,012393938 | 1 |
| CLEC12B | 0,454642725 | 0,014497235 | 1 |
| NUSAP1 | 0,453901785 | 0,038290692 | 1 |
| P2RY13 | 0,444467981 | 0,011625924 | 1 |
| C9orf139 | 0,442046919 | 0,019929476 | 1 |
| TYMP | 0,440500161 | 0,005591969 | 1 |
| MT-ATP6 | 0,432481895 | 0,039013272 | 1 |
| TICAM2 | 0,427650408 | 0,018495703 | 1 |
| UHRF1 | 0,427277685 | 0,017741661 | 1 |
| EPS8L1 | 0,426598084 | 0,030550779 | 1 |
| IRF1 | 0,424670196 | 0,014617206 | 1 |
| S100A13 | 0,420386919 | 0,049248263 | 1 |
| DDO | 0,416633561 | 0,035031103 | 1 |
| NAT1 | 0,413670028 | 0,032232423 | 1 |
| RPGRIP1 | 0,413229016 | 0,029692081 | 1 |
| PARP9 | 0,411534011 | 0,011503059 | 1 |
| FSTL3 | 0,409229824 | 0,042162937 | 1 |
| TNFAIP2 | 0,399135073 | 0,014081365 | 1 |
| HSPA6 | 0,398426692 | 0,011541368 | 1 |
| KCNMB1 | 0,393645458 | 0,012947058 | 1 |
| MT-ND6 | 0,390070066 | 0,046631752 | 1 |
| FANCL | 0,388642912 | 0,027142752 | 1 |
| ZNF684 | 0,387625487 | 0,01041037 | 1 |
| MT-ND2 | 0,386072627 | 0,044502769 | 1 |
| FAM26F | 0,384404995 | 0,037197222 | 1 |
| MT-CO1 | 0,381938213 | 0,029537893 | 1 |
| ZMYND15 | 0,38091803 | 0,045315862 | 1 |
| ECHDC3 | 0,377353262 | 0,046912018 | 1 |
| TRIM69 | 0,376236471 | 0,027204276 | 1 |
| J01415.25 | 0,375372852 | 0,035392278 | 1 |
| CES4A | 0,374279573 | 0,047361718 | 1 |
| LRRC27 | 0,372686606 | 0,012191912 | 1 |
| FAM176C | -0,372699564 | 0,021249875 | 1 |
| CD247 | -0,373601814 | 0,047762817 | 1 |
| ST3GAL3 | -0,377607622 | 0,037671208 | 1 |
| SYNE2 | -0,379180659 | 0,037297328 | 1 |
| YPEL1 | -0,381880614 | 0,025500207 | 1 |

| | | | |
|---|---|---|---|
| ACACB | -0,383224259 | 0,040739728 | 1 |
| MYBL1 | -0,38719472 | 0,04464206 | 1 |
| RP11-1000B6.4 | -0,388639614 | 0,0468449 | 1 |
| IKZF2 | -0,399388604 | 0,021510074 | 1 |
| CLIC3 | -0,400131458 | 0,035843828 | 1 |
| SBK1 | -0,406436272 | 0,045781816 | 1 |
| RP11-277P12.7 | -0,410040371 | 0,022980627 | 1 |
| TLE1 | -0,41215121 | 0,033816508 | 1 |
| ZFPM1 | -0,41284894 | 0,01168682 | 1 |
| GZMM | -0,413036446 | 0,049802174 | 1 |
| FCRL6 | -0,414475953 | 0,040292514 | 1 |
| CCDC144A | -0,419136586 | 0,047712206 | 1 |
| SLAIN1 | -0,419191877 | 0,025081596 | 1 |
| TKTL1 | -0,433273247 | 0,022781026 | 1 |
| RAB30 | -0,434651822 | 0,040036936 | 1 |
| CST7 | -0,435674472 | 0,01922203 | 1 |
| NCALD | -0,43658179 | 0,012586912 | 1 |
| ZNF853 | -0,439461289 | 0,037295084 | 1 |
| RALGPS2 | -0,439585199 | 0,014245521 | 1 |
| NSG1 | -0,4419114 | 0,029636123 | 1 |
| TSPAN5 | -0,443457068 | 0,025841639 | 1 |
| NKG7 | -0,443595085 | 0,019161666 | 1 |
| BICD1 | -0,446308637 | 0,006748931 | 1 |
| APBA2 | -0,446638838 | 0,043694812 | 1 |
| BCL7A | -0,447556619 | 0,027998413 | 1 |
| TGFBR3 | -0,45011796 | 0,024936332 | 1 |
| PRF1 | -0,450913705 | 0,024223918 | 1 |
| ODZ1 | -0,453200436 | 0,015824294 | 1 |
| DUSP14 | -0,455604709 | 0,034462795 | 1 |
| S1PR5 | -0,458137919 | 0,020863878 | 1 |
| HOPX | -0,458427845 | 0,020743564 | 1 |
| KIAA1377 | -0,458659193 | 0,03114167 | 1 |
| BANK1 | -0,460052878 | 0,032338396 | 1 |
| PTCH1 | -0,460556447 | 0,016763216 | 1 |
| ZNF831 | -0,460697733 | 0,02702505 | 1 |
| SH2D1B | -0,461225575 | 0,00812831 | 1 |
| CCL4 | -0,463561883 | 0,039497886 | 1 |
| SLFN12L | -0,472405125 | 0,031816351 | 1 |
| CCNJL | -0,474170767 | 0,008047792 | 1 |
| DLG3 | -0,476414349 | 0,014885564 | 1 |
| PKIG | -0,476434034 | 0,040815151 | 1 |
| F2R | -0,488892189 | 0,014278129 | 1 |
| TPD52 | -0,489713857 | 0,038071167 | 1 |
| TMEM182 | -0,490816156 | 0,04734152 | 1 |
| MYOM2 | -0,492099626 | 0,041615676 | 1 |

| | | | |
|---|---|---|---|
| LRFN3 | -0,494504291 | 0,043369254 | 1 |
| FAIM3 | -0,499637977 | 0,038238886 | 1 |
| PCDH1 | -0,502047296 | 0,037726825 | 1 |
| KLRD1 | -0,504481521 | 0,01425441 | 1 |
| SLC38A1 | -0,509394109 | 0,029474403 | 1 |
| AGAP1 | -0,509692461 | 0,028713559 | 1 |
| C1orf21 | -0,510947176 | 0,011027812 | 1 |
| DTHD1 | -0,514596186 | 0,004913806 | 1 |
| PRSS23 | -0,515501534 | 0,017573282 | 1 |
| MICAL3 | -0,516296685 | 0,015747501 | 1 |
| MAGEE1 | -0,523746835 | 0,033274446 | 1 |
| ZNF570 | -0,526085188 | 0,016430441 | 1 |
| NCR1 | -0,529873842 | 0,014836874 | 1 |
| BCL2 | -0,536146943 | 0,023566452 | 1 |
| TNFRSF13B | -0,541079191 | 0,025341118 | 1 |
| PAWR | -0,551909136 | 0,024349633 | 1 |
| ID3 | -0,567827047 | 0,034610038 | 1 |
| KLRB1 | -0,571230226 | 0,004903743 | 1 |
| CADM1 | -0,573608916 | 0,041904879 | 1 |
| ARL4D | -0,574269887 | 0,013595168 | 1 |
| OSBPL10 | -0,576863528 | 0,015454784 | 1 |
| LPL | -0,577805251 | 0,031316038 | 1 |
| SEZ6L | -0,593641217 | 0,031822139 | 1 |
| C14orf45 | -0,620995817 | 0,009404158 | 1 |
| FCRL2 | -0,64213295 | 0,02215549 | 1 |
| KIR3DL1 | -0,651587879 | 0,01219294 | 1 |
| NOS3 | -0,65337981 | 0,020387454 | 1 |
| KIR2DL1 | -0,653937262 | 0,019597256 | 1 |
| RNF165 | -0,67222023 | 0,004987637 | 1 |
| KLRC4 | -0,695047506 | 0,033990308 | 1 |
| ADAMTS1 | -0,704489831 | 0,029959714 | 1 |
| MANEAL | -0,708707868 | 0,005460479 | 1 |
| ZNF365 | -0,714988955 | 0,010056548 | 1 |
| MYEOV | -0,7367308 | 0,041151876 | 1 |
| HCAR2 | -0,737938864 | 0,032048177 | 1 |
| B3GNT7 | -0,755433666 | 0,012924886 | 1 |
| NT5E | -0,758367992 | 0,037755749 | 1 |
| FCRL1 | -0,806086183 | 0,013468328 | 1 |
| FCRLA | -0,816093861 | 0,008902949 | 1 |
| BEND5 | -0,835404712 | 0,014744109 | 1 |
| PAX5 | -0,835450924 | 0,031484747 | 1 |
| CD79A | -0,842368253 | 0,006208487 | 1 |
| AKAP5 | -0,848275099 | 0,011655695 | 1 |
| GRIP1 | -0,866412273 | 0,015946217 | 1 |
| FAM203B | -0,922599113 | 0,034347834 | 1 |

| | | | |
|---|---|---|---|
| CXCR5 | -0,928303049 | 0,007193372 | 1 |
| CA1 | -1,034054431 | 0,021892299 | 1 |
| ZNF415 | -1,056847483 | 0,007692159 | 1 |
| KCNG1 | -1,096160008 | 0,017440004 | 1 |
| CD200 | -1,192987911 | 0,009404661 | 1 |
| FOSB | -1,217341649 | 0,025144828 | 1 |
| IL8 | -1,349314992 | 0,014854739 | 1 |
| HBA2 | -1,571906696 | 0,008096242 | 1 |
| SLC4A1 | -1,60444022 | 0,031414864 | 1 |
| HBB | -1,672919008 | 0,008461082 | 1 |

**Supplementary Table 7 - Table with the differentially expressed genes between pre-BCG1 and pre-BCG7.**
This table shows the differentially expressed genes at punadj<0.05 between pre-BCG1 and pre-BCG7.
Candidate trained immunity genes are highlighted in the final column.

| Gene name | log2 Fold Change | p value | adjusted p value | Candidate trained immunity gene |
|---|---|---|---|---|
| RYR1 | 2,941 | 0,000117803 | 0,072446462 | |
| ACE | 0,874 | 0,000140314 | 0,082971722 | |
| ITGA7 | 1,190 | 0,000215953 | 0,120733696 | |
| HBA1 | -1,855 | 0,000272809 | 0,136969217 | |
| TCN1 | -1,052 | 0,000326889 | 0,155547135 | |
| HBA2 | -2,062 | 0,000513276 | 0,225467342 | |
| CTSG | -1,383 | 0,000660736 | 0,278314732 | |
| DEFA3 | -1,496 | 0,000748616 | 0,311070294 | |
| HBB | -2,096 | 0,000974312 | 0,394199025 | |
| PGLYRP1 | -1,421 | 0,001210993 | 0,477395128 | |
| CEACAM6 | -1,532 | 0,001203513 | 0,477395128 | |
| ELANE | -1,041 | 0,001850065 | 0,693751727 | |
| CFL2 | -0,701 | 0,002652792 | 0,971079608 | |
| AREGB | 2,247 | 0,032443113 | 1 | |
| KCNMA1 | 1,272 | 0,002974942 | 1 | |
| MEF2BNB-MEF2B | 1,269 | 0,02137519 | 1 | |
| SAP25 | 1,149 | 0,005258327 | 1 | |
| OVCA2 | 0,965 | 0,046970215 | 1 | |
| SAPCD1 | 0,946 | 0,033293913 | 1 | |
| NME2 | 0,750 | 0,003645995 | 1 | |
| MUC1 | 0,710 | 0,01302624 | 1 | |
| FAM20A | 0,708 | 0,024165881 | 1 | |
| FPR2 | 0,614 | 0,023707981 | 1 | |
| MAMDC4 | 0,592 | 0,002876706 | 1 | |
| HES4 | 0,585 | 0,033228181 | 1 | |
| TWF1 | 0,582 | 0,027081585 | 1 | |
| SIGLEC11 | 0,576 | 0,049585346 | 1 | |
| GBP1 | 0,569 | 0,008447504 | 1 | Yes |
| RUFY4 | 0,566 | 0,008329557 | 1 | |
| TGM2 | 0,553 | 0,008460209 | 1 | |
| SLAMF8 | 0,545 | 0,016438939 | 1 | |
| CCDC153 | 0,541 | 0,048089647 | 1 | |
| ADORA3 | 0,523 | 0,040045555 | 1 | |
| EFEMP2 | 0,522 | 0,018114543 | 1 | |
| ODF3B | 0,522 | 0,012071538 | 1 | |
| AP000295,9 | 0,517 | 0,005486546 | 1 | |
| C1QA | 0,501 | 0,034607742 | 1 | |
| SAP30 | 0,492 | 0,048799636 | 1 | |
| S100A13 | 0,487 | 0,022434082 | 1 | |
| AIM2 | 0,483 | 0,042747173 | 1 | Yes |

| | | | | |
|---|---|---|---|---|
| SLC26A1 | 0,469 | 0,020017057 | 1 | |
| CASP5 | 0,461 | 0,047213855 | 1 | Yes |
| MSR1 | 0,453 | 0,03842017 | 1 | |
| DDO | 0,445 | 0,024154433 | 1 | |
| SAMD4A | 0,432 | 0,046899458 | 1 | |
| TAGLN | 0,428 | 0,009172894 | 1 | |
| EPSTI1 | 0,426 | 0,040747824 | 1 | |
| RELL2 | 0,425 | 0,01457999 | 1 | |
| SDC3 | 0,425 | 0,038277269 | 1 | |
| HSPA6 | 0,424 | 0,007136701 | 1 | |
| IL15RA | 0,414 | 0,025549244 | 1 | |
| CTSL1 | 0,413 | 0,049300247 | 1 | |
| NAT1 | 0,410 | 0,033721274 | 1 | |
| XAF1 | 0,405 | 0,00844103 | 1 | |
| GALNT4 | 0,399 | 0,033197071 | 1 | |
| P2RY6 | 0,393 | 0,041702275 | 1 | |
| ADAMTSL4 | 0,393 | 0,019912378 | 1 | |
| MYOF | 0,391 | 0,038590773 | 1 | |
| MAP3K6 | 0,391 | 0,014946439 | 1 | |
| ITPKC | 0,391 | 0,006776491 | 1 | |
| TRIM69 | 0,378 | 0,026317866 | 1 | |
| PLB1 | 0,378 | 0,015327207 | 1 | |
| ECHDC3 | 0,378 | 0,046631846 | 1 | |
| ACSL5 | 0,373 | 0,041736198 | 1 | |
| FGFBP2 | -0,383 | 0,036547248 | 1 | |
| MXRA7 | -0,384 | 0,035192129 | 1 | |
| MSRB3 | -0,387 | 0,049149542 | 1 | |
| TIMM23B | -0,390 | 0,035140792 | 1 | |
| TGFBR3 | -0,401 | 0,045763291 | 1 | |
| CDCA7L | -0,411 | 0,041833477 | 1 | |
| CNR2 | -0,416 | 0,041451689 | 1 | |
| IL2RB | -0,419 | 0,021796056 | 1 | |
| C20orf94 | -0,421 | 0,044428167 | 1 | |
| SLAIN1 | -0,424 | 0,023593214 | 1 | |
| NEDD4L | -0,426 | 0,018960637 | 1 | |
| GNG7 | -0,434 | 0,005636585 | 1 | |
| KLHL11 | -0,437 | 0,041508149 | 1 | |
| RALGPS2 | -0,437 | 0,014769466 | 1 | |
| MLLT4 | -0,456 | 0,011866502 | 1 | |
| AKAP12 | -0,462 | 0,042275449 | 1 | |
| SPIN4 | -0,463 | 0,019648875 | 1 | |
| PEAK1 | -0,467 | 0,025074986 | 1 | |
| NCAM1 | -0,474 | 0,033383214 | 1 | |
| DLG3 | -0,476 | 0,014970698 | 1 | |
| IMMP1L | -0,481 | 0,032162565 | 1 | |

| | | | |
|---|---|---|---|
| OSBPL10 | -0,486 | 0,041137124 | 1 |
| ODZ1 | -0,490 | 0,009195945 | 1 |
| RGPD8 | -0,490 | 0,041652867 | 1 |
| ZNF530 | -0,500 | 0,044470425 | 1 |
| C1orf150 | -0,511 | 0,027267936 | 1 |
| COBLL1 | -0,519 | 0,011564114 | 1 |
| FCGBP | -0,521 | 0,049809413 | 1 |
| CHCHD6 | -0,523 | 0,011609497 | 1 |
| GAMT | -0,526 | 0,024019948 | 1 |
| CPA3 | -0,538 | 0,019933671 | 1 |
| CTD-2370N5,3 | -0,548 | 0,007710109 | 1 |
| STAP1 | -0,565 | 0,039062657 | 1 |
| PKIG | -0,566 | 0,015368835 | 1 |
| MKL2 | -0,578 | 0,023032849 | 1 |
| FAM46C | -0,581 | 0,034520654 | 1 |
| CEBPE | -0,582 | 0,037397591 | 1 |
| MS4A2 | -0,591 | 0,021189027 | 1 |
| BOLA2B | -0,593 | 0,04410199 | 1 |
| CCDC144A | -0,601 | 0,00472668 | 1 |
| HRH4 | -0,602 | 0,030961183 | 1 |
| KIAA0125 | -0,629 | 0,029173614 | 1 |
| B3GNT7 | -0,650 | 0,032271663 | 1 |
| MPO | -0,665 | 0,004482082 | 1 |
| BOK | -0,669 | 0,022048785 | 1 |
| FCRL2 | -0,679 | 0,015633107 | 1 |
| CD79A | -0,683 | 0,026484315 | 1 |
| KLRC4 | -0,687 | 0,036115179 | 1 |
| FCRLA | -0,695 | 0,025595649 | 1 |
| CXCR5 | -0,704 | 0,041310191 | 1 |
| CYP4F3 | -0,704 | 0,02310979 | 1 |
| AKAP5 | -0,707 | 0,035020594 | 1 |
| FCRL1 | -0,722 | 0,026794196 | 1 |
| PAWR | -0,724 | 0,003336898 | 1 |
| PHLDB2 | -0,740 | 0,004978222 | 1 |
| MS4A3 | -0,749 | 0,003834872 | 1 |
| ZNF682 | -0,794 | 0,017502381 | 1 |
| BPI | -0,848 | 0,032467668 | 1 |
| PAX5 | -0,861 | 0,026631795 | 1 |
| OLR1 | -0,880 | 0,045409412 | 1 |
| AZU1 | -0,932 | 0,011789709 | 1 |
| MMP9 | -0,933 | 0,007418439 | 1 |
| ANXA3 | -0,955 | 0,013815894 | 1 |
| RNASE3 | -1,025 | 0,010708942 | 1 |
| ARG1 | -1,029 | 0,031452281 | 1 |
| PRTN3 | -1,061 | 0,003816369 | 1 |

| | | | |
|---|---|---|---|
| MMP8 | -1,184 | 0,026781144 | 1 |
| MYH3 | -1,202 | 0,008838566 | 1 |
| CHI3L1 | -1,243 | 0,019414398 | 1 |
| SLPI | -1,246 | 0,003432196 | 1 |
| CA1 | -1,275 | 0,005140457 | 1 |
| DEFA4 | -1,301 | 0,014181587 | 1 |
| LCN2 | -1,335 | 0,005350238 | 1 |
| OLFM4 | -1,365 | 0,010038908 | 1 |
| CAMP | -1,457 | 0,003688937 | 1 |
| LTF | -1,494 | 0,00594477 | 1 |
| CRISP3 | -1,501 | 0,00300253 | 1 |
| DEFA1 | -1,563 | 0,017337127 | 1 |
| CEACAM8 | -1,593 | 0,003724853 | 1 |
| ALAS2 | -1,668 | 0,010983346 | 1 |
| SLC4A1 | -1,990 | 0,007938291 | 1 |
| HBG2 | -2,243 | 0,004283171 | 1 |

**Supplementary Table 8 – Maintained upregulated and downregulated genes.**

This table shows the differentially expressed genes which were upregulated at BCG6 and remain upregulated at pre-BCG7, as well as the downregulated genes at BCG6 which remain downregulated at pre-BCG7 (all at punadj<0.05). Candidate trained immunity genes are indicated.

| **Genes that remain upregulated between BCG6 and Pre-BCG7** | **Candidate trained immunity gene** |
|---|---|
| ACE | |
| AIM2 | Yes |
| C1QA | |
| CASP5 | Yes |
| DDO | |
| ECHDC3 | |
| EPSTI1 | |
| FAM20A | |
| GBP1 | Yes |
| HSPA6 | |
| IL15RA | |
| ITGA7 | |
| KCNMA1 | |
| MEF2BNB-MEF2B | |
| MUC1 | |
| NAT1 | |
| NME2 | |
| ODF3B | |
| RUFY4 | |
| S100A13 | |
| SAMD4A | |
| SDC3 | |
| SLAMF8 | |
| SLC26A1 | |
| TGM2 | |
| TRIM69 | |
| **Genes that remain downregulated between BCG6 and Pre-BCG7** | **Candidate trained immunity gene** |
| AKAP5 | |
| B3GNT7 | |
| BOK | |
| CA1 | |
| CCDC144A | |
| CD79A | |
| CXCR5 | |
| DLG3 | |
| FCGBP | |
| FCRL1 | |
| FCRL2 | |
| FCRLA | |

| |
|---|
| FGFBP2 |
| HBA1 |
| HBA2 |
| HBB |
| IL2RB |
| KLRC4 |
| ODZ1 |
| OSBPL10 |
| PAWR |
| PAX5 |
| PKIG |
| RALGPS2 |
| SLAIN1 |
| SLC4A1 |
| TGFBR3 |

**Supplementary Table 9 - Characteristics of NMIBC patients from the BlaZIB and UroLife studies that were part of the study into BCG exposure and respiratory infections.**

| | BCG-exposed | Partially BCG-exposed | BCG-unexposed |
|---|---|---|---|
| Total n | 109 | 298 | 250 |
| Female sex (%) | 15 (13.8) | 49 (16.4) | 52 (20.8) |
| Age at questionnaire (median (range)) | 71 (42-85) | 70 (44-90) | 70 (27-92) |
| Flu vaccination in previous 2 years (%) | 72 (66.1) | 189 (63.4) | 164 (65.6) |
| BCG vaccination ever (%) | 16 (14.7) | 56 (18.8) | 50 (20.0) |
| History of chronic lung disease (asthma, COPD, bronchitis, emphysema) | 16 (14.7) | 42 (14.1) | 38 (15.2) |
| Never smoker (%) | 17 (15.6) | 59 (19.8) | 55 (22.0) |
| Previous smoker (%) | 79 (72.5) | 216 (72.5) | 169 (67.6) |
| Current smoker (%) | 13 (11.9) | 23 (7.7) | 26 (10.4) |

**Supplementary Table 10 - Results of the multivariable association analysis for BCG exposure and respiratory infections among NMIBC patients in the BlaZIB and UroLife studies.**

| | **NMIBC BCG** | | | **NMIBC no BCG** | **Adjusted* ORs (95% CI) NMIBC patients** | | |
|---|---|---|---|---|---|---|---|
| | Total | Exposed | Partially exposed | | Exposed versus no BCG | Partially exposed versus no BCG | BCG versus no BCG |
| Total | 407 | 109 | 298 | 250 | - | - | - |
| Pneumonia (%) | 23 (5.7) | 4 (3.7) | 19 (6.4) | 22 (8.8) | 0.39 (0.13-1.18) | 0.68 (0.36-1.32) | 0.60 (0.32-1.12) |
| Bronchitis (%) | 11 (2.7) | 5 (4.6) | 6 (2.0) | 10 (4.0) | 1.23 (0.40-3.80) | 0.53 (0.19-1.51) | 0.72 (0.29-1.75) |
| Laryngitis (%) | 20 (4.9) | 6 (5.5) | 14 (4.7) | 16 (6.4) | 0.93 (0.35-2.51) | 0.76 (0.36-1.62) | 0.80 (0.40-1.61) |
| Flu (%) | 72 (17.7) | 19 (17.4) | 53 (17.8) | 50 (20.0) | 0.89 (0.49-1.61) | 0.87 (0.56-1.34) | 0.87 (0.58-1.31) |
| Cold (%) | 223 (54.8) | 53 (48.6) | 170 (57.0) | 155 (62.0) | 0.60 (0.38-0.95) | 0.85 (0.60-1.20) | 0.77 (0.56-1.07) |
| All above RTI (%) | 254 (62.4) | 63 (57.8) | 191 (64.1) | 171 (68.4) | 0.65 (0.40-1.04) | 0.85 (0.59-1.22) | 0.79 (0.56-1.11) |

*Adjusted for age (continuous variable), sex (female versus male), smoking (3 categories), chronic lung disease (yes vs no), flu vaccination in 2018/2019 (yes vs no).

**Supplementary Table 11 - Characteristics of the Nijmegen Bladder Cancer Study population that received at least 5 BCG instillations and was included in the exome chip analyses for NMIBC recurrence and progression.**

| | | **n = 215 (%)** |
|---|---|---|
| **Gender** | Male | 189 (88.1) |
| | Female | 26 (11.9) |
| **Median age at diagnosis (range; in years)** | | 65 (39 – 83) |
| **Smoking status (cigarettes) at time of diagnosis** | Never smoker | 34 (15.8) |
| | Former smoker | 109 (50.7) |
| | Current smoker | 40 (18.6) |
| | Missing | 32 (14.9) |
| **Primary tumor stage[1]** | | |
| | Ta | 16 (7.4) |
| | Ta+CIS | 17 (7.9) |
| | T1 | 83 (38.6) |
| | T1+CIS | 46 (21.4) |
| | CIS | 49 (22.8) |
| | Unknown | 4 (1.9) |
| **Primary tumor grade[1]** | Low grade | 13 (6.0) |
| | High grade | 200 (93.0) |
| | Unknown | 2 (0.9) |
| **BCG instillations** | iBCG (≥5 instillations) | 215 (100.0) |
| | iBCG + mBCG (≥7 instillations) | 137 (63.7) |
| **Recurrence[2]** | | 79 (36.7) |
| **Median time at risk for recurrence[2] (IQR; in years)** | | 2.8 (1.2 – 5.0) |
| **Progression[2]** | | 47 (21.9) |
| **Median time at risk for progression[2] (IQR; in years)** | | 4.0 (2.1 – 5.0) |
| **Progression to MIBC[2]** | | 17 (7.9) |
| **Median time at risk for progression to MIBC[2] (IQR; in years)** | | 4.5 (2.4 – 5.0) |
| **Median follow-up time (IQR; in years)** | | 4.7 (2.9 – 8.5) |

CIS = Carcinoma in situ; BCG = Bacillus Calmette-Guérin; iBCG = induction therapy of BCG; mBCG = maintenance therapy of BCG; IQR = interquartile range; 1 = highest tumor stage/grade based in transurethral resection of the (bladder) tumor (TURT) and reTURT combined. 2 = restricted to the first five years after diagnosis.

**Supplementary Table 12 - Results of the association analysis of low and rare-frequency variants and clinical outcome in patients of the Nijmegen Bladder Cancer Study (NBCS).**

The clinical outcomes which were assessed were recurrence free survival and progression free survival.

Each table shows the candidate trained immunity genes and their association with RFS or PFS after 5 or more and 7 or more BCG instillations.

RFS=recurrence free survival. PFS=progression free survival.

**Supplementary Table 12.1 - RFS and ≥5 BCG instillations.**
**Displays the genes that associate with recurrence free survival after 5 or more BCG instillations.**

| gene_name | p.value | Q | n.marker.test | n.indiv | df | npatients | nsnvs_incl_monom |
|---|---|---|---|---|---|---|---|
| AIM2 | 0,6668 | -24,75979723 | 3 | 215 | 1,057379136 | 215 | 3 |
| ATG16L1 | 0,1454 | 132,0898301 | 3 | 215 | 1,038824298 | 215 | 4 |
| ATG16L2 | 1 | -1,086464384 | 1 | 215 | 0,984934949 | 215 | 2 |
| ATG2B | **0,0552** | 97,7492291 | 9 | 215 | 1,431773137 | 215 | 19 |
| ATG7 | **0,0518** | 103,4755519 | 3 | 215 | 1,360649517 | 215 | 6 |
| CASP1 | 0,9941 | -63,77052844 | 17 | 215 | 3,132342687 | 215 | 25 |
| CASP5 | 0,3916 | -4,601733516 | 5 | 215 | 2,388367352 | 215 | 10 |
| EHMT2 | 0,92 | -36,51371431 | 4 | 215 | 1,388883661 | 215 | 5 |
| GBP1 | 0,9872 | -29,65079869 | 5 | 215 | 1,858544486 | 215 | 9 |
| GBP2 | 0,7278 | -29,06109049 | 3 | 215 | 1,238003061 | 215 | 3 |
| GBP3 | 0,7812 | -80,66169322 | 4 | 215 | 1,450503533 | 215 | 5 |
| GBP4 | 0,6805 | -92,65114213 | 3 | 215 | 1,024674623 | 215 | 9 |
| GBP5 | 0,2625 | 0,703323814 | 2 | 215 | 1,729617129 | 215 | 5 |
| HK2 | 0,6194 | -31,79680752 | 7 | 215 | 2,115140923 | 215 | 11 |
| HNF1A | 0,1993 | 155,1363473 | 9 | 215 | 1,52379629 | 215 | 11 |
| HNF1B | 0,3863 | -12,36817055 | 3 | 215 | 1,230580783 | 215 | 3 |
| IL18 | 0,3705 | -0,651576851 | 7 | 215 | 2,593914192 | 215 | 8 |
| IL1RN | 0,5804 | -23,88814133 | 2 | 215 | 1,014198219 | 215 | 3 |
| IL32 | 1 | -0,21718186 | 1 | 215 | 0,990179793 | 215 | 3 |
| IL6 | 0,9394 | -197,9905753 | 17 | 215 | 2,56983184 | 215 | 26 |
| KDM4A | 1 | -27,82896042 | 2 | 215 | 1,095446347 | 215 | 6 |
| KDM4B | 0,341 | -0,04469697 | 1 | 215 | 0,966828439 | 215 | 4 |
| KDM4C | 0,3043 | 41,91837758 | 9 | 215 | 5,674905477 | 215 | 11 |
| KDM4D | 0,753 | -24,48704195 | 7 | 215 | 1,826573384 | 215 | 8 |
| NLRP3 | 0,8646 | -57,36367658 | 3 | 215 | 1,197414773 | 215 | 5 |
| NOD2 | 0,4167 | -17,53755037 | 14 | 215 | 2,245635429 | 215 | 23 |
| PFKP | 0,2341 | 11,30863118 | 5 | 215 | 1,319013323 | 215 | 7 |
| RIPK2 | 0,5976 | -6,938540745 | 1 | 215 | 1,015876256 | 215 | 2 |
| TNF | 0,336 | 126,3212538 | 127 | 215 | 28,81735127 | 215 | 202 |

**Supplementary Table 12.2 - RFS and ≥7 BCG instillations.**
**Displays the genes that associate with recurrence free survival after 7 or more BCG instillations.**

| gene_name | p.value | Q | n.marker.test | n.indiv | df | npatients | nsnvs_incl_monom |
|---|---|---|---|---|---|---|---|
| AIM2 | 0,0781 | 33,65764805 | 2 | 137 | 1,100489021 | 137 | 3 |
| ATG16L1 | **0,058** | 151,9812802 | 3 | 137 | 0,944930974 | 137 | 4 |
| ATG16L2 | 1 | -1,181467244 | 1 | 137 | 1,000094082 | 137 | 2 |
| ATG2B | 0,1165 | 26,59945526 | 8 | 137 | 1,508927714 | 137 | 19 |
| ATG7 | **0,0521** | 59,55106554 | 2 | 137 | 1,19111002 | 137 | 6 |
| CASP1 | 0,6011 | -13,23106529 | 15 | 137 | 2,993560518 | 137 | 25 |
| CASP5 | 0,5268 | -13,73028304 | 5 | 137 | 2,533260981 | 137 | 10 |
| EHMT2 | 0,6743 | -14,17080557 | 4 | 137 | 1,254431329 | 137 | 5 |
| GBP1 | 0,1929 | 8,401708316 | 5 | 137 | 1,867480934 | 137 | 9 |
| GBP2 | 0,2868 | 2,605382105 | 3 | 137 | 1,558320155 | 137 | 3 |
| GBP3 | 0,614 | -28,43534132 | 4 | 137 | 1,444531064 | 137 | 5 |
| GBP4 | 0,1745 | 39,0865001 | 3 | 137 | 0,97726995 | 137 | 9 |
| GBP5 | 0,476 | -0,514678761 | 2 | 137 | 1,86123812 | 137 | 5 |
| HK2 | 0,8324 | -23,86979026 | 6 | 137 | 2,10002091 | 137 | 11 |
| HNF1A | 0,6898 | -96,21051718 | 9 | 137 | 1,49764708 | 137 | 11 |
| HNF1B | **0,0427** | 96,88458053 | 3 | 137 | 1,108303292 | 137 | 3 |
| IL18 | 0,4928 | -11,71938247 | 7 | 137 | 2,770039732 | 137 | 8 |
| IL1RN | 0,6515 | -13,39071923 | 2 | 137 | 1,018522656 | 137 | 3 |
| IL32 | 1 | -0,160342383 | 1 | 137 | 0,999331758 | 137 | 3 |
| IL6 | 0,3123 | 11,37240346 | 15 | 137 | 2,242202684 | 137 | 26 |
| KDM4A | 0,2429 | 3,661767252 | 2 | 137 | 1,07127914 | 137 | 6 |
| KDM4C | 0,5878 | -27,97011101 | 9 | 137 | 5,715745233 | 137 | 11 |
| KDM4D | 0,7464 | -11,92794231 | 6 | 137 | 1,874059787 | 137 | 8 |
| NLRP3 | 1 | -35,06664189 | 2 | 137 | 1,20817639 | 137 | 5 |
| NOD2 | 0,6083 | -20,31787809 | 13 | 137 | 2,166036649 | 137 | 23 |
| PFKP | 0,3695 | -2,111171642 | 4 | 137 | 1,210588537 | 137 | 7 |
| RIPK2 | 0,8329 | -4,877370114 | 1 | 137 | 0,998522945 | 137 | 2 |
| TNF | 0,1967 | 181,7768161 | 118 | 137 | 24,36444595 | 137 | 202 |

**Supplementary Table 12.3 - PFS and ≥5 BCG instillations.**
**Displays the genes that associate with progression free survival after 5 or more BCG instillations.**

| gene_name | p.value | Q | n.marker.test | n.indiv | df | npatients | nsnvs_incl_monom |
|---|---|---|---|---|---|---|---|
| AIM2 | 0,5732 | -12,6009947 | 3 | 215 | 1,035972928 | 215 | 3 |
| ATG16L1 | 0,2317 | 28,9698611 | 3 | 215 | 1,027726132 | 215 | 4 |
| ATG16L2 | 0,0899 | 3,479819289 | 1 | 215 | 1,025741716 | 215 | 2 |
| ATG2B | **0,008** | 153,4451019 | 9 | 215 | 1,428609768 | 215 | 19 |
| ATG7 | 0,4691 | -11,10950223 | 3 | 215 | 1,305199874 | 215 | 6 |
| CASP1 | 0,7624 | -26,44204776 | 17 | 215 | 2,932521748 | 215 | 25 |
| CASP5 | 0,7842 | -33,1395669 | 5 | 215 | 2,377722736 | 215 | 10 |
| EHMT2 | **0,0053** | 130,6742915 | 4 | 215 | 1,482973027 | 215 | 5 |
| GBP1 | 0,8392 | -13,65944477 | 5 | 215 | 1,995389007 | 215 | 9 |

| | | | | | | | |
|---|---|---|---|---|---|---|---|
| GBP2 | 0,4482 | -8,651714394 | 3 | 215 | 1,346006045 | 215 | 3 |
| GBP3 | 0,8938 | -53,75550237 | 4 | 215 | 1,387036502 | 215 | 5 |
| GBP4 | 0,6771 | -57,22545556 | 3 | 215 | 0,971590925 | 215 | 9 |
| GBP5 | 0,5051 | -0,576733035 | 2 | 215 | 2,01332749 | 215 | 5 |
| HK2 | 0,8156 | -26,83707069 | 7 | 215 | 2,211541382 | 215 | 11 |
| HNF1A | 0,1557 | 154,5863827 | 9 | 215 | 1,578299375 | 215 | 11 |
| HNF1B | 0,4879 | -13,10416181 | 3 | 215 | 1,349908748 | 215 | 3 |
| IL18 | 0,2266 | 30,61977526 | 7 | 215 | 2,4513315 | 215 | 8 |
| IL1RN | 0,5132 | -12,09519724 | 2 | 215 | 1,007439781 | 215 | 3 |
| IL32 | 1 | -0,132201666 | 1 | 215 | 0,988142959 | 215 | 3 |
| IL6 | 0,2526 | 40,66687734 | 17 | 215 | 2,388292864 | 215 | 26 |
| KDM4A | 1 | -16,29219747 | 2 | 215 | 1,050785424 | 215 | 6 |
| KDM4B | 0,2735 | 0,190147994 | 1 | 215 | 0,986347146 | 215 | 4 |
| KDM4C | 0,2852 | 29,85031216 | 9 | 215 | 5,251538918 | 215 | 11 |
| KDM4D | 0,378 | -3,044346329 | 7 | 215 | 1,751831719 | 215 | 8 |
| NLRP3 | 0,3568 | -4,713791167 | 3 | 215 | 1,121792775 | 215 | 5 |
| NOD2 | 0,8412 | -42,98981127 | 14 | 215 | 2,211179458 | 215 | 23 |
| PFKP | 0,6274 | -10,74069042 | 5 | 215 | 1,47312329 | 215 | 7 |
| RIPK2 | 1 | -5,976955764 | 1 | 215 | 0,996566772 | 215 | 2 |
| TNF | 0,2839 | 125,9999296 | 127 | 215 | 23,89939862 | 215 | 202 |

**Supplementary Table 12.4 - PFS and ≥7 BCG instillations.**
**Displays the genes that associate with progression free survival after 7 or more BCG instillations.**

| gene_name | p.value | Q | n.marker.test | n.indiv | df | npatients | nsnvs_incl_monom |
|---|---|---|---|---|---|---|---|
| AIM2 | 0,1546 | 9,063543939 | 2 | 137 | 1,009003898 | 137 | 3 |
| ATG16L1 | 0,327 | -0,571700501 | 3 | 137 | 1,064808361 | 137 | 4 |
| ATG16L2 | 0,0715 | 4,704832233 | 1 | 137 | 0,970200993 | 137 | 2 |
| ATG2B | **0,0325** | 44,60020007 | 8 | 137 | 1,33311765 | 137 | 19 |
| ATG7 | 0,3489 | -1,559441784 | 2 | 137 | 1,261291555 | 137 | 6 |
| CASP1 | 0,6829 | -10,96007196 | 15 | 137 | 2,771220887 | 137 | 25 |
| CASP5 | 0,7527 | -15,41816113 | 5 | 137 | 2,438596936 | 137 | 10 |
| EHMT2 | **5,00E-04** | 110,0870458 | 4 | 137 | 1,208050539 | 137 | 5 |
| GBP1 | 0,4174 | -1,58329662 | 5 | 137 | 2,029893373 | 137 | 9 |
| GBP2 | 0,2807 | 2,059582358 | 3 | 137 | 1,651557473 | 137 | 3 |
| GBP3 | 0,908 | -26,42021847 | 4 | 137 | 1,473808013 | 137 | 5 |
| GBP4 | 0,4963 | -15,87192298 | 3 | 137 | 1,007245417 | 137 | 9 |
| GBP5 | 0,371 | -0,045487272 | 2 | 137 | 1,909587288 | 137 | 5 |
| HK2 | 0,5669 | -6,933682488 | 6 | 137 | 2,506234496 | 137 | 11 |
| HNF1A | 0,7746 | -79,25404939 | 9 | 137 | 1,414903873 | 137 | 11 |
| HNF1B | 0,6077 | -9,348132027 | 3 | 137 | 1,14339123 | 137 | 3 |
| IL18 | 0,3446 | 1,579622961 | 7 | 137 | 2,446238991 | 137 | 8 |
| IL1RN | 0,5007 | -5,833158786 | 2 | 137 | 0,981415478 | 137 | 3 |
| IL32 | 1 | -0,076845213 | 1 | 137 | 0,991893191 | 137 | 3 |
| IL6 | 0,6251 | -40,22948081 | 15 | 137 | 2,068295195 | 137 | 26 |
| KDM4A | 1 | -7,581690349 | 2 | 137 | 1,015381476 | 137 | 6 |
| KDM4C | 0,6696 | -26,06678812 | 9 | 137 | 4,870546662 | 137 | 11 |
| KDM4D | 0,462 | -3,311955951 | 6 | 137 | 1,541533778 | 137 | 8 |
| NLRP3 | 1 | -20,25122951 | 2 | 137 | 1,152741114 | 137 | 5 |
| NOD2 | 0,6842 | -16,71527872 | 13 | 137 | 2,406863266 | 137 | 23 |
| PFKP | 0,3 | 0,477739781 | 4 | 137 | 1,238970309 | 137 | 7 |
| RIPK2 | 0,9095 | -3,376995089 | 1 | 137 | 0,981292619 | 137 | 2 |
| TNF | 0,5785 | -50,09080898 | 118 | 137 | 18,97301254 | 137 | 202 |

**Supplementary Table 13 - Primers that were used for ChIP-PCR analysis.**

| Gene | Primer name | Forward/Reverse | Sequence |
|---|---|---|---|
| *IL6* | IL-6 region 1 | Forward | TCGTGCATGACTTCAGCTTT |
| *IL6* | IL-6 region 1 | Reverse | GCGCTAAGAAGCAGAACCAC |
| *IL6* | IL-6 region 2 | Forward | AGGGAGAGCCAGAACACAGA |
| *IL6* | IL-6 region 2 | Reverse | GAGTTTCCTCTGACTCCATCG |
| *TNF* | TNF region 1 | Forward | CAGGCAGGTTCTCTTCCTCT |
| *TNF* | TNF region 1 | Reverse | GCTTTCAGTGCTCATGGTGT |
| *TNF* | TNF region 2 | Forward | AGAGGACCAGCTAAGAGGGA |
| *TNF* | TNF region 2 | Reverse | AGCTTGTCAGGGGATGTGG |
| *IL1B* | IL1B region 1 | Forward | AATCCCAGAGCAGCCTGTTG |
| *IL1B* | IL1B region 1 | Reverse | AACAGCGAGGGAGAAACTGG |
| *IL1B* | IL1B region 2 | Forward | CATGGCTGCTTCAGACACCT |
| *IL1B* | IL1B region 2 | Reverse | ACACATGAACGTAGCCGTCA |
